# Synergy between *cis*-regulatory elements can render cohesin dispensable for distal enhancer function

**DOI:** 10.1101/2024.10.04.615095

**Authors:** Karissa L. Hansen, Annie S. Adachi, Luca Braccioli, Smit Kadvani, Ryan M. Boileau, Bozhena Pokorny, Rini Shah, Erika C. Anderson, Moreno Martinovic, Kaite Zhang, Irié Carel, Kenya Bonitto, Robert Blelloch, Geoffrey Fudenberg, Elzo de Wit, Elphège P. Nora

## Abstract

Enhancers are critical genetic elements controlling transcription from promoters, but the mechanisms by which they convey regulatory information across large genomic distances remain elusive. Here, we engineered pluripotent stem cells in which cohesin loop extrusion can be inducibly disrupted without causing confounding cell cycle defects. While evident, transcriptional dysregulation was cell-type specific, and not all loci with distal enhancers depend equally on cohesin extrusion. Using comparative genome editing, we demonstrate that enhancer-promoter communication across as little as 20 kilobases can rely on cohesin. However, promoter-proximal regulatory elements can support long-range, cohesin-independent enhancer action – either upon disabling extrusion or across strong CTCF insulators. Finally, transcriptional dynamics and the emergence of new embryonic cell types in response to differentiation cues remained largely robust to disrupting cohesin extrusion. Beyond establishing novel experimental strategies to study cohesin functions in enhancer biology, our work provides mechanistic insight accounting for both cell type- and genomic context-specificity.

## Introduction

Distal regulatory elements, such as enhancers, can convey essential regulatory information to target promoters across large genomic distances (*1*). In mammalian genomes, enhancers can lie tens to hundreds of kilobases away from the promoters they regulate (*2*). While current technologies can identify candidate enhancers across the genome (*3*), a key question remains: how do enhancers communicate with target promoters across large genomic distances?

Dynamic folding patterns of the chromatin fiber can bring together enhancers and promoters in the 3D space of the cell nucleus (*4*, *5*). DNA loop extrusion by the cohesin complex is a major contributor to chromosome folding and its dynamics (*6*). There is already strong evidence that cohesin loop extrusion, and the long-range interactions it creates, can in some cases facilitate the action of remote enhancers during development (*7–9*). Indeed, genetic evidence has long implicated cohesin in transcriptional regulation during development (*10*), with early screens for factors mediating long-range enhancer-promoter communication in *Drosophila* incriminating the NIPBL homolog (*11*), a cohesin co-factor later discovered to be essential for DNA loop extrusion. Furthermore, mutations in NIPBL or cohesin can cause multisystem disorders, such as Cornelia de Lange syndrome, as well as tissue-specific diseases, including several types of cancer (*10*).

However, the extent to which cohesin, and 3D genome folding in general, contribute to long-range enhancer function remains highly debated. Cohesin removal triggers widespread chromosome misfolding (*12*, *13*), but due to the essential role of cohesin in sister chromatid cohesion, studies in cycling cells are constrained to a brief time window of only several hours, during which transcriptional mis-regulation is minimal (*14*). Therefore, the full scope of loci relying on cohesin extrusion, as well the extent of their dependency, remains to be determined.

Loci with distal enhancers are not uniformly dependent on cohesin (*15–17*). Synthetic approaches, employing either a reconstructed reporter locus (*18*) or artificial enhancer activation (*19*), have suggested that cohesin may be especially important for long-range rather than short-range enhancer-promoter communication. Moving beyond these synthetic systems, understanding what makes some loci depend more on cohesin than others would require a tractable model where both loop extrusion and genomic context can be manipulated in concert.

Moreover, the lack of controlled developmental models to inducibly disable cohesin loop extrusion in cycling cells, without introducing confounding cell cycle defects, also leaves pressing questions about the importance of the *cis*-regulatory networks facilitated by cohesin. Using inducible knockouts, it has been proposed that cohesin may be more important for establishing new transcriptional programs than for maintaining them (*20*). Nevertheless, other work reported that cohesin loss may not be a general impediment to transcriptional induction (*21*), and that constitutive genes in various quiescent cell types can still depend on cohesin (*13*, *22*). Complicating the picture, some loci can switch from being reliant upon cohesin to being independent of it, depending on developmental stage (*16*). The contribution of cohesin extrusion to developmental processes therefore remains to be clarified.

Here, we developed a novel genetic approach based on acute NIPBL depletion in pluripotent stem cells, which disrupts long-range cohesin-dependent interactions while still allowing cell cycle progression. We leverage this new system to address long-standing questions, including the identification of loci that rely on cohesin loop extrusion, the genomic basis for their sensitivity, and the relevance of long-range chromosomal interactions in developmental processes such as cell differentiation and embryonic morphogenesis. Our observations establish that enhancer-promoter communication relies on extrusion across genomic distances much shorter than previously appreciated, while also revealing an unexpected buffering mechanism, independent of cohesin, and mediated by promoter-proximal elements.

## Results

### NIPBL depletion can inhibit cohesin loop extrusion without compromising cell division

We created mouse embryonic stem cells (ESCs) where endogenous NIPBL, a cohesin cofactor essential for loop extrusion (Fig. 1A) (*23*), can be acutely depleted with an N-terminal inducible degron (FKBP12^F36V^ system (*24*), hereafter abbreviated FKBP, fig. S1A). In these cells NIPBL levels dropped below 10% within hours of exposure to dTAG-13 (hereafter abbreviated dTAG, fig. S1B), causing drastic loss of long-range chromatin interactions at the scale of Topologically Associating Domains (TADs), reminiscent of cohesin (RAD21)-depleted cells (Fig. 1B and fig. S2). However, NIPBL-depleted cells continued to proliferate normally (Fig. 1C and fig. S1C, H-I), without abnormal nuclear morphology (Fig. 1B) or major cell cycle defects (Fig. S1E-G) – in stark contrast to cohesin-depleted cells. As NIPBL KO ESCs are not viable (Fig. S3), it appears NIPBL-depleted cells can maintain normal cell division because of trace NIPBL leftover.

**Figure 1.**
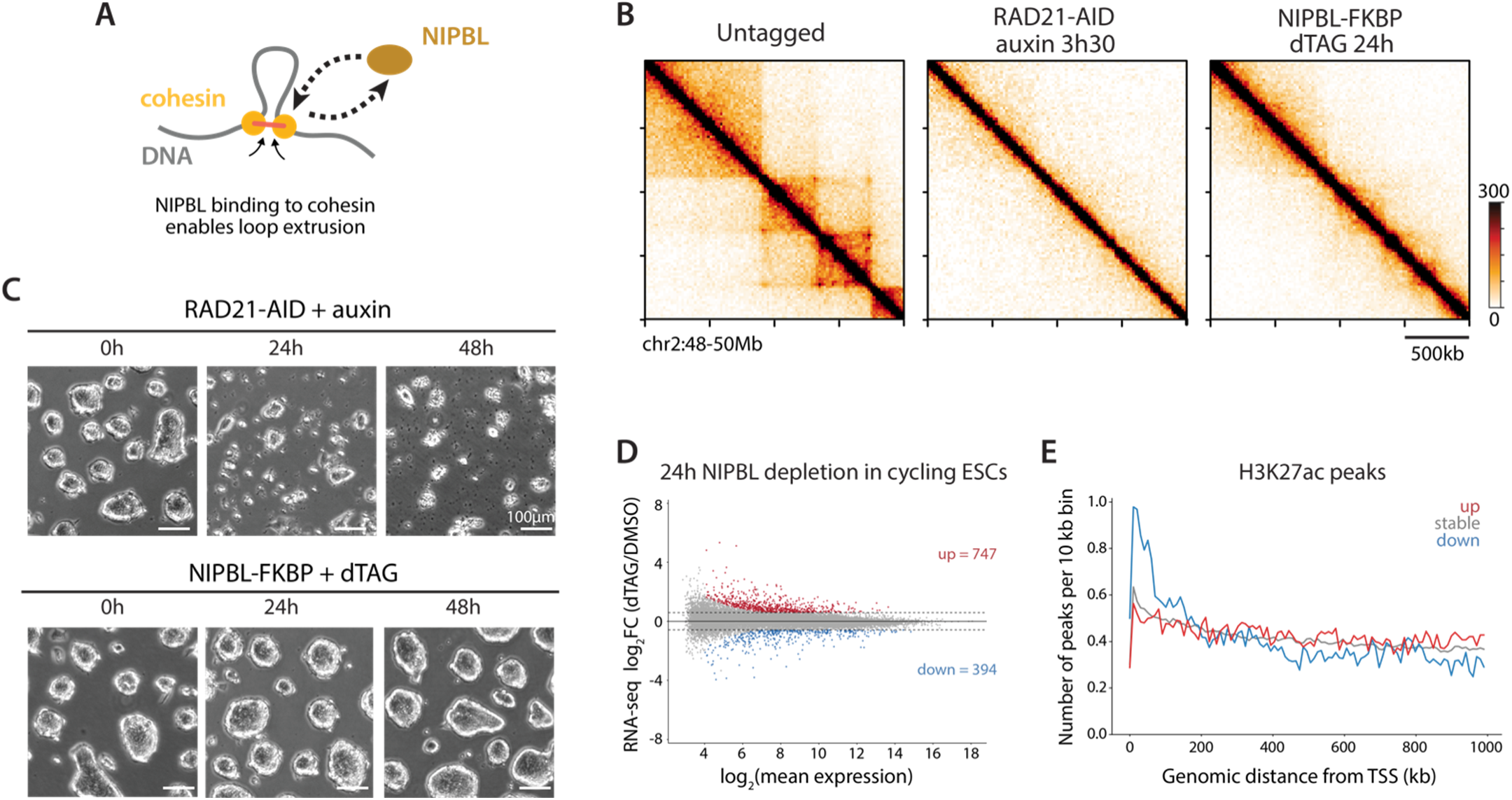
Depleting NIPBL reduces interphase loop extrusion without affecting cell division and leads to transcriptional changes. **(A)** NIPBL binding to cohesin is required for DNA loop extrusion. **(B)** Hi-C contact maps demonstrating that depleting either the core cohesin subunit RAD21 with the Auxin Inducible Degron (AID), or NIPBL with dTAG in FKBP-degron mouse embryonic stem cells (ESCs), both reduce long-range interactions. **(C)** Brightfield microscopy showing that depleting RAD21 rapidly leads to cell death, while depleting NIPBL does not alter viability. **(D)** Depleting NIPBL for 24 hours (h) in ESCs dysregulates hundreds of genes by RNA-seq [False Discovery Rate (FDR) < 0.05 and Fold Change (FC) ≥ 1.5 or ≤ −1.5, n = 3 replicates]. **(E)** The genomic neighborhood of downregulated genes in ESCs has a higher density of active elements, defined by H3K27ac peaks, up to ∼200 kb from the Transcription Start Sites (TSS).

### Loop extrusion is required for transcriptional maintenance at a subset of loci

By mitigating the requirements of cohesin in mitosis, NIPBL-degron cells provide an unprecedented opportunity to dissect the relevance of loop extrusion to transcriptional regulation without confounding cell cycle defects. RNA-seq in NIPBL-FKBP ESCs revealed hundreds of dysregulated genes within 24h of depletion (Fig. 1D). This pattern was reproducible with a time course using an independent cell line generated with the Auxin-Inducible Degron system (NIPBL-AID, fig. S4). Wild-type (WT) untagged cells were largely unaffected by dTAG treatment (fig. S5A). Gene set enrichment analysis (GSEA) underscored the disruption of developmentally relevant pathways without highlighting cell cycle terms, in contrast to cohesin-depleted ESCs (Fig. S5B-E) (*8*).

We then investigated the molecular basis for the selective dysregulation of some genes upon NIPBL depletion. H3K27ac and H3K27me3 CUT&Tag signal remained largely stable even after 24h of NIPBL depletion (fig. S6), indicating the intrinsic regulatory potential of control elements is not altered after short-term disruption of loop extrusion. Instead, we found that downregulated genes tended to have a higher density of H3K27ac and NIPBL peaks within ∼200 kb (Fig. 1E and fig. S7), revealing that they tend to reside in neighborhoods rich in distal regulatory elements, in the genomic range where extrusion enriches physical interactions (fig. S2). Upregulated genes did not exhibit obvious genomic signatures.

Depleting NIPBL from neural progenitor cells (NPCs) also dysregulated hundreds of genes within 24h (Fig. 2A-B and fig. S8), yet the transcriptional response to NIPBL depletion was largely distinct from ESCs (Fig. 2C). Indeed, very few genes were dysregulated in both cell types – which is consistent with the cell-type specific nature of enhancers. Altogether, these analyses indicate that NIPBL, and by extension proper cohesin loop extrusion, are essential for the maintenance of transcriptional profiles, and that the reliance of some genes on loop extrusion is controlled by cell type-specific processes.

**Figure 2.**
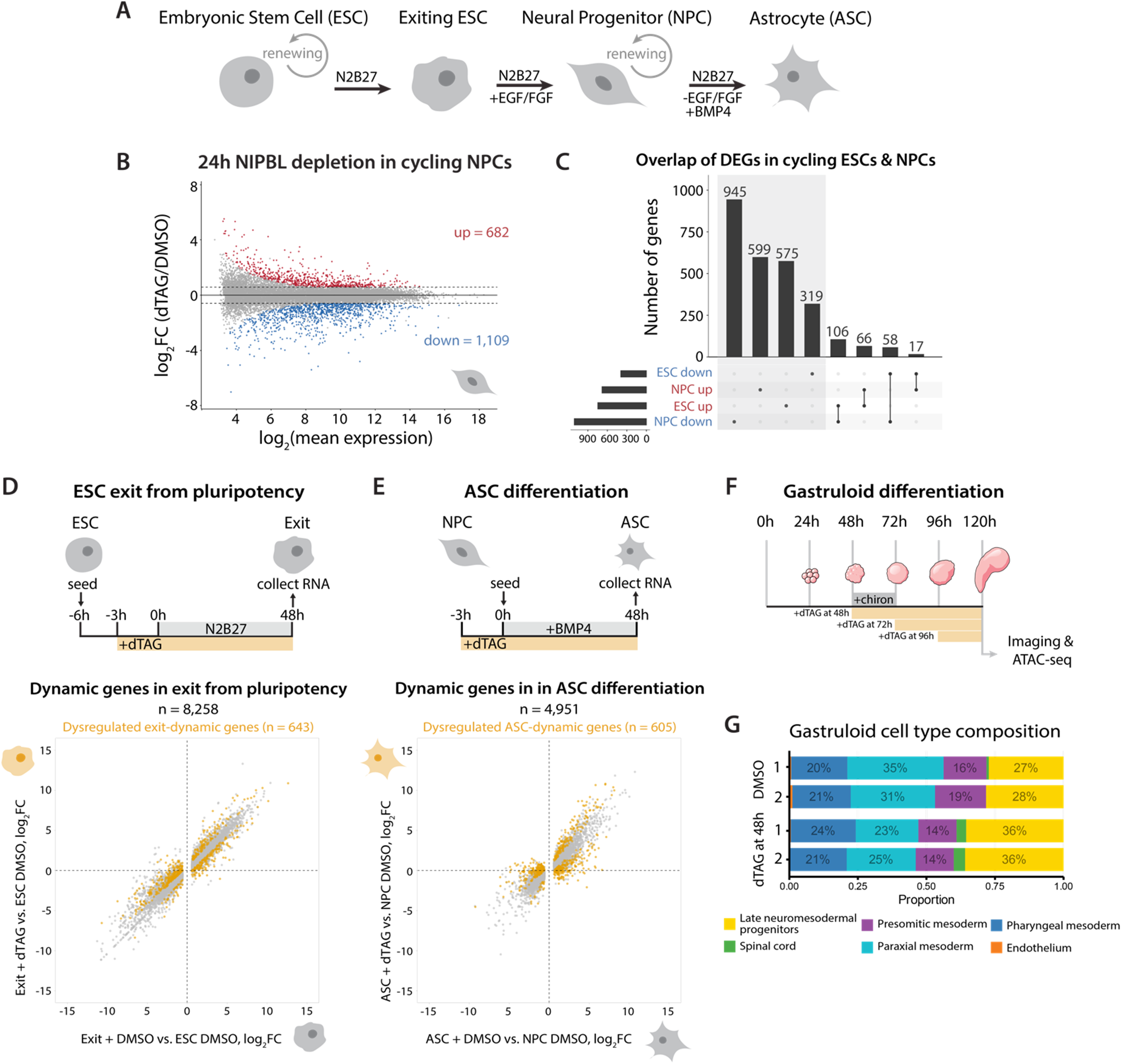
Loop extrusion regulates transcription of distinct gene loci across cell types, yet is largely dispensable for specification of new cell types during differentiation. **(A)** Differentiation scheme from ESCs. **(B)** Hundreds of genes are dysregulated by RNA-seq in NPCs after depleting NIPBL for 24h (FDR < 0.05 and FC ≥ 1.5 or ≤ −1.5, n = 3 replicates). **(C)** Differentially expressed genes (DEGs) across ESCs and NPCs are largely non-overlapping. **(D)** Out of the 8,258 dynamic genes during the exit from pluripotency, NIPBL depletion only alters 643 genes (8%, in goldenrod) specifically during ESC differentiation. Note that despite being called dysregulated, these genes still trend in the expected direction. Fold changes (log_2_) are relative to control (DMSO) ESCs (n = 3 differentiations). **(E)** Similarly, out of the 4,951 dynamic genes during NPC transition to ASCs, NIPBL depletion only alters 605 genes (12%, in goldenrod) specifically during NPC differentiation. Again, these gene largely trend in the expected direction. Fold changes (log_2_) are shown relative to control (DMSO) NPCs (n = 3 differentiations). **(F)** Experimental layout for gastruloid differentiation. **(G)** Cell type composition from bulk ATAC-seq deconvolution in control (DMSO) and NIPBL-depleted gastruloids (dTAG at 48h), showing that all cell types are specified in overall similar proportions (n = 2). See fig. S11 for more timepoints and controls.

### Disrupting loop extrusion does not prevent overall transcriptional dynamics or cell type diversification during differentiation

Cohesin was proposed to play a more essential role in transcriptional induction, for example in response to differentiation cues, rather than for maintaining transcription in steady-state conditions (*20*). However, since acute cohesin degradation prevents cell cycle progression (*25*), the specific role of extrusion in developmental processes that require cell division has remained elusive.

Taking advantage of the proliferative capacity of NIPBL-depleted ESCs, we induced depletion across two developmental transitions. We used RNA-seq assays to read out cellular state and differentiation efficiency, refraining from correlating dysregulated expression with genomic features as the secondary defects arising during these longer depletion times would confound such analyses. First, we assayed the ability of NIPBL-depleted ESCs to exit pluripotency. After 48h in differentiation medium, we observed the expected downregulation of pluripotency markers and upregulation of early differentiation markers (Fig. S9A-C). NIPBL depletion only misregulated 8% of the differentiation-dynamic genes (after subtracting genes already dysregulated in ESC, Fig. 2D and fig. S9D-E), and while expression changes were statistically significant, most of these genes still trended in the expected direction (Fig. 2D). Second, we assayed the ability of NIPBL-depleted NPCs to convert to astrocytes (ASCs). NIPBL depletion did not prevent the transition to an ASC-like transcriptome, with only 12% of differentiation-specific genes being misregulated, but not severely disrupted (Fig. 2E and fig. S10). Thus, the long-range interactions created by cohesin loop extrusion appear largely dispensable for transcriptional dynamics during these cell state transitions, although a small subset of genes can be sensitive. Finally, we address the contribution of DNA loop extrusion to cell fate decisions by using 3D gastruloids (*26*) (Fig. 2F). Estimating cell type composition by deconvolving bulk ATAC-seq signal with a reference scATAC-seq timecourse (*27*, *28*) indicated that all differentiated cell types were present in NIPBL-depleted gastruloids, in nearly normal proportions, even though the morphology of the gastruloids appeared atypical (Fig. 2G and fig. S11).

Altogether, our observations indicate that cohesin loop extrusion is not generally more important for inducing transcriptional changes than for steady-state maintenance. The *cis*-regulatory logic driving transcriptional dynamics and emergence of new cell types therefore appears to operate largely independently of the long-range interactions created by cohesin loop extrusion.

### Cohesin can become essential for enhancer-promoter communication over only tens of kilobases

We next sought to elucidate why some loci rely on loop extrusion for enhancer-promoter communication. Previous studies employing enhancer manipulations – either using synthetic reporters at an engineered locus (*18*), or through artificial enhancer activation (*19*) – indicated that cohesin depletion inhibits distal (>100 kb) but not proximal enhancer-promoter communication. However, it remains unknown how this concept applies to endogenous loci with physiologically active enhancers.

In the absence of a comprehensive list of genetically validated distal enhancers and their target genes, we cannot simply estimate how many *bona fide* enhancer-promoter pairs are sensitive to NIPBL depletion. We therefore focused on select loci for deep functional investigation.

In ESCs, mRNA levels of *Car2* are decreased nearly 70% after depleting NIPBL for 24h and keep decreasing over time (Fig. 3A-B and fig. S12A-B). Depleting RAD21 or CTCF also downregulates *Car2* (Fig. S12C) (*8*, *29*), highlighting its general reliance on the extrusion pathway (Fig. S12C). The simple organization of the *Car2* locus makes it tractable for functional assays: the *Car2* Topologically Associating Domain (TAD) does not harbor any other protein-coding gene but contains two distal H3K27ac clusters, over 100 kb from the transcription start site (TSS) (fig. S12A). Hi-C interactions between the *Car2* TSS and these H3K27ac peaks are reduced upon NIPBL depletion (Fig. S12D-E). CTCF and RAD21 bind within and upstream of the *Car2* promoter, but not elsewhere in the TAD (fig. S12A). To enable single-cell monitoring of *Car2* expression, we tagged one endogenous allele with 2A-2xmClover (GFP) and the other with 2A-tdTomato (RFP), in the NIPBL-FKBP degron cell line (Fig. 3C and fig. S13A). Depleting NIPBL for 24h decreased both GFP and RFP expression nearly 60% after 24h and 80% after 3 days, closely tracking with the transcriptome data (Fig. 3D and fig. S13B-C). This single-cell analysis also revealed that the entire cell population uniformly downregulates *Car2* after depleting NIPBL (Fig. S13A-C).

**Figure 3.**
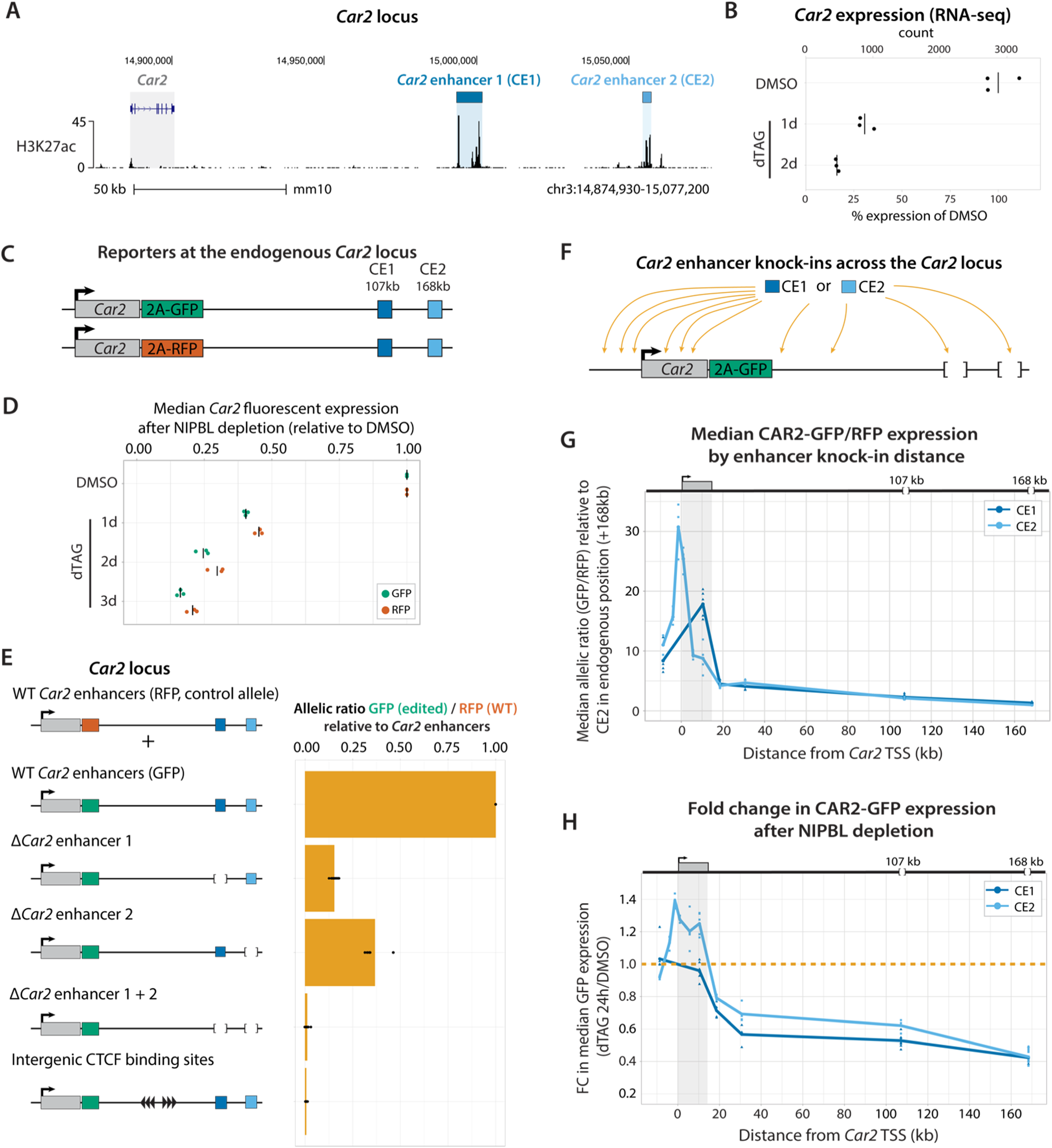
Endogenous locus engineering demonstrates that altering cohesin loop extrusion impairs distal but not proximal enhancer-promoter communication. **(A)** H3K27ac CUT&Tag at the *Car2* locus indicating the locations of putative *Car2* enhancer 1 (CE1) and *Car2* enhancer 2 (CE2). **(B)** *Car2* mRNA expression decreases after 1-2 days of NIPBL depletion in NIPBL-FKBP ESCs. Counts are size factor normalized. Each point is an individual replicate, and bars indicate averages (n = 3). **(C)** *Car2* fluorescent reporter ESCs created by inserting 2A-GFP and 2A-RFP cassettes at the 3’ end of endogenous *Car2.* **(D)** *Car2* fluorescent reporters measured by flow cytometry recapitulate the rapid downregulation observed by RNA-seq. Median fluorescence is shown relative to undepleted control for each allele. Each point is an individual flow replicate and notches indicates averages (n ≥ 3 flows from independent cultures). **(E)** *Car2* reporter expression becomes undetectable after deleting both CE1 and CE2 elements. Inserting an intervening CTCF cassette to locally block cohesin extrusion also abrogates *Car2* expression. Median CAR2-GFP expression (edited allele) is normalized to the WT RFP control allele (allelic ratio). Allelic ratios are shown relative to the cell line with both WT *Car2* enhancers. Each point is an individual flow replicate and bars indicates averages (n ≥ 3 flows). **(F)** Clonal ESCs generated by reinserting either CE1 or CE2 at various locations across an enhancer-less *Car2* allele. **(G)** Promoter-proximal enhancer reinsertion drove higher basal *Car2* expression. The locus map indicates the position of deleted endogenous enhancers CE1 (107 kb) and CE2 (168 kb). Median allelic ratios for enhancer knock-ins across the *Car2* locus are shown relative to the cell line with CE2 in its endogenous location. Each point is an individual flow replicate for the same clonal cell line, and line connects averages (n ≥ 3 flows). **(H)** Fold change in median CAR2-GFP expression after 24h NIPBL depletion, demonstrating that proper loop extrusion is no longer required for *Car2* expression when an enhancer is within 11 kb of the TSS. See fig. S14 for supporting analyses and cell lines (n ≥ 3 flows).

We used this inter-allelic reporter system to precisely quantify the contribution of distal regulatory elements to *Car2* expression. Deleting either of the two H3K27ac clusters on the GFP allele downregulated *Car2* in *cis*, indicating that these act as enhancers; deleting both completely abolished *Car2* expression (Fig. 3E and fig S13E-F). Therefore, both the *Car2* enhancer 1 (CE1) and *Car2* enhancer 2 (CE2) regions are required for *Car2* expression, and act synergistically. Expression of the WT RFP allele is largely unaffected by the enhancer deletions, demonstrating that these enhancers act strictly in *cis*, and that changes in *Car2* expression (which encodes a dispensable carbonate anhydrase) does not trigger confounding secondary *trans* effects (Fig. S13E-F). Disambiguating *cis* versus *trans* effects, by normalizing expression of the edited GFP allele to that of the unedited control RFP allele present in the same cell (*30*), was especially important as we found that differences in cell density between samples can affect basal *Car2* expression (Fig. S13D). Finally, inserting strong CTCF sites between *Car2* and its enhancers completely abolished *Car2* expression in *cis,* indicating that loop extrusion across the locus is absolutely needed to connect *Car2* with its distal enhancers for proper activation (Fig. 3E and fig. S13E-F).

Given recent reports that cohesin may be important for distal, but not proximal, enhancer-promoter communication (*18*, *19*, *22*), we addressed whether the reliance of *Car2* on loop extrusion arises from the large genomic distance between the promoter and its enhancers (107 kb to CE1 and 168 kb to CE2). We generated a collection of clonal ESC lines by reinserting either CE1 or CE2 at various positions across the *Car2* locus, on an allele with both endogenous enhancers deleted, and analyzed at least 2 independent clones for each insertion (Fig. 3F). We observed much stronger *Car2* expression when enhancers are positioned close to the promoter (Fig. 3G and fig. S14A), indicating that longer genomic distances strongly impair enhancer-promoter communication. Of note, we observed similar expression of *Car2* when driven by either CE1 or CE2 from the same position. This indicates that the two enhancers have similar autonomous strengths, and that deleting CE1 has a larger effect on *Car2* expression simply because it is 60 kb closer to the promoter than CE2.

Depleting NIPBL for 24h in these cell lines only downregulated *Car2* when the enhancer was reinserted at least 18 kb from the promoter, with greater fold changes for the most distal positions (Fig. 3H and fig. S14B). In contrast, lines with enhancer insertions within 11 kb of the *Car2* promoter were not downregulated upon NIPBL depletion. Mutating the *Car2* promoter to abolish CTCF binding did not alter the reliance of *Car2* on cohesin loop extrusion (fig. S15). Finally, *Car2* was not significantly downregulated upon depleting NIPBL in NPCs, where the CE1 and CE2 enhancers are not active (fig. S16).

Altogether, our experiments establish that cohesin loop extrusion can be essential to support the communication between distal, but not proximal, enhancer-promoter pairs. Our observations also clarify that reliance on cohesin can arise at much shorter genomic separation than previously appreciated from synthetic loci or artificial enhancer activation systems (*18*, *19*).

### Genomic context elicits distinct behaviors across loci

Intriguingly, our transcriptomic analyses indicate that some loci appear to disobey the simple principle of distance-dependence evident at *Car2*. The *Sox2* locus with its distal super-enhancer in ESCs, the *Sox2* Control Region (SCR) (*31*, *32*), is widely used in the field as a model to understand the mechanisms of long-range enhancer-promoter communication (*30*, *33–35*). Contrasting with our observations at *Car2*, multiple studies indicate that 3D genome folding can be largely decoupled from transcriptional regulation at *Sox2* (*36–38*).

Indeed, we observed striking differences in the behavior of the *Sox2* and *Car2* loci upon inhibiting cohesin loop extrusion. Despite the SCR being over 100 kb away from *Sox2* (Fig. 4A and fig. S17A), NIPBL depletion only downregulated *Sox2* expression by ∼25%, even after 4 days (Fig. 4B and fig. S17B-E), by which time *Car2* expression had dropped by ∼80% (Fig. S12B). This prompted us to investigate why some loci with distal enhancers, such as *Car2*, are more reliant on cohesin extrusion than others, like *Sox2,* and address how these seemingly contrasting behaviors can be reconciled. We therefore tested what accounts for the robustness of the *Sox2*-SCR locus to loss of extrusion: is it the nature of the SCR, the *Sox2* promoter or the presence of additional elements in the genomic context?

**Figure 4.**
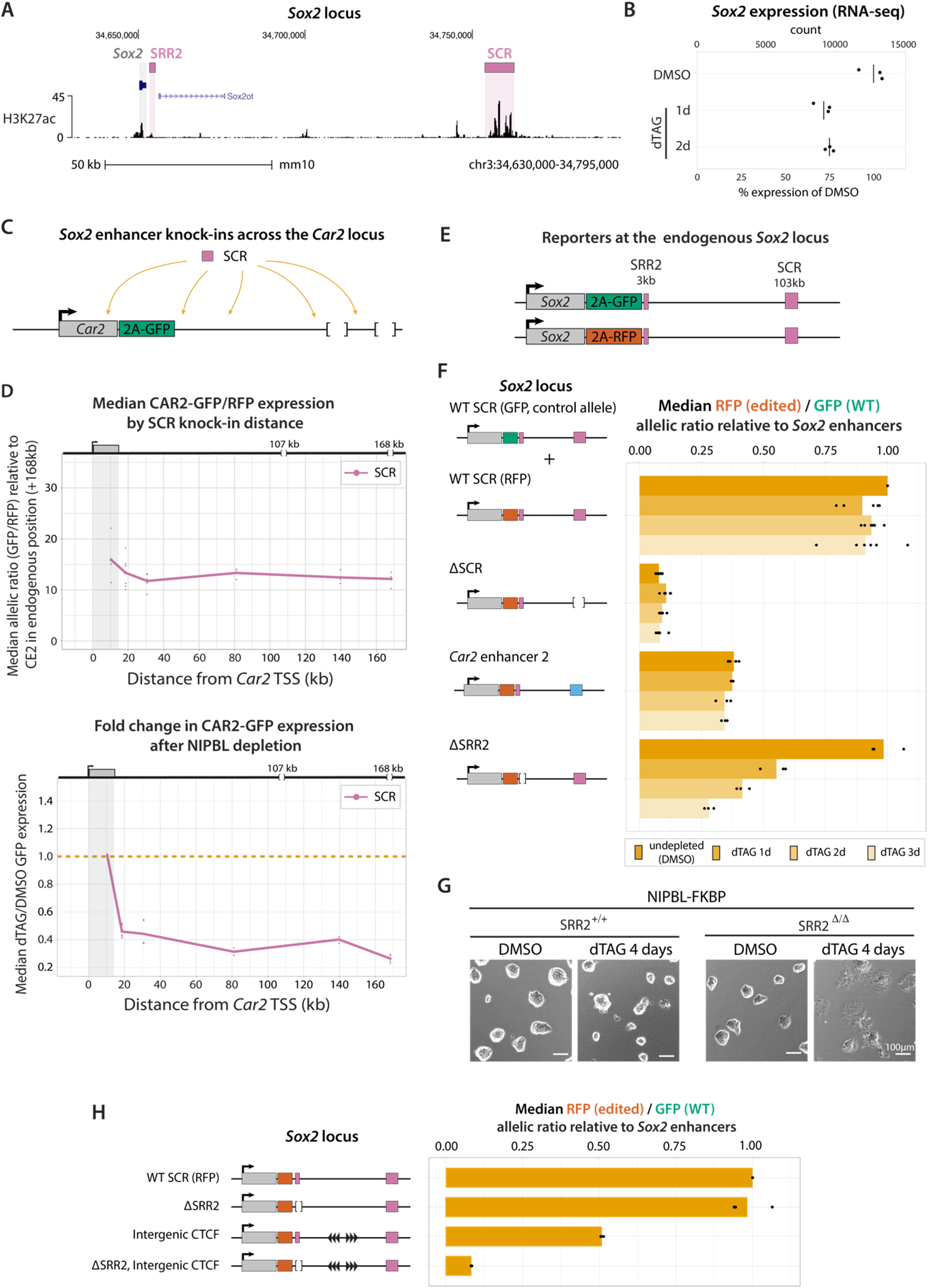
Dispensable promoter-proximal regulatory elements can support cohesin-independent communication with distal enhancers. **(A)** H3K27ac CUT&Tag at the *Sox2* locus highlighting the location of the *Sox2* Control Region (SCR) and *Sox2* regulatory region 2 (SRR2). **(B)** *Sox2* mRNA expression only decreases by ∼25% after 1-2 days of NIPBL depletion in FKBP ESCs. **(C)** Clonal ESCs generated by inserting the SCR at various locations across an enhancer-less *Car2* allele. **(D)** Inserting the SCR drove higher basal *Car2* expression than for CE1 or CE2 elements (Fig. 4 and fig. S14). These chimeric *Car2* loci driven by the SCR remained sensitive to NIPBL depletion for promoter-distal locations. **(E)** *Sox2* fluorescent reporter ESCs created by inserting 2A-GFP and 2A-RFP cassettes at the 3’ end of endogenous *Sox2* (n ≥ 3 flows). **(F)** Replacing the SCR with the CE2 element at the endogenous locus drives lower *Sox2* expression, but expression remains stable upon depleting NIPBL. In contrast, removing the SRR2 element renders *Sox2* dependent on loop extrusion, even when driven by the SCR at its endogenous location (n ≥ 3 flows). **(G)** SRR2 is required to prevent the differentiation of ESCs upon prolonged NIPBL depletion. **(H)** SCR-mediated activation of *Sox2* is only partially blocked by strong intervening CTCF sites, but deleting the SRR2 element largely prevents insulator bypass (n ≥ 3 flows).

We first considered whether inherent properties of the SCR enable it to work independently of cohesin extrusion. To test this, we inserted the *Sox2* SCR at several locations across the *Car2* TAD (Fig. 4C). We observed higher basal *Car2* expression for all positions, indicating the SCR is stronger than either of the *Car2* enhancers (Fig. 4D and fig. S14A). However, SCR-driven *Car2* expression remained equally sensitive to NIPBL depletion beyond the same 10-18 kb threshold (Fig. 4D and fig. S14B). This indicates that despite its strength, the SCR still relies on loop extrusion at the *Car2* locus. The resilience of *Sox2* to loss of extrusion is therefore unlikely to arise from properties intrinsic to the SCR.

We then tested if the *Sox2* promoter is somehow able to communicate with distal enhancers independently of loop extrusion. Replacing the *Car2* promoter with the *Sox2* promoter did not affect the dependance on loop extrusion (fig. S18), indicating that promoter sequence alone cannot account for the resilience of *Sox2* to loss of extrusion. Thus, other context features in the extended *Sox2* locus, beyond the sequence of the promoter and the main enhancer, must be at play.

To understand how the *Sox2* locus can sustain long-range regulatory communication upon NIPBL depletion, we created *Sox2* reporter ESCs with 2A-mClover (GFP) and 2A-mScarlet3 (RFP) reporters at the endogenous locus (Fig. 4E). NIPBL depletion only caused ∼15% downregulation of *Sox2* reporters within 24h (fig. S19), consistent with the mild reliance on loop extrusion identified from the transcriptomics, and in line with previous observations after RAD21 depletion (Fig. S17C) (*8*, *39*, *40*). We again used an inter-allelic normalization strategy (*30*) to normalize for secondary *trans*-effects, here manipulating the RFP allele and using the unedited GFP allele as an internal control. As reported (*31*, *32*), deleting the SCR reduced *Sox2* expression to ∼10% (Fig. 4F). *Sox2* expression was lower when we replaced the SCR with the *Car2* CE2 (Fig. 4F and fig. S20), in line with CE2 being a weaker enhancer than the SCR. Yet *Sox2* remained largely insensitive to NIPBL depletion (Fig. 4F and fig. S20), affirming that neither enhancer sequence nor strength explains the autonomy of the *Sox2* locus from cohesin loop extrusion.

We next addressed whether changing the distance between *Sox2* and the SCR would alter the reliance of the locus on loop extrusion. Relocating the SCR further away (either 265 or 364 kb) lowered basal expression, but only made the locus slightly more sensitive to NIPBL depletion (fig. S20). Even with the SCR at these distal positions and prolonged depletion, *Sox2* expression did not drop more than ∼50% after 3 days.

Reinserting the SCR closer to *Sox2* (either 4 kb or 17 kb downstream of the TSS) rendered the locus entirely insensitive to NIPBL depletion (fig. S21), demonstrating that the mild but detectable downregulation of *Sox2* when the SCR is naturally positioned at 100 kb (<25%) arises from its distal location. Yet, nlike what we observed at *Car2*, basal *Sox2* expression does not increase when the enhancer is at these proximal positions. This contrasting behavior arises from idiosyncrasies of the *Sox2* locus, not the SCR: inserting the weaker *Car2* E2 at these proximal positions did not drive higher expression than at 100 kb either, despite the basal expression being much lower than when the SCR was at the same location (fig. S21).

When inserted at an ectopic locus, the *Sox2* promoter and the SCR display a strong coupling between transcription and genomic distance (*41*). Therefore, the limited distance- and loop extrusion-dependence of the native *Sox2* locus must arise from genetic elements other than the *Sox2* promoter or the SCR.

### Promoter-proximal regulatory elements can buffer the effect of NIPBL depletion

In addition to the SCR, other elements with weak regulatory activity have been identified, both immediately upstream of *Sox2* and between *Sox2* and the SCR (*32*, *33*). These elements have some autonomous enhancer activity but were reported to only contribute marginally to basal *Sox2* expression. We wondered whether these auxiliary intervening elements may nevertheless contribute to relaying the influence of the distal SCR, in a way that would be redundant with cohesin-mediated long-range interactions and would therefore become crucial only upon defective loop extrusion. We therefore systematically removed all these elements in *Sox2* reporter cells (fig. S22A) by inducing 5 heterozygous deletions in parallel.

While basal *Sox2* expression remained largely stable across all deletions, removing the SRR2 element rendered *Sox2* expression highly reliant on NIPBL (Fig. 4F). None of the other deletions exhibited this behavior (Fig. S22B). Deleting SRR2 on an allele with the SCR relocated at 364 kb rendered *Sox2* expression nearly undetectable by 3 days of NIPBL depletion (fig. S23A). Notably, deleting SRR2 homozygously in cells without any other alteration on *Sox2* (no reporters, SCR at 100 kb) caused cells to differentiate upon NIPBL depletion (Fig. 4G), underscoring its physiological relevance for pluripotency.

Therefore, the resistance of the *Sox2* locus to disrupting loop extrusion arises from the existence of the promoter-proximal SRR2 element. SRR2, located 3 kb downstream of *Sox2*, is not bound by CTCF or cohesin (fig. S17A), but possesses weak enhancer activity: it can drive modest activity in episomal reporter assays (*32*, *42*), and is responsible for the ∼10% residual expression of *Sox2* that persists after removing the SCR (Fig. 23B). These observations indicate that synergy between proximal and distal regulatory elements plays a previously unappreciated role in the robustness of cohesin-independent long-range enhancer-promoter communication.

### Promoter-proximal regulatory elements can help bypass a strong CTCF insulator

We next asked if the SRR2 can maintain cohesin-independent communication with the SCR in other contexts beyond NIPBL depletion. For this, we inserted intervening CTCF sites between *Sox2* and the SCR to prevent cohesin from extruding across the locus. Baseline *Sox2* expression was reduced less than two-fold, highlighting the ability of the SCR to bypass the extrusion block (Fig. 4H and fig. S24A) - contrasting with the situation at *Car2,* where the same CTCF cassette completely blocked the enhancers (Fig. 3E), However, further removal of the SRR2 caused dramatic downregulation of *Sox2* (Fig. 4H and fig. S24B), with a large fraction of cells displaying nearly undetectable expression (Fig. S24C). This demonstrates that the SRR2 can also sustain cohesin-independent regulatory communication with the SCR across a strong CTCF insulator.

Altogether, these observations highlight that at least two distinct, complementary mechanisms can support long-range enhancer action: cohesin loop extrusion and synergy with promoter-proximal elements. Only in the absence of such proximal elements, as in the *Car2* locus, does cohesin extrusion prove essential for efficient regulatory communication between promoters and distal enhancers (Fig. 5). Conversely, promoter-proximal regulatory elements, as in the *Sox2* locus, can provide the genomic context that enables distal enhancers to efficiently communicate across long distances, and possibly across TAD boundaries, without the need for cohesin loop extrusion.

**Figure 5.**
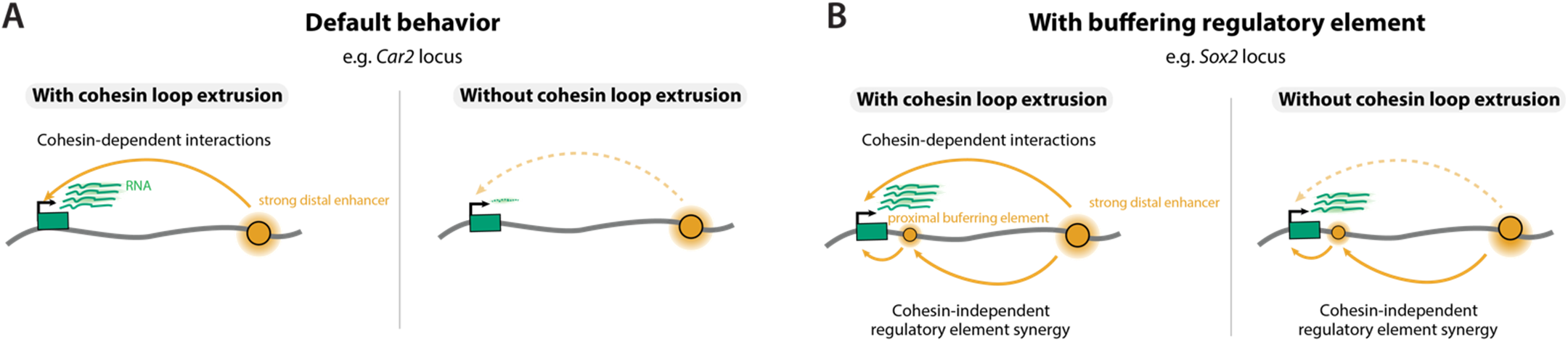
Working model explaining how genomic context can modulate the relevance of cohesin in distal enhancer-promoter communication. **(A)** At loci relying solely on essential distal enhancers (e.g. *Car2*), cohesin is crucial to enable long-range interactions, resulting in loss of transcription when loop extrusion is disrupted. **(B)** In contrast, at loci with additional regulatory elements (e.g. *Sox2*), synergy with promoter-proximal elements can compensate for cohesin disruption, forming a redundant regulatory axis that provides robustness against the loss of loop extrusion.

## Discussion

Beyond establishing novel experimental strategies to study cohesin functions in enhancer biology, our work ascertains that loop extrusion is essential for the proper regulation of a subset of gene loci, in a way that is both cell type- and genomic context-specific.

Our observations in embryonic stem cells and neural precursors demonstrate that loop extrusion can be important for the maintenance of transcriptional profiles, not simply for inducing new ones. By leveraging the differentiation potential of NIPBL-degron ESCs, we report that loci across the genome exhibit pervasive shifts in cohesin-sensitivity, in line with the highly cell type-specific nature of enhancers. Collectively, this illustrates the relevance of our findings for understanding genetic dependance across development.

Transcriptomic profiling revealed a divergent set of outcomes upon disrupted loop extrusion: while many loci like *Sox2* are minimally impacted, other loci like *Car2* are highly dependent on cohesin loop extrusion. Through systematic locus editing experiments at these contrasting loci, we identify when and how enhancer-promoter communication relies on loop extrusion. First, sensitivity to loop extrusion requires sufficiently distal and essential enhancers. Second, promoters and enhancers do not themselves specify sensitivity to loop extrusion. Third, we identify a previously unappreciated mechanism through which promoter-proximal regulatory elements can bypass loop extrusion for distal regulatory communication.

Our observations illustrate how the independent, parallel contributions of cohesin loop extrusion and proximal regulatory elements can impart robustness to long-range enhancer action: only upon disrupting both axes does *Sox2* lose the ability to communicate with the distal SCR enhancer. These findings clarify previous reports suggesting that 3D genome folding may be largely decoupled from long-range regulation, which were based on studying *Sox2* (*36*, *38*). Moreover, they explain how strong CTCF insulators can be bypassed at the *Sox2* locus (*30*, *37*, *38*). At loci without such additional promoter-proximal supporting elements, like *Car2*, 3D genome folding and long-range regulation remains tightly coupled, providing more informative and direct models to decipher the basic principles linking chromatin architecture and transcription control.

The cohesin-independence conferred by the auxiliary SRR2 element at *Sox2* constitutes a previously uncharacterized regulatory behavior. Unlike ultra long-range regulation by CTCF (*43–45*), the mechanism is cohesin-independent. Our observations instead expand the list of recently discovered of long-range regulatory mechanisms that work by potentiating enhancer activity, such as facilitation (*46*) and range-extension (*47*). However, unlike facilitators and range-extenders, which were described to lie directly within or flanking core trans-activator binding sites in (super)-enhancers, our findings describe the buffering activity of an element distinct from the core enhancer, and several kilobases away from the promoter.

Cohesin extrusion and the synergy with the auxiliary promoter-proximal element SRR2 appear redundant in supporting the action of the SCR, as *Sox2* expression in only lost upon disrupting both regulatory axes. However, we do not expect the two pathways to necessarily compensate for each other in every situation. Indeed, regulatory element synergy may become especially important for distal enhancer function in contexts where loop extrusion is naturally ineffective, for example across strong TAD boundaries or during the exit of mitosis.

Future work will clarify the biochemical basis for how promoter-proximal elements, like SRR2, synergize with distal enhancers to support cohesin-independent long-range regulation. We anticipate this will relate to how promoter-proximal enhancers can bolster the influence of distal enhancers over extensive genomic separations (*48*, *49*). In this framework, understanding how a distal enhancer delivers regulatory information to its target promoter will require accounting for the synergistic effects of nearby control elements across the extended locus, raising parallels to the notion of the holo-enhancer (*50*). Alternatively, DNA looping pathways other than cohesin may be at play (*51*, *52*).

Our observations raise further questions. How do enhancers activate promoters in the first place, and how is cohesin loop extrusion benefiting the process? Earlier genomic correlations reported that cohesin reliance scales with increased genomic distance between enhancers and their target (*17*, *22*). Together with recent high-throughput CRISPRi-based approaches to repress enhancers (*15*), our comparative genome engineering observations ascertain that cohesin is typically necessary for long-range enhancer regulation, unless additional auxiliary proximal elements introduce deviations from this behavior.

It remains unclear how physical enhancer-promoter interactions, or their proximity in 3D, modulate the transcription cascade and at which step (*53*). Long-range regulatory effects inversely scale with genomic distance, following Hi-C contact frequency (*41*, *54*). Unexpectedly, we observed a different scaling between *Car2* and *Sox2*, and recent work further demonstrated a non-uniform relationship to 3D contact frequency across genomic contexts (*55*). The list of extrusion-dependent and -independent loci provided by our transcriptomic assays will guide comparative dissection of other loci beyond *Car2* and *Sox2* (examples in Fig. S25-26). Beyond interactions detected by snapshot assays like Hi-C, loop extrusion modulates other processes that could be relevant to transcriptional regulation – from chromosome dynamics (*56*, *57*), to transcription factor diffusion (*14*, *58*, *59*), pausing of RNA polymerase II (*60*), and higher-order chromatin folding and nuclear organization (*61–63*).

Altogether, our work sheds light on when and how cohesin is required for distal enhancer-promoter communication, and provides a suite of tractable biological systems, tools, and methods to further dissect enhancer biology in native loci. Our experiments ascertain that cohesin is important for distal enhancer-promoter communication, unless additional auxiliary proximal elements introduce deviations from this base principle. The novel insight we provide guides future investigations into the molecular nature of the regulatory information delivered by enhancers to promoters – a compelling question in transcriptional regulation.

## Supporting information

Supplementary Table - CellLines-Vectors-sgRNAs

## ACKNOWLEDGEMENTS

We thank the Nora lab, Luca Giorgetti, Daniele Canzio and their groups for thoughtful discussions. We thank Barbara Panning, Nadav Ahituv, and Vijay Ramani for feedback throughout. We are grateful to Abigail Buchwalter, Benoit Bruneau, Daniele Canzio, Licia Selleri and Luca Giorgetti for comments on the manuscript. We are thankful to Benoit Bruneau for support in the independent pursuit of early observations made by EN in his laboratory. We thank the group of Akiko Hata for access to flow cytometry, the Histology and Light Microscopy Core of the Gladstone Institute for access to instruments, and the sequencing platform of the CZ Biohub San Francisco.

## AUTHORS CONTRIBUTIONS

KH and EN conceived the project. KH designed, performed, or supervised all experiments except gastruloids, with support from AA and EA for ESC manipulation and cytometry, RS for Hi-C and ChIP-seq, KZ for transcriptomics, IC for cell culture, KB for cell cycle profiling, and RMB in the lab of RB for CUT&Tag. LB designed and performed gastruloid experiments in the lab of EdW. SK and BP contributed to transcriptomic analyses in the lab of GF. MM contributed to transcriptomic analyses in the lab of EdW. KH and EN wrote the manuscript with input from all authors.

## DATA AND MATERIAL AVAILABILITY

Cell lines and plasmids can be made available from the corresponding author upon request and MTA. Key plasmids are available at https://www.addgene.org/Elphege_Nora/. Source data will be available through in NCBI’s Gene Expression Omnibus GEO.

## METHODS

### Plasmid Construction

Plasmids were assembled using Gibson assembly (NEB HiFi DNA Assembly Master Mix E2621) or restriction-ligation (NEB Quick Ligase M2200). The list of plasmids generated in this study can be found in supplementary table and their annotated sequence maps as supplementary information. Key plasmids were made available through Addgene.

### Cell Culture

Parental WT E14Tg2a (karyotype 19, XY, 129/Ola isogenic background) mouse embryonic stem cells (ESCs) and subclones were cultured in 2iSL medium [DMEM + GlutaMAX + sodium pyruvate (ThermoFisher 10569044) supplemented with 15% fetal bovine serum (Gibco SKU A5256701), 550 µM 2-mercaptoethanol (ThermoFisher 21985-023), 1X nonessential amino acids (ThermoFisher 11140-050), 10^4^ U of Leukemia Inhibitory Factor (Millipore ESG1107), 1 µM PD0325901 (Apex Bio/Fischer A3013-25), and 3 µM CHIR99021 (Apex Bio/Fischer A3011-100)]. Cells were maintained at a density of 0.2-1.5 x 10^5^ cells/cm^2^ by passaging using TrypLE (ThermoFisher 12605010) every 24-48 hours on 0.1% gelatin-coated dishes (Sigma-Aldrich G1890-100G in 1XPBS) at 37 °C and 5% CO_2_. The medium was changed daily when cells were not passaged. Cells were checked for mycoplasma infection every 6 months and tested negative. A full list of the cell lines used and generated in this study, with unique identifier numbers, can be found in as supplementary table.

To establish neural progenitors (NPCs) and astrocytes (ASCs), 0.2 million WT E14 or NIPBL-FKBP ESCs were seeded in a T25 gelatinized flask in 2iSL medium. The following day, cells were rinsed twice in 1X PBS and switched to N2B27 medium [50% DMEM/F12 medium (Gibco 31330-038), 50% Neurobasal medium (Gibco 21103-049), 1X GlutaMAX (Gibco 35050061), 0.5X B27 (Gibco 17504-044), 0.5X N2 (Millipore SCM012), and 0.1 mM 2-mercaptoethanol] and changed daily. After 7 days, cells were detached using TrypLE and seeded on nongelatinized bacterial dishes for suspension culture at 3 million cells per 75 cm^2^ and cultured in N2B27 containing 10 ng/mL EGF and FGF (Peprotech 315-09 and 100-18B). After 3 days, floating aggregates were seeded on gelatinized dishes. After 2-4 days, poorly attached aggregates were dislodged and discarded, and remaining adherent NPC cells were dissociated using Accutase (Innovative Cell Technologies AT104), passaged twice on gelatinized dishes in N2B27 + EGF/FGF and cryopreserved after expansion in 10% DMSO in N2B27 + EGF/FGF. For experiments in self-renewing NPCs, cells were thawed and cultured on gelatinized dishes in N2B27 + EGF/FGF and passaged using Accutase for 2-3 minutes at room temperature (RT). For differentiating NPCs into quiescent astrocytes, dishes were first precoated with 10 µg/mL poly-L-ornithine-hydrobromide (Sigma P4957) for 2 hours at 37°C, washed twice with PBS, and coated with 2 µg laminin (Thermofisher 23017-015) diluted in 1X PBS at least overnight at 4°C. Adherent NPC cultures were washed twice with N2B27 (no EGF/FGF), dissociated with Accutase, seeded on the pre-coated dishes, and cultured for at least 48 hours (as indicated) with N2B27 + 20 ng/µL BMP-4 (Peprotech 315-27). For CUT&Tag adherent ASCs were detached by rinsing once with 1X PBS (without Ca^2+^/Mg^2+^) and incubating for 15-20 minutes in 0.25% Trypsin + EDTA (GenClone 25-510). Trypsin was then diluted with culture medium, and cells were lifted with a cell scraper.

FKBP12^F36V^ depletion was triggered using 500 nM of PROTAC dTAG-13 (dTAG) (Sigma-Aldrich SML2601) final, diluted in culture medium. AID depletion was triggered using 500 mM of Indole-3-Acetic Acid sodium salt (IAA, auxin analog) (Tocris Bioscience 7932) final, diluted in culture medium.

### Gastruloid Culture

Before the aggregation of gastruloids, NIPBL-FKBP ESCs were maintained in 2i-LIF medium, comprising a mixture of 50% DMEM/F12 (Thermo Fisher Scientific, 11320-033) and 50% Neurobasal media (Thermo Fisher Scientific, 21103-049). This medium was supplemented with 0.5X N2 (Thermo Fisher Scientific, 17502-048), 0.5X B27 + retinoic acid (Thermo Fisher Scientific, 17504-044), 0.05% BSA (Thermo Fisher Scientific), 3 μM CHIR99021 (BioConnect), 1 μM PD03259010 (BioConnect), 2mΜ Glutamine (Thermo Fisher Scientific), 1.5 × 10^-4^ M 1-thioglycerol (Sigma-Aldrich), 100 U/mL LIF (BioConnect, GFM200-5), and 50 U/mL penicillin-streptomycin (Thermo Fisher Scientific). These cells were cultured on gelatin-coated 10 cm petri dishes within a humidified incubator set at 5% CO_2_ and 37°C. The protocol for gastruloid culture followed previously established methods(*65*). Briefly, gastruloids were formed through the aggregation of ESCs in differentiation medium (N2B27). This medium consisted of a mix of DMEM/F12 and Neurobasal medium, supplemented with 2mΜ Glutamine, β-mercaptoethanol, and 50U/mL penicillin-streptomycin. U-bottom 96-well plates (Thermo Fisher Scientific) were employed for culture, and the process was conducted in a humidified incubator maintained at 5% CO_2_ and 37°C. Cell dissociation was carried out in N2B27 medium using a serological pipette in conjunction with a 200 µL pipette tip. Subsequently, 3.75×10^4^ cells were transferred into N2B27 medium, resulting in a final volume of 5 mL. Each well received a 40 µL aliquot of this suspension, containing approximately 300 cells. After 48 hours, 150 µL of N2B27 medium containing 3 μM CHIR99021 was added using the Hamilton Star R&D liquid handling platform. Gastruloids were treated with either DMSO or 500 nM dTAG at the specified time-points. Starting from the subsequent day, the culture medium was refreshed with fresh N2B27 medium daily. At 120 hours, gastruloids were fixed with 4% formaldehyde for 30 minutes at RT (Sigma-Aldrich), underwent a single wash with PBS, and were subsequently stored at 4°C.

### Gastruloid morphological analysis

For morphological analysis, imaging was conducted at 120 hours using an inverted wide-field microscope (Zeiss Axio Observer Z1 Live). The axis ratio and area of gastruloids were measured using Fiji(*66*).

### Genome Engineering

For transfection, plasmids were prepared using the Nucleobond Midi kit (Macherey Nagel 740410.5) followed by isopropanol precipitation. Constructs were not linearized. All transfections were done using the Neon system (Thermo Fisher) with a 100 µL tip and 1 million cells at 1400 V, 10 ms, 3 pulses. To create knock-in cells, ESCs were co-transfected with 5 µg of Cas9-sgRNA expressing vector and 10-15 µg of targeting construct. If more than one Cas9-sgRNA or targeting construct was used we used 2.5 µg and 7.5 µg each, respectively.

sgRNAs were typically cloned into a Cas9-2A-puro vector (pEN243, identical to pX459 Addgene 62988). Homology arms of targeting vectors ranged from 800 base pairs (bp) - 1 kilobase (kb) in length.

After electroporation, cells were seeded in a 9 cm^2^ well and left to recover for 48 hours. Cells were plated at limited dilution and grown for around 8 days changing medium every other day. until single colonies could be picked. Individual colonies were genotyped by polymerase chain reaction (PCR) and validated with Sanger sequencing. Homozygous or heterozygous clones were identified, expanded, validated by western blot or flow cytometry, and cryopreserved. For serial editing, resulting validated clonal lines were used to repeat the transfection process with another targeting construct.

When the targeting construct contained a resistance cassette, we added antibiotic in the medium from the moment they were serially diluted until colony picking. Final antibiotic concentrations: Puromycin: 1 µg/mL; Blasticidin: 10 µg/mL; Hygromycin: 200 µg/mL; Geneticin: 200 µg/mL.

When the targeting construct did not contain a resistance cassette, we used sgRNAs cloned into the Cas9-2A-puro vector (pEN243, identical to pX459 Addgene 62988) and used puromycin for 48 hours starting 24 hours after transfection. This was not strictly necessary as transfection efficiencies were typically above 80% but can help kill off untransfected cells to increase the chance of picking knock-in clones.

Cas9 cutting between *Sox2* and its ESC enhancer, the SCR, results in transient (1-2 day) downregulation of *Sox2*. For cell lines with heterozygous genomic edits at the *Sox2* locus, a Cas9-2A-puro vector with a sgRNA targeting within the deletion or knock-in was transfected using L2000. Briefly, 120 ng of Cas9-sgRNA plasmid was added to 60 uL of Opti-MEM per sample and 0.6 uL of Lipofectamine 2000 (L2000) reagent was added to 60 uL of Opti-MEM per sample and incubated in the dark for 5 minutes. The DNA and L2000 mixtures were combined and incubated in the dark for 1 hour. 120 uL of the DNA-L2000 was then added to 0.1M freshly seeded ESCs in 2 cm^2^. Decreased *Sox2* expression was detected by flow cytometry 24 hours post-transfection to identify the edited allele (fig. S27).

The list of cell lines, genotypes, order of transfection and corresponding vectors is provided as supplementary table.

### Cell cycle analysis by EdU incorporation and DAPI staining

The Click-iT Plus EdU Alexa Fluor 647 Flow Cytometry Assay Kit (Thermo Fisher Scientific C10634) was used following the manufacturer’s instructions. We used 20 µM EdU, pulsed for 1 hour and 30 minutes, and proceeded with 1 million cells, incorporating DAPI after EdU staining.

### Cell proliferation

10,000 cells were seeded in a single well of a 6-well plate. Medium was changed daily -/+dTAG and cells were manually counted with an Hemacytometers (Medon Surgical DHCN015) 4 days after the initial seeding.

### Crystal violet staining for clonogenicity

Around 100 cells were seeded into a well of a 6-well plate, seeding two distinct wells from the same cell suspension in parallel (one for ‘untreated’ and one for ‘dTAG’ condition). Medium was changed daily -/+ dTAG for around 7 days. Plates were then washed with PBS and fixed with diluted in 1% Formaldehyde 1% Methanol 0.05% crystal violet (diluted from 1% solution, Aldon Corp SE cat CC0455-100ML) for 20 min. Plates were thoroughly rinsed with tap water and air dried and photographed with a cell phone camera.

### Nuclear morphology by DAPI staining

Cells were grown for 24h in ibidi chambers, washed with 1X PBS, fixed with 3% formaldehyde diluted in 1X PBS (Electron Microscopy Sciences Cat 100503-912), washed three times with 1XPBS, permeabilized with 0.5% Triton-X diluted in 1X PBS, incubated in 1ug/mL DAPI (Adipogen Corporation 501687572) diluted in 1X PBS, washed twice with 1X PBS and stored at 4°C. Imaging was performed on a Zeiss spinning disk with 60× objective and 405nm laser.

### Flow cytometry

ESCs were dissociated with TrypLE, resuspended in culture medium, spun, and resuspended in 10% FBS-PBS before live cell flow cytometry on an Attune NxT instrument (Thermo Fisher). Dissociation, wash, and flow buffers were supplemented with dTAG, when appropriate, to avoid re-expression of the NIPBL-FKBP fusion. Analyses were performed using the FlowJo software. Briefly, live cells were gated using FSC-H x SSC-A followed by the isolation of singlets using FSC-H x FSC-W before extracting statistics for approximately 50,000 final gated cells. To account for any effects in *trans* in the dual fluorescent reporter cell lines, the population median expression of the edited allele was divided by the population median of the unedited allele with E14 untagged autofluorescence removed, all relative to a control cell line. For a subset of the flow cytometry, the single-cell fluorescent ratios were evaluated against the population-level calculation and the results were comparable. Thus, utilizing the population medians enabled the subtraction of autofluorescence and facilitated the rapid calculation of fluorescent ratios without the need to extract single-cell data. Replicate datapoints were obtained by growing and flowing the same cell line on different days. For each cell line two independent clones are presented in supplementary information.

### Western blotting

ESCs were dissociated using TrypLE, resuspended in culture medium, pelleted at 100*g* for 5 minutes at RT. Cell pellets were washed in PBS, pelleted again, snap frozen in dry ice and stored in −70°C freezer. In total, 10-20 million cells were used to prepare nuclear extracts. Cell pellets stored at −70°C were taken out on ice, resuspended in 10 mM HEPES (pH 7.9), 2.5 mM MgCl_2_, 0.25 M sucrose, 0.1% NP40, 1 mM DTT, and 1X HALT protease inhibitors (ThermoFisher 78437) and kept on ice for 10 minutes to allow swelling. After centrifugation at 500*g* at 4°C for 5 minutes, the supernatant was discarded and the pellets were resuspended on ice in 25 mM HEPES (pH 7.9), 1.5 mM MgCl_2_, 700 mM NaCl, 0.5 mM DTT, 0.1 mM EDTA, 20% glycerol, 1 mM DTT, and 250 U benzonase (Sigma-Aldrich E1014), and incubated on ice for 30-60 minutes. The nuclear lysates were centrifuged at 18,000*g*, 4°C, 10 minutes, to eliminate the cellular debris. Protein concentration from the supernatants was measured using Pierce BCA Kit (ThermoFisher 23225).

Equal amounts of protein lysates were heated in 1X Laemmli buffer (Biorad 1610747) at 95°C for 10 minutes and subjected to electrophoresis on a 4-15% polyacrylamide gel (Biorad 17000927). For NIPBL westerns, 60 µg protein was used. The separated proteins were transferred onto a polyvinylidene difluoride (PVDF) membrane (Millipore IPFL00010) using phosphate-based transfer buffer (10 mM sodium phosphate monobasic, 10 mM sodium phosphate dibasic) at 4°C, 600 mA, 2 hours. After the completion of transfer, membranes were blocked in 5% blocking buffer (5% skimmed milk prepared in 1X TBS-T) and incubated overnight at 4°C with the primary antibodies prepared in 5% blocking buffer. Next day, the membranes were washed three times with 1X TBS-T buffer (20 mM Tris-Cl buffer-pH 7.4, 500 mM NaCl and 0.1% Tween 20) and incubated with the appropriate secondary antibodies conjugated with horseradish peroxidase (HRP) for an hour at RT. Following this, the membranes were again washed three times with 1X TBS-T buffer. The blots were developed using Pierce ECL western blotting substrate (ThermoScientific 32209) and detected using Chemidoc XRS+ imager (Biorad).

### RNA-seq library generation

For ESCs, 0.5 million E14 or NIPBL-FKBP cells were seeded in 9 cm^2^ with DMSO or dTAG and collected after either 24 or 48 hours for RNA extraction. For NPCs 0.22 million E14 or NIPBL-FKBP cells were seeded in 9 cm^2^, either at time of seeding (48-hour depletion) or after 1 day (24-hour depletion) medium was changed and supplemented with either DMSO or dTAG. Cells were collected for RNA extraction 2 days after seeding. For the ESC exit from pluripotency differentiation, 0.25 million NIPBL-FKBP ESCs grown in 2iSL were seeded in 9 cm^2^. DMSO or dTAG was added to the culture medium 3 hours after seeding. Once cells had attached, 6 hours after seeding, cells were washed twice with 1X PBS and changed into differentiation medium (N2B27) supplemented with DMSO or dTAG. For the ASC differentiation, NIPBL-FKBP NPCs were treated with DMSO or dTAG for 3 hours. Then, 0.1 million NPCs were washed twice with N2B27 (without EGF/FGF) and seeded in 9 cm^2^ in differentiation medium (N2B27 + BMP4) with either DMSO or dTAG. Cells were collected for RNA extraction 2 days after seeding for both differentiations.

For collection, cells were washed with PBS and lysed directly with 1 mL TRIzol (Thermo Fisher 15596-018), transferred to a 1.5 mL tube, vortexed, and incubated at room temperature for 5 minutes. After incubation, mixture was supplemented with 250 µL Chloroform, vortexed again, and centrifuged at 12,000*g* at 4°C for 15 minutes. The upper phase was transferred to a new tube, mixed with an equal volume of isopropanol, and spun at 12,000*g* at 4°C for 15 minutes. The pellet was washed with 70% ethanol, air dried, and resuspended in 25 µL water. RNA quality was assessed by running 200 ng on a 2% agarose gel, quantified using Qubit dsDNA High Sensitivity kit (Invitrogen Q32854), and stored at −70°C before moving on to library preparation.

2 µg of total RNA was DNase treated with Turbo DNase kit (Invitrogen AM1907). 100-500 ng of total RNA was used as input for poly(A) capture and library preparation using the Illumina Stranded mRNA kit (Illumina 20040532) per manufacturer’s instructions, and DNA quality was assessed using the Bioanalyzer High Sensitivity DNA kit (Agilent Technologies 5067-4626). Libraries were sequenced using paired-end 100 bp on HiSeq4000 or NextSeq2000 to obtain a minimum of 20M reads. A subset of libraries was sequenced using paired-end 150 bp on a NovaSeq6000 instrument and trimmed to 100 bp prior to analysis.

### RNA-seq analysis

Reads were trimmed to 100 bp when needed using cutadapt(*67*). Alignment of raw RNA-seq data was produced using STAR version 2.7.7a (*68*) with default parameters and basic two-pass mapping in mm10. GENCODE VM23 (GRCm38.p6) was used as the reference gene set. Size factor normalization and differential expression analyses were done using DESeq2 version 1.36.0 (*69*) after filtering out genes with <10 raw reads. Only genes with a False Discovery Rate (FDR) < 0.05 and fold change (FC) ≤ −1.5 or ≥ 1.5 were called as differentially expressed. Gene set enrichment analysis was performed using clusterProfiler v4.4.4 (*70*) for biological processes.

To compute the density of features around dysregulated transcription start sites (TSSs), positions were expanded by 1Mb and overlapped with positions of ChIP-seq peaks using bioframe (*72*). Distances between peak positions and TSSs were then binned into 40 bins of 25 kb each and computed separately or up/down/nonsignificant categories with numpy (*73*) and visualized with matplotlib (*74*).

To predict the differential expression of genes as up, down, or nonsignificant, logistic regression models were trained with L2 regularization using *scikit-learn* (*75*) using features around each TSS. ChIP-seq or CUT&Tag peak positions +/-1 Mb around each TSS were binned by distance into 40 bins of 50 kb each to generate a feature set. TSSs were split into train/test sets with a 4:1 ratio. To address class imbalance in the training data, the RandomUnderSampler method from the imbalanced-learn library (*76*) method was used. To evaluate model performance, Area Under the Receiver Operating Characteristic (AUC-ROC) curve was computed using *scikit-learn* on the test dataset.

### ATAC-seq library generation on gastruloids

The Tn5 protein purification was carried out by the NKI protein facility as previously described(*27*). Transposon annealing was achieved by combining 10 µL of 10X TE buffer with 45 µL each of 100 µM Tn5MErev oligonucleotides and 100 µM corresponding Tn5ME-A and Tn5ME-B oligonucleotides. The adapter solution was then subjected to incubation at 95°C for 10 minutes followed by gradual cooling to 4°C at a rate of 0.1°C per second. For transposome formation, the annealed adapters were diluted with an equal volume of H2O. This resulting adapter solution was mixed with 0.2 mg/mL Tn5 at a 1:20 ratio and incubated for 1 hour at 37°C. The transposomes were either utilized immediately for tagmentation or stored at −20°C for up to 2 weeks. Subsequently, gastruloids were dissociated into a single-cell suspension through gentle trypsinization and frozen with the addition of 10% DMSO and 10% fetal bovine serum. These samples were then stored at −70°C until further use. A total of 50,000 cells were collected in cold PBS and lysed using a 2X lysis buffer (1M Tris-HCl pH 7.5, 5M NaCl, 1M MgCl_2_, 10% IGEPAL). Following centrifugation, the resulting pellet was treated with 2X TD buffer and 2 µL of transposon mix. PCR amplification was performed twice using KAPA HiFi HotStart ReadyMix (Roche) and P5 and P7 indexed primers (please refer to the oligonucleotide list). Fragments ranging from 200 to 700 bp were purified using AMPure XP beads (Beckman Coulter), and the DNA quality was assessed through Bioanalyzer High Sensitivity DNA analysis (Agilent).

### ATAC-seq analysis

ATAC-seq reads underwent mapping using bwa mem v0.7.17-r1188123 to the mm10 mouse reference genome assembly(*77*). Reads that were uniquely mapped and not assigned to mitochondrial DNA, with MAPQ > 10, were retained through selection using SAMtools v1.12 (*78*). Duplicate reads were then removed using the Picard v2.25.6 “MarkDuplicates” function. To decipher the cell type compositions within bulk ATAC-seq samples from both control and NIPBL-depleted gastruloids, a previously published single-cell ATAC-seq atlas of gastruloid development was used(*27*). Initially, pseudobulk ATAC-seq profiles for each cell type (cluster) were generated from single-cell data by summing counts in each peak across all cells belonging to the same clusters. Subsequently, read counts of DMSO and dTAG treated bulk samples within the same peak set utilized in the single-cell data were generated. These bulk and pseudobulk counts were then concatenated, normalized using the median of ratios normalization method implemented in DESeq2 v1.30.1147(*69*), and transformed using the variance stabilizing transform (VST). Since bulk samples represent linear combinations of various cell types, independent component analysis (ICA) was employed to decipher the contributions of constituent cell types in the bulk samples. ICA is a method for segregating a multivariate signal into a predefined number of additive, mutually independent components. Briefly, ICA was conducted on VST-transformed and zero-centered pseudobulk counts using the FastICA algorithm, implemented in the “icafast” function from the ica R package. The number of components was set to 7 with a maximum of 100 algorithm iterations. The source signal estimates and the unmixing matrix obtained from ICA were then applied to the VST-transformed bulk count matrix to deconvolve the cell type contributions. Finally, cell type proportion estimates were derived by setting the negative contributions to zero and normalizing contributions to the sum of the contributions for each sample.

### CTCF Calibrated ChIP-seq

For crosslinking, ESCs with CTCF site deletion in the *Car2* promoter (KH105.1 and KH105.2) were dissociated using TrypLE, resuspended in culture medium and counted. 15 x 10^6^ ESCs were pelleted at 100*g* for 5 minutes at RT. Supernatant was discarded and cell pellets were resuspended in 5 mL of 10% FBS prepared in 1X PBS. To this, 5 mL of freshly prepared 2% formaldehyde solution was added to get final formaldehyde concentration of 1%. 2% formaldehyde was diluted from 16% methanol-free formaldehyde in 1X PBS (ThermoFisher 28908). Resuspended cells were incubated on rocker to allow for mixing for 10 minutes at RT. Quenching was performed using 0.125M Glycine prepared in 1X PBS and incubated on rocker for 5 minutes at RT. Tubes were then stored on ice for 15 minutes followed by centrifuging at 100*g* for 5 minutes at 4° C. Supernatant was discarded and cell pellets were washed twice with 5 mL of ice-cold 1X PBS. Cell pellets were resuspended in 1 mL of ice-cold PBS and transferred to 1.5 ml tubes, centrifuged at 100*g* for 5 minutes at 4° C. Supernatant was discarded and pellets were snap frozen on dry ice and stored at −70°C.

Fixed cells were thawed on ice, resuspended in 1 mL of ice-cold lysis buffer [10 mM Tris HCl pH 7.5, 10 mM NaCl, 0.2% NP-40 (IGEPAL), 0.2% TritonX-100, 1 mM EDTA, 0.5 mM EGTA, supplemented with fresh protease inhibitor] and incubated on ice for 10 minutes. 9.75 x 10^6^ ESC nuclei were transferred to a pre-chilled 1.5 mL tube and 0.25 x 10^6^ crosslinked HEK nuclei were added as spike-in. Nuclei were pelleted at 400*g* for 5 minutes at 4°C, supernatant discarded and nuclear pellets resuspended in 900 μL of shearing buffer (0.5% SDS, 10 mM Tris HCl pH 7.5, supplemented with fresh protease inhibitor). Chromatin was sheared using Covaris with settings PIP 105, Duty factor 5, Cycles per burst 200, total time 15 minutes. After shearing, samples were collected in pre-chilled 1.5 mL tubes and spun at 16,000*g* for 5 minutes at 4° C. Supernatant was collected in a fresh tube. 5% sample was removed as Input and stored at −20°C.

For immunoprecipitation (IP), 40 μl Dynabeads/ChIP were prepared by washing them once in 1 mL of 1X ChIP incubation buffer (diluted from 5X ChIP incubation buffer in 1X PBS; 5% TritonX-100, 0.75 M NaCl, 5 mM EDTA, 2.5 mM EGTA, 50 mM Tris-Cl pH7.5) and resuspended in the 40 μL of same buffer. IP reaction was assembled as follows: 800 μL chromatin + 40 μL beads + 10 μL CTCF antibody at 1μg/ μL + 22.5 μL protease inhibitor + 442 μL 5X ChIP incubation buffer + 935.5 μL water For IP, tubes were incubated overnight on end-to-end rocker at 10 rpm, 4°C.

Following overnight incubation, beads were collected using magnetic rack and washed sequentially with different wash buffers for 5 minutes each on end-to-end rocker at 10 rpm, 4°C. Beads were washed twice with 500 μL of cold wash buffer 1 (0.1% SDS, 0.1% Sodium deoxycholate, 1% TritonX-100, 0.15 M NaCl, 1 mM EDTA, 0.5 mM EGTA, 10 mM Tris pH 8). Beads were then washed once each with 500 μL of cold wash buffer 2 (0.1% SDS, 0.1% Sodium deoxycholate, 1% TritonX-100, 0.5 M NaCl, 1 mM EDTA, 10 mM Tris pH 8) and 500ul of cold wash buffer 3 [0.25 M LiCl, 0.5% Sodium deoxycholate, 0.5% NP-40 (IGEPAL), 1 mM EDTA, 0.5 mM EGTA, 10 mM Tris pH 8]. Finally, beads were washed twice with 500 μL of cold wash buffer 4 (1 mM EDTA, 0.5 mM EGTA, 10 mM Tris pH 8).

For elution, beads were resuspended in 200 μL of elution buffer (1% SDS, 0.1M NaHCO3) and incubated on end-to-end rocker at 10 rpm for 20 minutes, RT. Beads were then separated using magnetic rack and the eluate (supernatant) was transferred to a fresh tube. 5% input samples were thawed on ice and elution buffer was added to bring the volume to 200 μL. To each sample, 2 μL of 1 mg/ml RNaseA was added and incubated at 37°C for 30 minutes. For reverse crosslinking, 8 μL of 5 M NaCl and 5 μL of 10 mg/mL Proteinase K was added and incubated overnight at 65°C. Following reverse crosslinking, DNA was eluted using MinElute kit from Qiagen (28604). Eluted DNA was quantified using Qubit and libraries were prepared using NEBNext Ultra II DNA library prep kit as per manufacturer’s protocol (E7645S). All of ChIP samples and 4 ng of Input samples were used for library prep. Final library amplification was done for 17 cycles, followed by 0.9X SPRI bead purification (Bulldog Bio CNGS005). Eluted in 10 μL of 10 mM Tris-Cl pH7.5. Entire eluate was run on a 2% E-gel (Invitrogen G401002) and fragments between 200-600 bp were excised and eluted using MinElute gel extraction kit. Quality and quantity of libraries were assessed using Qubit and Bioanalyzer High Sensitivity DNA kit. Libraries were sequenced using single-end 73 bp on NextSeq 2000.

### Calibrated ChIP-seq analysis

Fastq files were aligned to the catenated mm10 and hg38 reference genome with Bowtie2 (*79*). Reads with a mapq score ≥ 30 were retained and duplicates were removed, using Samtools. Bigwigs were generated with the bamCoverage function from deepTools (*80*) with scaling factors calculated using the formula (human reads in input) / (mouse reads in input) / (human reads in ChIP) x 15000000 (*81*, *82*). The multiplier of 15000000 was chosen arbitrarily to make the final scaling factors close to 1.

### Hi-C sample preparation

Hi-C was performed using the kit from Arima Genomics per manufacturer’s instructions (202103–1577). Briefly, ESCs were dissociated using TrypLE, resuspended in culture medium and counted. 5-10 million cells were spun down at 100*g* for 5 minutes at RT. Supernatant was discarded and the cell pellets were resuspended in 5 mL of 1X PBS. Cells were crosslinked using 2% formaldehyde (diluted from 37% formaldehyde - Electron Microscopy Sciences 15686), mixed by inverting the tubes 10 times and incubated at RT for 10 minutes. Quenching was done using the Stop solution 1 provided with the kit, mixed by inverting the tubes 10 times and incubated at RT for 5 minutes. Samples were then incubated on ice for 15 minutes, spun down at 100*g* for 5 minutes at 4°C. After discarding the supernatant, cells were resuspended in cold 1X PBS, counted, aliquoted at a density of 1×10^6^ cells/tube and spun down at 100*g* for 5 minutes at 4°C. Supernatant was discarded, cell pellets were snap frozen on dry ice and stored at −70°C.

Fixed cells were thawed, and Hi-C was performed per manufacturer’s guidelines. Libraries were prepared using TruSeq DNA Nano LP kit from Illumina (20016328). Quality and quantity of libraries were assessed using Qubit and Bioanalyzer High Sensitivity DNA kit. Libraries were sequenced on NextSeq 500 using paired-end 75 bp or NextSeq 2000 using paired-end 101 bp (and then trimmed to 75 bp before analysis).

### Hi-C analysis

Reads were trimmed to 75bp (when sequenced beyond that) using *cutadapt* (*67*). Data was mapped to mm10 using the *distiller-nf* pipeline (https://github.com/open2c/distiller-nf), iteratively corrected and converted into .cool format with *cooler* (*83*), only considering read pairs with both sides mapping with high confidence (mapq ≥ 30). After ascertaining the overall similarity of each replicate using either *cooltools* (*84*) or *Genova*(*85*) (P(s) curves and aggregate peak analysis), the independent Hi-C replicates (up to two) were merged for each condition. Multiresolution coolers (.mcool) were then generated with *cooler* and balanced using iterative correction. P(s) plots and Hi-C contact maps were generated using *cooltools* (*84*). Aggregate TAD analysis was performed with the ATA function of *Genova* (*85*) using the TAD list for ESCs from Bonev et al. 2017 (*86*). Hi-C peaks (also referred to as ‘Hi-C dots’ or ‘Hi-C loops’) were identified from the merged .mcool of WT (untagged) cells with *mustache* (*87*) using -r 10000 -st 0.8. For aggregate peak analysis the APA function of *Genova* (*85*) was used. The loop spectrum analysis was performed by extracting, for each sample, the observed/expected fold change at the Hi-C peaks detected in WT untagged cells, and plotted as a function of the genomic distance separating the two anchors.

### CUT&Tag sample preparation

Set up for cell collection was the same as RNA-seq, except 0.5 million ESCs were seeded in gelatinized T25 flasks for ESCs and the exit from pluripotency differentiation. For NPCs and ASC differentiations, 1.3 million and 0.6 million NPCs were seeded in prepared 10 cm plates, respectively.

CUT&Tag was conducted using the protocol from (*88*)with the following modifications: freshly collected cells were bound to Concanavalin A beads at a ratio of 2 x 10^5^ cells/7 μL beads in CR wash at RT. At least 1 x 10^5^ cells were used as input per sample. Bead-bound cells were then incubated rotating overnight at 4°C in CT Antibody buffer (CR Wash with 0.05% Digitonin, 2 mM EDTA, 1 mg/mL BSA) containing primary antibody. We used the following primary antibodies for CUT&TAG: 1:100 rabbit anti-H3K27ac (Abcam, ab4729) and 1:100 rabbit anti-H3K4me3 (Cell Signaling 9733). After primary, samples were washed 3 times for 5 minutes each using CR Dig-wash buffer and resuspended in 1:100 secondary antibody (Guinea pig anti-rabbit, Antibodies Online #ABIN101961) in CR Dig-wash buffer at 4°C for 1 hour rotating at 4°C. Samples were then incubated for 1 hour at 4°C with 50 μL of approximately 25 nM homemade pA-Tn5 in CT Dig300 wash buffer (20 mM HEPES, 300 mM NaCl, 0.01% Digitonin, 0.5 mM Spermidine with Roche cOmplete protease inhibitors added). Recombinant Tn5 was purified and loaded with adapters as previously described (*88*). After Tn5 incubation, samples were washed 3 times for 5 minutes each with CT Dig300 wash buffer. Tagmentation was then initiated for 1hr at 37°C in a thermocycler by adding MgCl_2_ to 10 mM final concentration in 50 μL volume. The tagmentation reaction was quenched immediately afterwards by adding 1.6 μL l of 0.5 M EDTA, 1 μL of 10 mg/mL Proteinase K, and 1 μL of 5% SDS. Samples were then incubated at 55°C for 2 hours in a thermocycler to denature Tn5 and solubilize tagmented chromatin. After incubation, samples were magnetized, and the supernatant was transferred to new wells where SPRI bead purification was performed using homemade beads to select all DNA fragment lengths larger than 100 bp. Samples were eluted in 0.1X TE and approximately half of each sample was used for library preparation using NEBNext HIFI Polymerase with custom indices synthesized by IDT. An appropriate number of cycles for each target was chosen to prevent overamplification bias. After amplification, libraries were purified with 1.2X homemade SPRI beads to select for fragments >250 bp and eluted in 0.1X TE. Quality and concentration of libraries were determined by an Agilent 4200 Tapestation with D1000 reagents before pooling for sequencing.

### CUT&Tag analysis

Fastq files for CUT&Tag samples were processed using Nextflow (*89*) and the nf-core CUT&RUN pipeline (*90*, *91*) v3.1. In brief, adapters were trimmed using Trim Galore. Paired-end alignment was performed using *Bowtie2* (*79*)and peaks were called using SEACR (*92*) with a peak threshold of 0.05 using spike in normalization performed using the *E. coli* genome K12.

Peaks in ENCODE blacklisted regions (*93*) were removed using the intersect function in *bedtools2* (*94*) v2.30.0. For genome-wide profiles and peak overlaps, peaks were filtered for a minimum signal of 1,000 and then generated using the plotProfile function in *deepTools2* (*80*) centered on peaks called from control (DMSO) cells and separated by replicate. Overlapping peaks between control and dTAG treated NIPBL-FKBP cells were determined using the intersect function in *bedtools2* with a minimum overlap fraction of 0.5.

### Liftover of published datasets to mm10

Public datasets used are listed in supplementary table. Processed .bw files not in mm10 were converted using the UCSC Liftover tool.

### Antibodies

For Western:

NIPBL sc-374625

MAU2 ab183033

### TBP CST 8515S

For ChIP-seq:

CTCF Active motif 61311

For CUT&Tag:

H3K27ac Abcam ab4729

H3K27me3 Cell Signaling 9733

---

**Figure S1.**
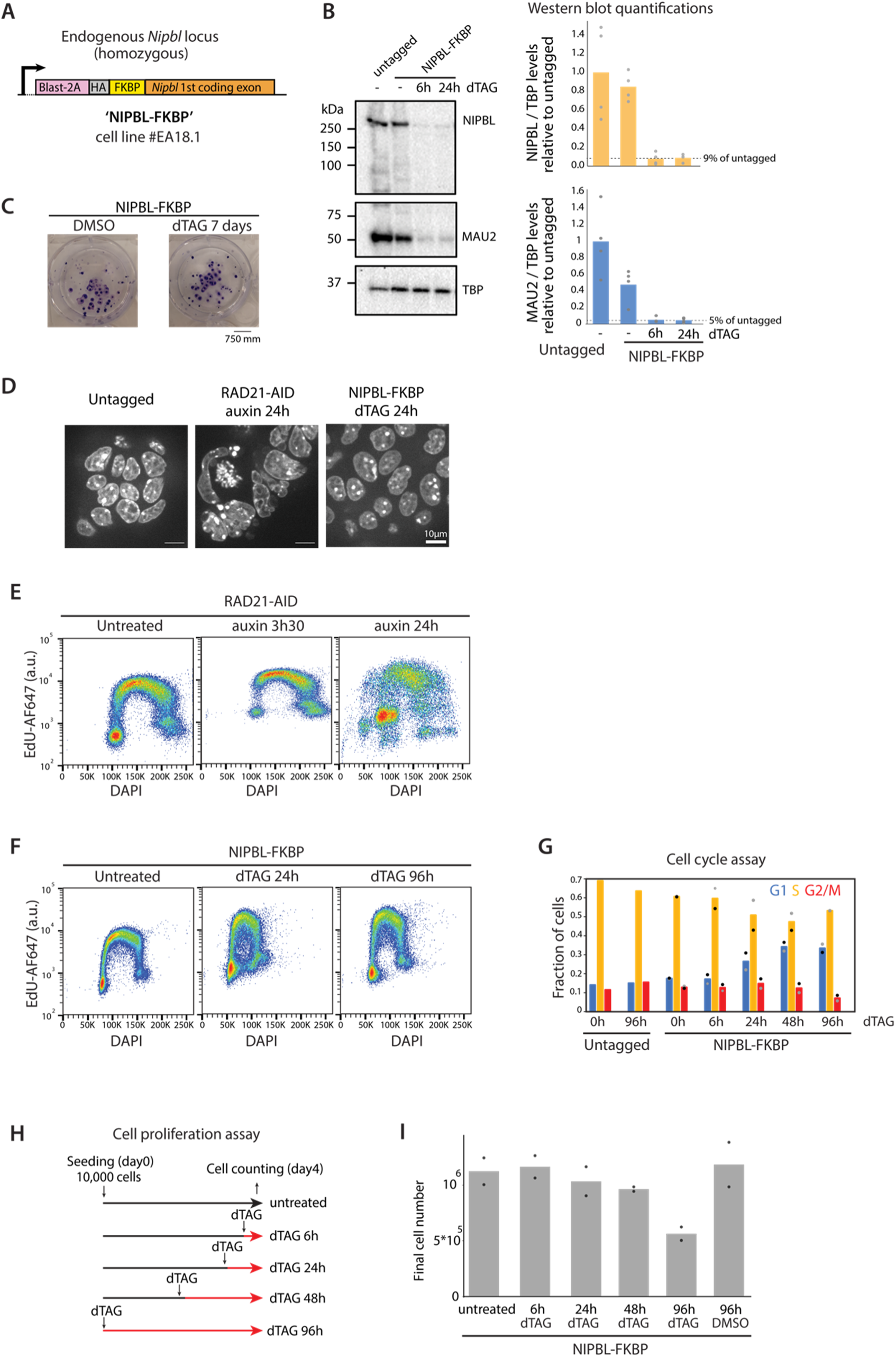
NIPBL degradation preserves the proliferative capacity of ESCs, in contrast to cohesin degradation. **(A)** Schematic of the NIPBL-FKBP degron cell line (#EA18.1). The *Nipbl* gene was edited to express blasticidin resistance, an HA-tag, and the FKBP12 inducible degron on the NIPBL N-terminus. **(B)** Western blot for NIPBL and MAU2 in untagged (E14) and NIPBL-FKBP ESCs treated with dTAG for 6 and 24h (left) and quantification of protein levels relative to untagged (right). Depletion leaves around 10% of NIPBL as detected with this N-terminal sc-374625 antibody (see Methods). **(C)** Crystal violet staining showing that NIPBL-depleted cells retain clonogenicity even after 7 days of depletion and limiting dilution. Depleting NIPBL for 24 hours (h) does not alter overall nuclear morphology as monitored by DAPI staining, in contrast to depleting the core cohesin subunit RAD21. **(E)** Flow cytometry after 1.5h of EdU incorporation and cohesin depletion in RAD21-AID cell cycle flow. Ploidy is completely aberrant after 24h of cohesin depletion. **(F)** In contrast NIPBL-FKBP dTAG-treated cells have near-normal progression through cell cycle. **(G)** NIPBL-FKBP cell cycle quantification. **(H)** Cell proliferation assay schematic. **(I)** Cell proliferation appears normal up to 48h depletion but slightly decreases 96h after depletion.

**Figure S2.**
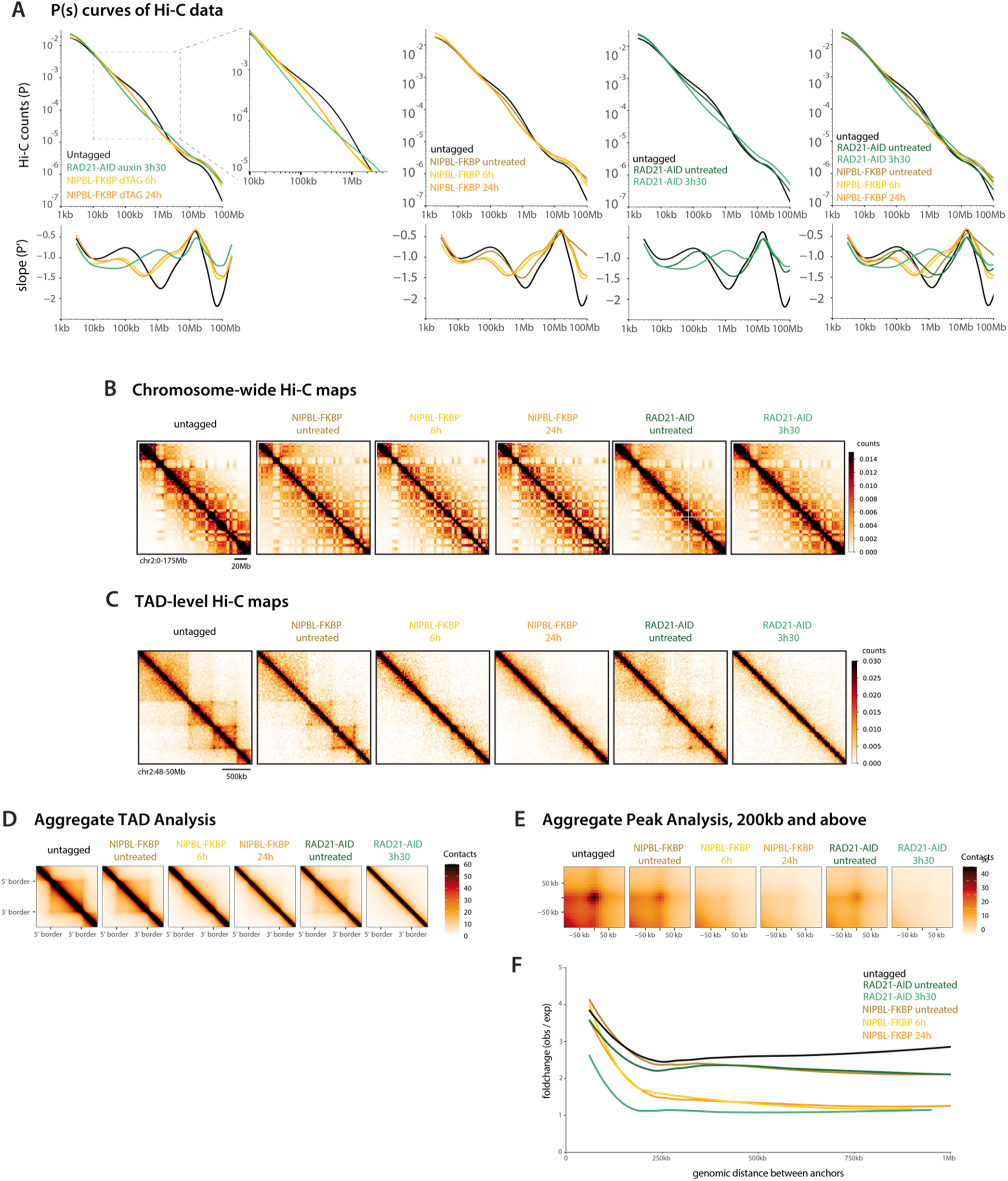
Supporting Hi-C data in NIPBL- and RAD21-degron ESCs. **(A)** P(s) curves of Hi-C data highlighting the decrease in long-range interactions around the 100 kb-1 Mb scale after NIPBL depletion. The NIPBL-FKBP ESCs appear minimally but detectably leaky, in line with the slight reduction of basal NIPBL levels observed by Western blot (Fig. 1). Bin size 1 kb. **(B)** Example of chromosome-wide Hi-C maps highlighting maintenance of large-scale chromosome structure. Bin size 1 Mb. **(C)** TAD-level Hi-C maps. Bin size 20 kb. **(D)** Aggregate TAD analysis showing loss of TAD level interactions after NIPBL depletion. Bin size 10 kb. **(E)** Aggregate peak analysis (200 kb and above) at Hi-C loops (peaks) detected in untagged cells (n=9671), showing loss of looping after NIPBL depletion. Bin size 10 kb. **(F)** Observed/expected Hi-C signal at Hi-C loops (peaks) detected in untagged cells (n=9671) as a function of the genomic separation between the two anchors. NIPBL depletion reduces the signal across all analyzed distances, eventually approaching the levels expected from genomic separation at large distances.

**Figure S3.**
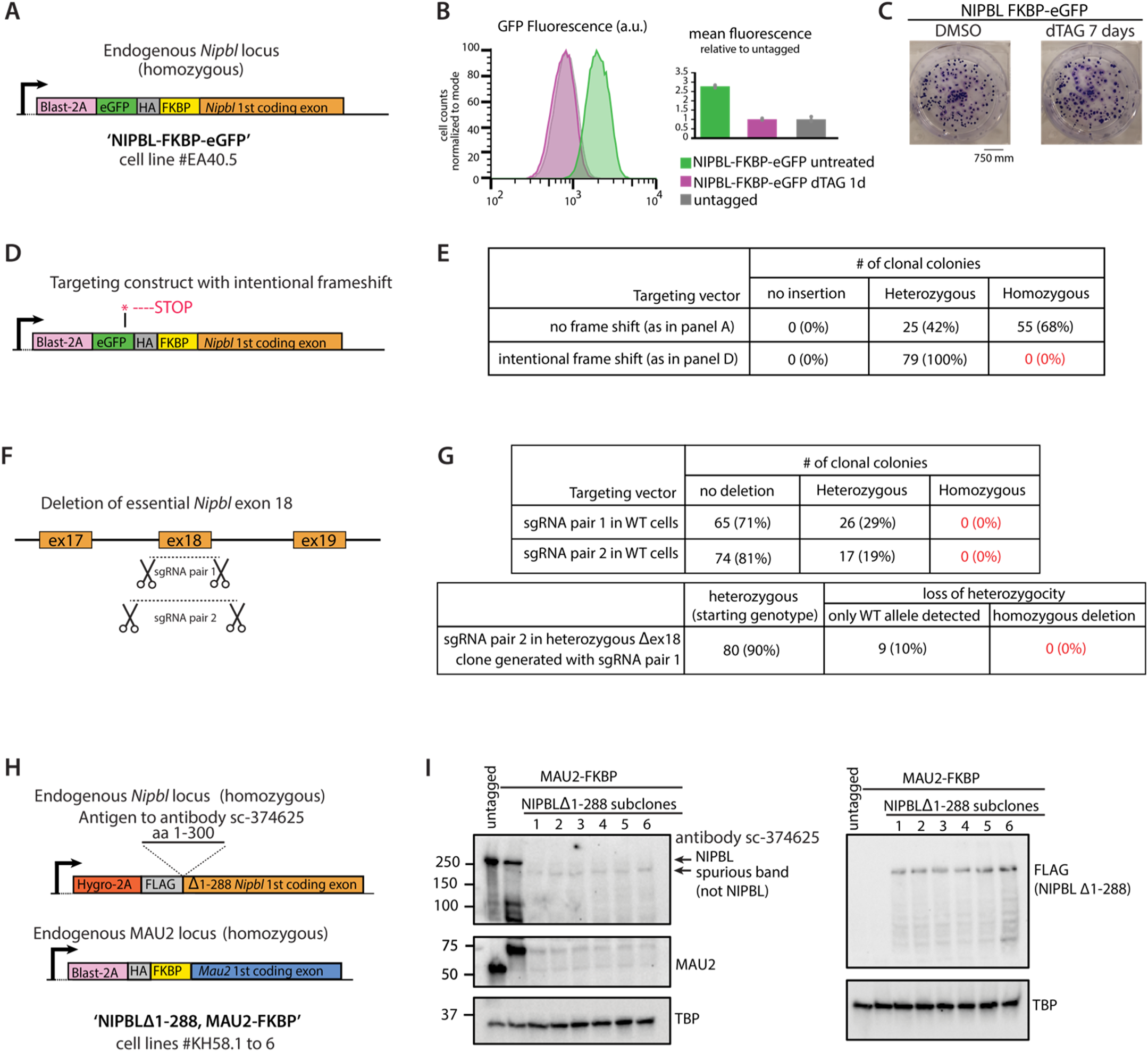
ESCs with leftover levels of NIPBL below 10% still proliferate, yet NIPBL remains essential. **(A)** NIPBL-FKBP-eGFP cell line schematic to enable precise quantification of NIPBL depletion using eGFP autofluorescence. **(B)** eGFP line flow cytometry indicates near-complete depletion of the tagged NIPBL **(C)** Crystal violet staining showing that this cell line with undetectable eGFP-NIPBL retains clonogenicity after depletion. **(D)** Schematic to introduce a frameshift in the NIPBL-degron targeting vector. **(E)** No homozygous frameshifted clones were obtained, indicating that the N-terminal NIPBL targeting vector tags all essential isoforms of NIPBL. **(F)** Schematic to delete essential exon 18, as described in Schwarzer et al. 2017 (*13*). **(G)** No homozygous exon 18 clones were obtained, indicating that NIPBL is essential in mouse embryonic stem cells. **(H)** Deleting the sc-374625 antibody recognition sequence in NIPBL and introducing a FLAG-tag in the background of MAU2-FKBP. **(I)** Westerns for NIPBL and FLAG after deletion of antibody recognition sequence indicating the existence of a spurious band that is not NIPBL with the sc-374625.

**Figure S4.**
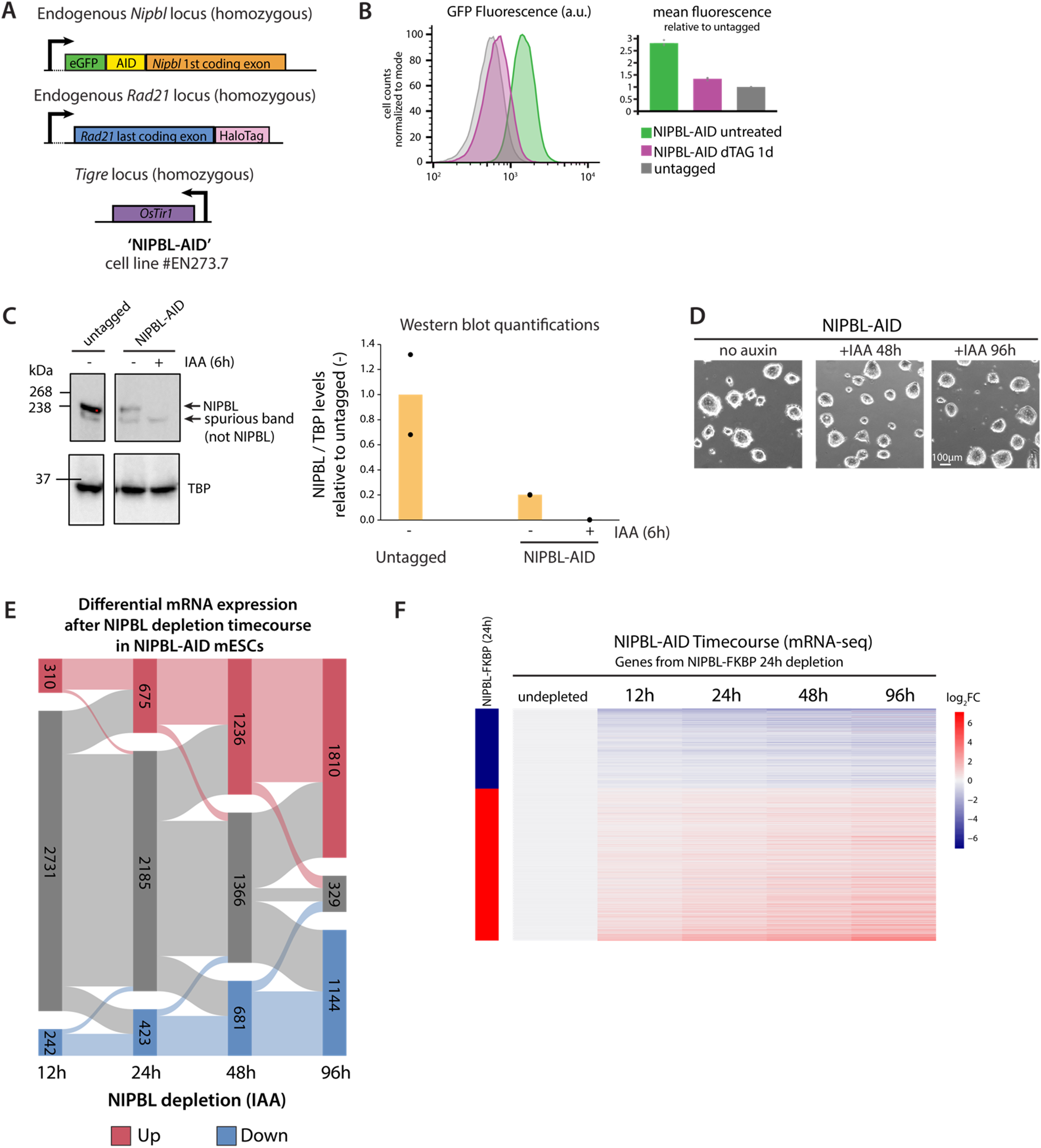
Supporting RNA-seq analyses NIPBL-AID ESCs. **(A)** Schematic of NIPBL-AID cell line (#EN273.7) - see Methods**. (B)** eGFP flow cytometry indicating depletion of the tagged NIPBL with the auxin-inducible degron system. **(C)** Western blot with the sc-374625 NIPBL antibody indicating minimal residual NIPBL remaining after degradation. Note the leakiness of the NIPBL-AID degron line. **(D)** NIPBL-AID depleted ESCs retain vitality despite nearly undetectable levels of NIPBL by Western blot **(E)** DEGs after 12h, 24h, 48h, and 96h of auxin (IAA)-induced depletion in NIPBL-AID ESCs highlighting the gradual increase of DEG number and their temporal trajectories. (FDR < 0.05 and FC ≥ 1.5 or ≤ −1.5). n = 3 replicates for all samples, except n = 2 for 12h depletion. **(F)** Heatmap of log_2_(FC) during NIPBL-AID depletion time course RNA-seq relative to untreated for all significantly differentially expressed genes in NIPBL-FKBP 24h depletion. Note how genes dysregulated in NIPBL-FKBP ESCs show a similar trend in NIPBL-AID ESCs.

**Figure S5.**
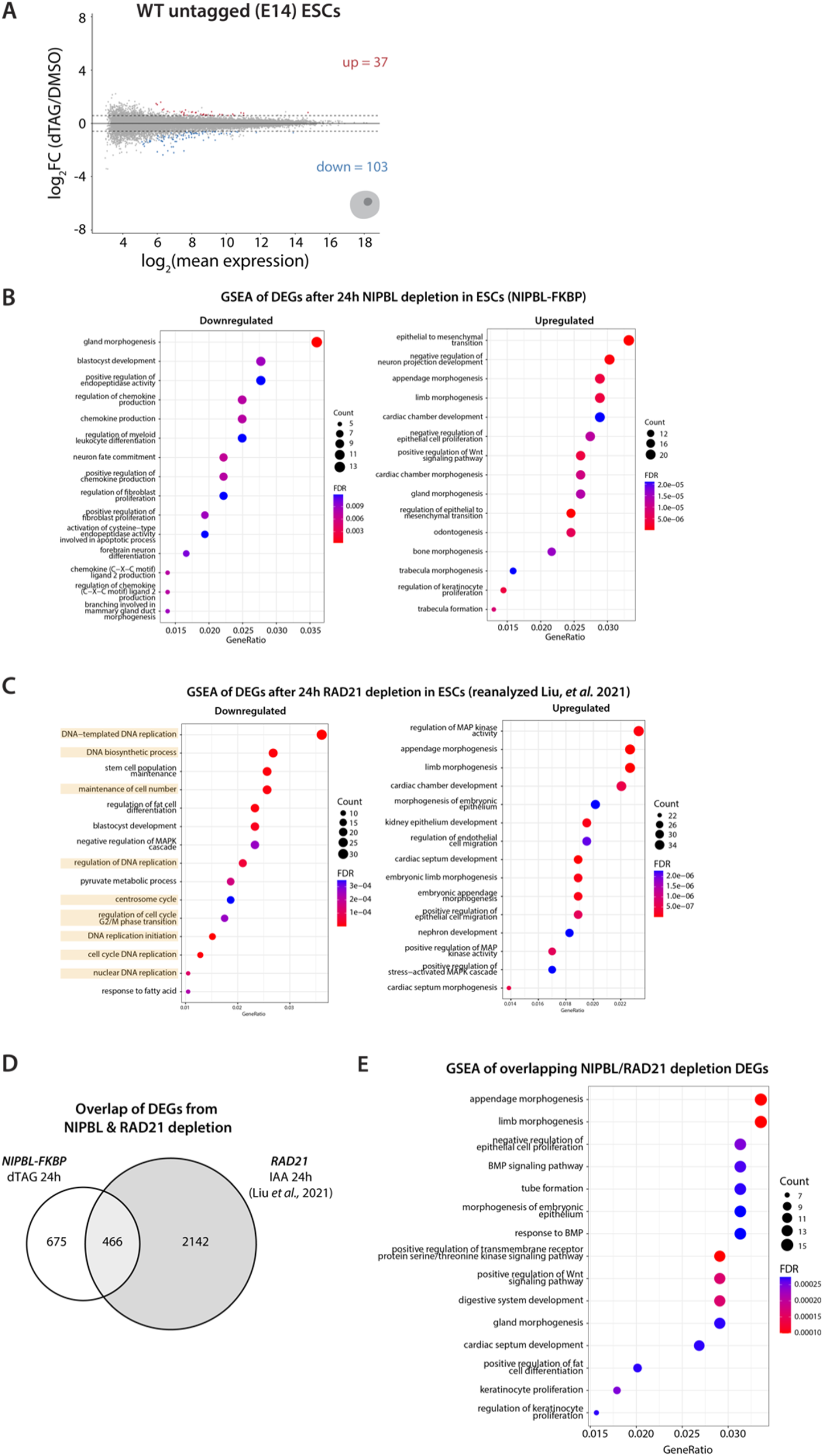
Supporting RNA-seq analyses comparing NIPBL- and RAD21-degron. **(A)** Very few differentially expressed genes (DEGs) are detected after dTAG treatment vs. DMSO in WT untagged (E14) ESCs. (FDR < 0.05 and FC ≥ 1.5 or ≤ −1.5, n = 3). **(B)** Gene set enrichment analysis (GSEA) of DEGs from RNA-seq in NIPBL-FKBP ESCs after 24h dTAG treatment, separated by up- and downregulated genes. Note the absence of terms related to DNA replication/mitosis. **(C)** GSEA of DEGs from published (*8*) and reanalyzed RNA-seq in RAD21-AID ESCs after 24h auxin (IAA) treatment. Note the abundance of terms related to DNA replication/mitosis, highlighted. **(D)** Overlap between DEGs in NIPBL-FKBP (this study) and RAD21-AID (from (*8*)) ESC depletion datasets. Note studies were conducted in different labs and RAD21-AID data were reanalyzed. **(E)** GSEA of DEGs that overlap between NIPBL-FKBP and RAD21-AID ESC depletion datasets, highlighting developmentally relevant rather than cell cycle-related terms.

**Figure S6.**
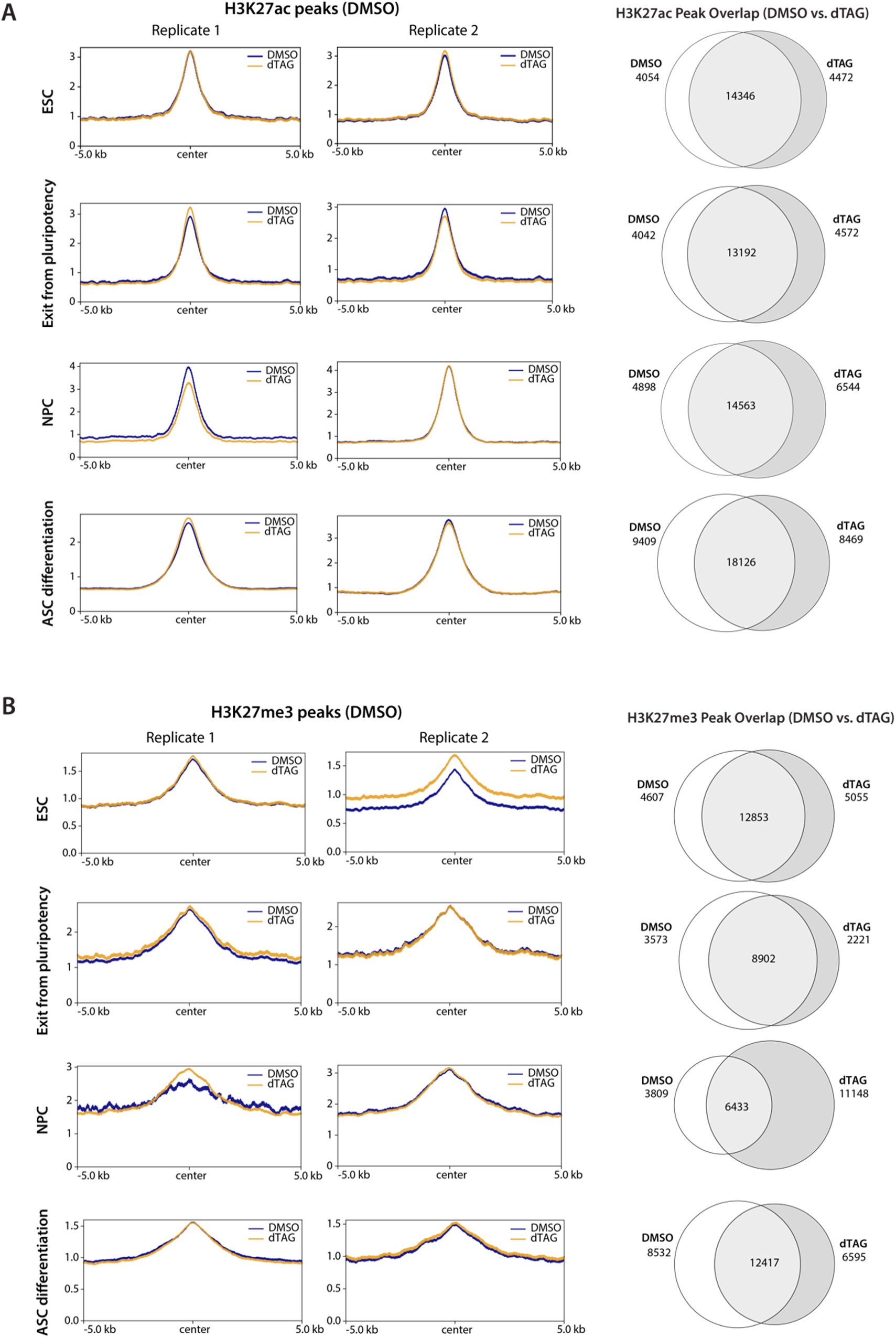
NIPBL depletion does not globally affect H3K27ac or H3K27me3 levels. **(A)** H3K27ac CUT&Tag profiles (left) and DMSO/dTAG peak overlap (right) around peaks detected in NIPBL-FKBP undepleted (DMSO) cells. NIPBL was depleted with dTAG for 24h in ESCs and NPCs, and 48h in ESC exit from pluripotency and ASC differentiations. H3K27ac signal is largely unaffected by NIPBL depletion, indicating that enhancers must retain their intrinsic regulatory potential. **(B)** H3K27me3 CUT&Tag profiles (left) and DMSO/dTAG peak overlap (right) around peaks detected in NIPBL-FKBP undepleted (DMSO) cells. NIPBL was depleted with dTAG for 24h in ESCs and NPCs, and 48h in ESC exit from pluripotency and ASC differentiations. H3K27me3 appears largely unaffected by NIPBL depletion. Note the overall lower signal in ESC DMSO replicate 2, which we attribute to inter-replicate technical variability.

**Figure S7.**
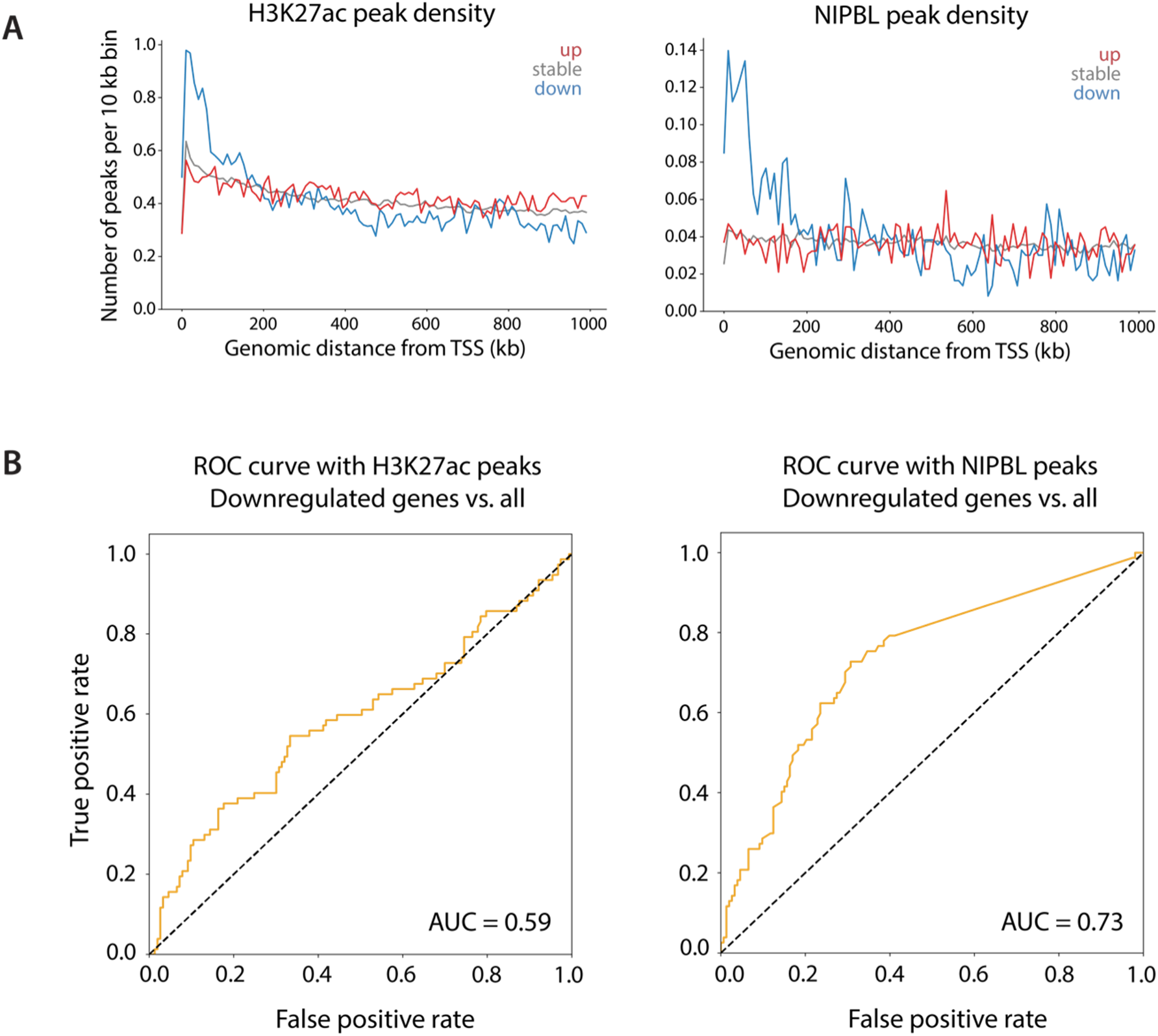
Downregulated genes in ESCs tend to have more putative enhancers nearby. **(A)** The genomic neighborhood of downregulated genes contains more H3K27ac CUT&Tag peaks (left) and NIPBL ChIP-seq peaks (right) within 100-200 kb of the TSS, indicating a higher local density of putative enhancers. Note NIPBL peaks largely overlap distal H3K27ac but are not found at promoters. Left panel reproduced from Fig. 1 for comparison. **(B)** Area under the receiver-operating characteristic curve (AUC ROC) indicating that NIPBL peak density is predictive of downregulation upon NIPBL depletion (right). In contrast, H3K27ac on its own is not a strong predictor, possibly due to the presence of peaks at some gene promoters (left).

**Figure S8.**
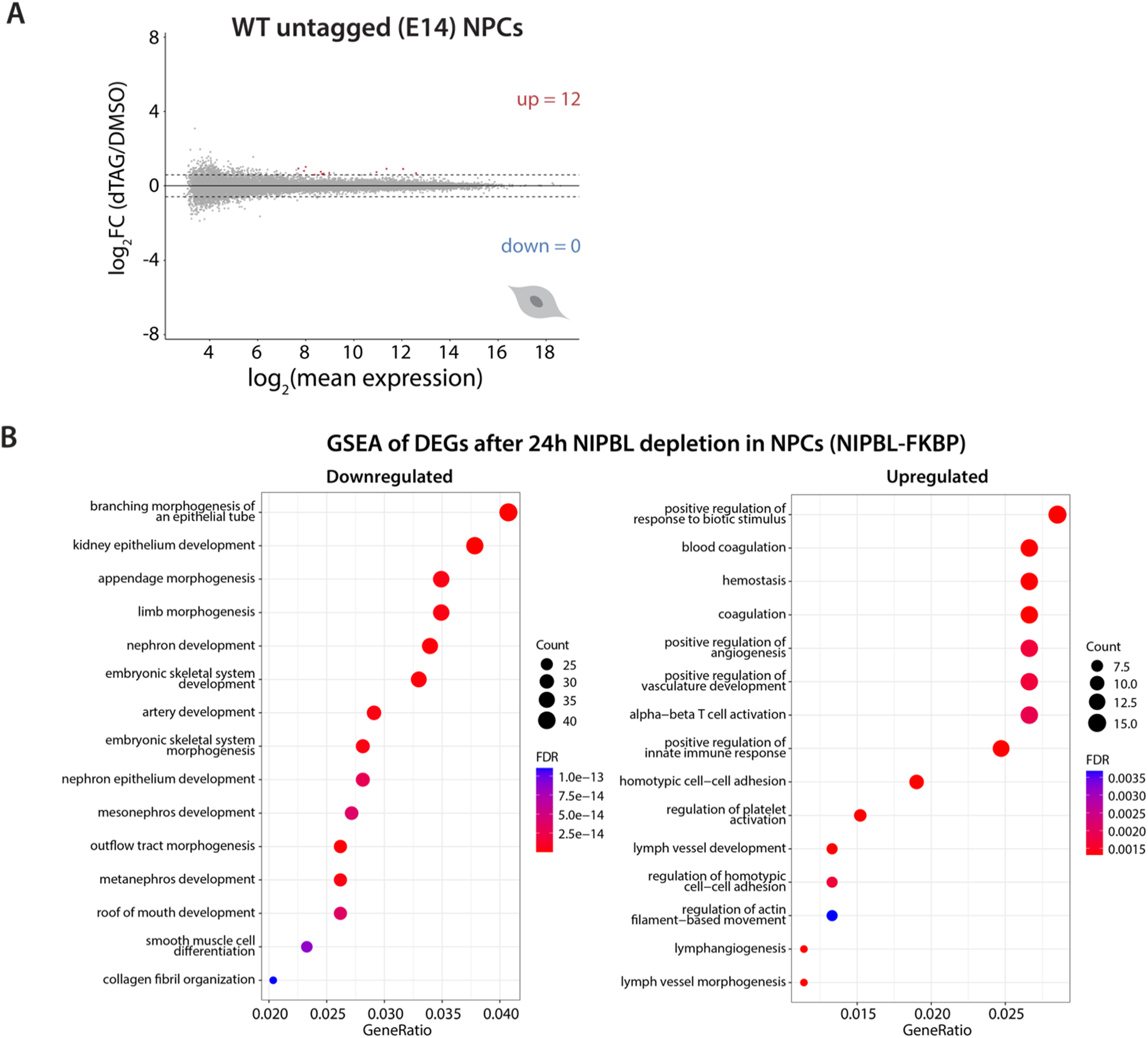
Supporting RNA-seq analyses in control and NIPBL-depleted NPCs. **(A)** Very few DEGs are detected after dTAG treatment vs. DMSO in untagged (WT) E14 untagged NPCs (FDR < 0.05 and FC ≥ 1.5 or ≤ −1.5). n = 3 replicates. **(B)** GSEA of DEGs from RNA-seq in NIPBL-FKBP NPCs after 24h dTAG treatment. Note the absence of cell cycle-related terms and the abundance of developmental terms.

**Figure S9.**
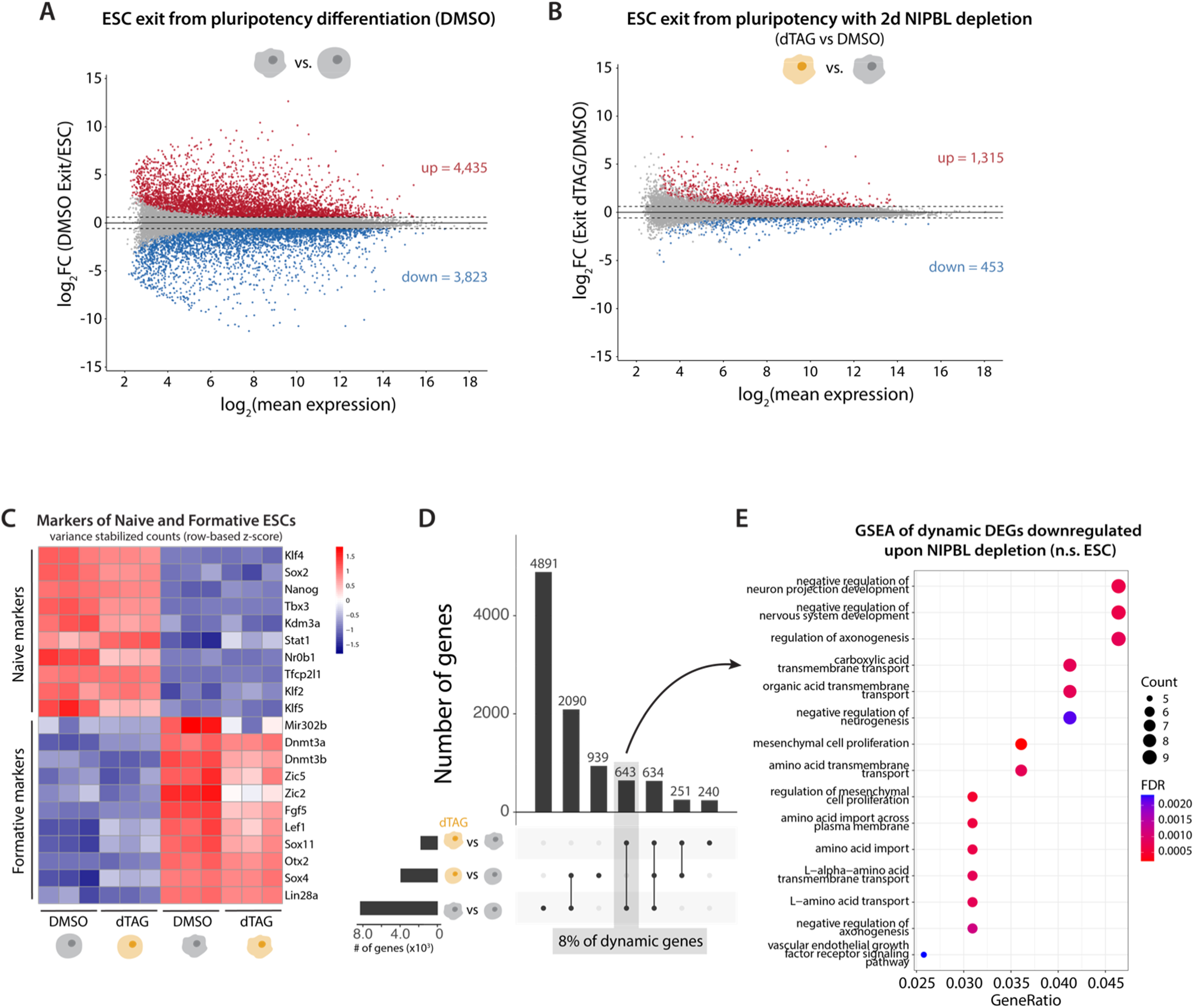
Supporting RNA-seq analyses in ESCs exiting pluripotency after NIPBL depletion. **(A)** Dynamic genes in the exit from pluripotency, defined as the DEGs in the undepleted (DMSO) differentiation vs. ESCs (DMSO) (FDR < 0.05 and FC ≥ 1.5 or ≤ −1.5). n = 3 replicates. **(B)** DEGs from RNA-seq comparing NIPBL depleted (+dTAG 2 days) vs. undepleted control during the exit from pluripotency (FDR < 0.05 and FC ≥ 1.5 or ≤ −1.5). n = 3 replicates. **(C)** Row based z-score variance stabilized counts for markers of naïve and formative pluripotency. Note the largely normal downregulation of pluripotency markers and the upregulation of differentiation markers, despite 2 days of NIPBL depletion. Each column is an individual replicate. **(D)** Upset plot highlighted the various categories of DEGs when analyzing NIPBL-depleted cells during differentiation, the shaded category highlights genes that are normally dynamic during normal differentiation but are called differentially expressed in NIPBL-depleted differentiated cells (and not dysregulated in undifferentiated ESCs). **(E)** GSEA of gene set highlighted in D.

**Figure S10.**
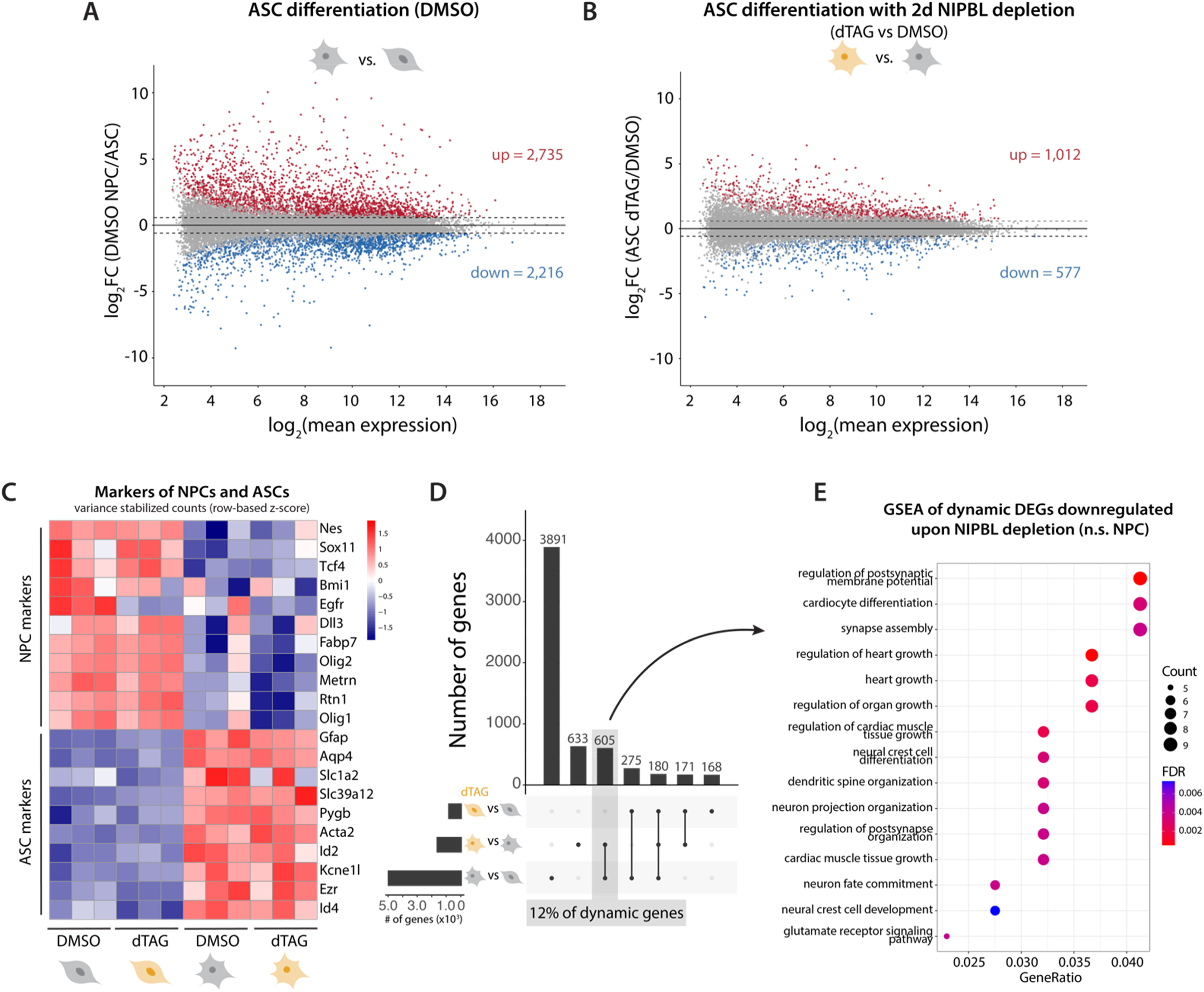
Supporting RNA-seq analyses in NPCs differentiating into ASCs after NIPBL depletion. **(A)** Dynamic genes in the differentiation of NPCs into ASCs, defined as the DEGs in the undepleted (DMSO) differentiation vs. NPCs (DMSO) (FDR < 0.05 and FC ≥ 1.5 or ≤ −1.5). n = 3 replicates. **(B)** DEGs from RNA-seq comparing NIPBL depleted (+dTAG 2 days) vs. undepleted control during the differentiation of NPCs into ASCs (FDR < 0.05 and FC ≥ 1.5 or ≤ −1.5). n = 3 replicates. **(C)** Row based z-score variance stabilized counts for markers of NPCs and ASCs. Note the largely normal downregulation of NPC markers and the upregulation of ASC markers despite 2 days of NIPBL depletion. Each column is an individual replicate. **(D)** Upset plot highlighted the various categories of DEGs when analyzing NIPBL-depleted cells during differentiation, the shaded category highlights genes that are normally dynamic during undepleted differentiation but are called differentially expressed in NIPBL-depleted differentiated cells (and not dysregulated in undifferentiated NPCs). **(E)** GSEA of gene set highlighted in D.

**Figure S11.**
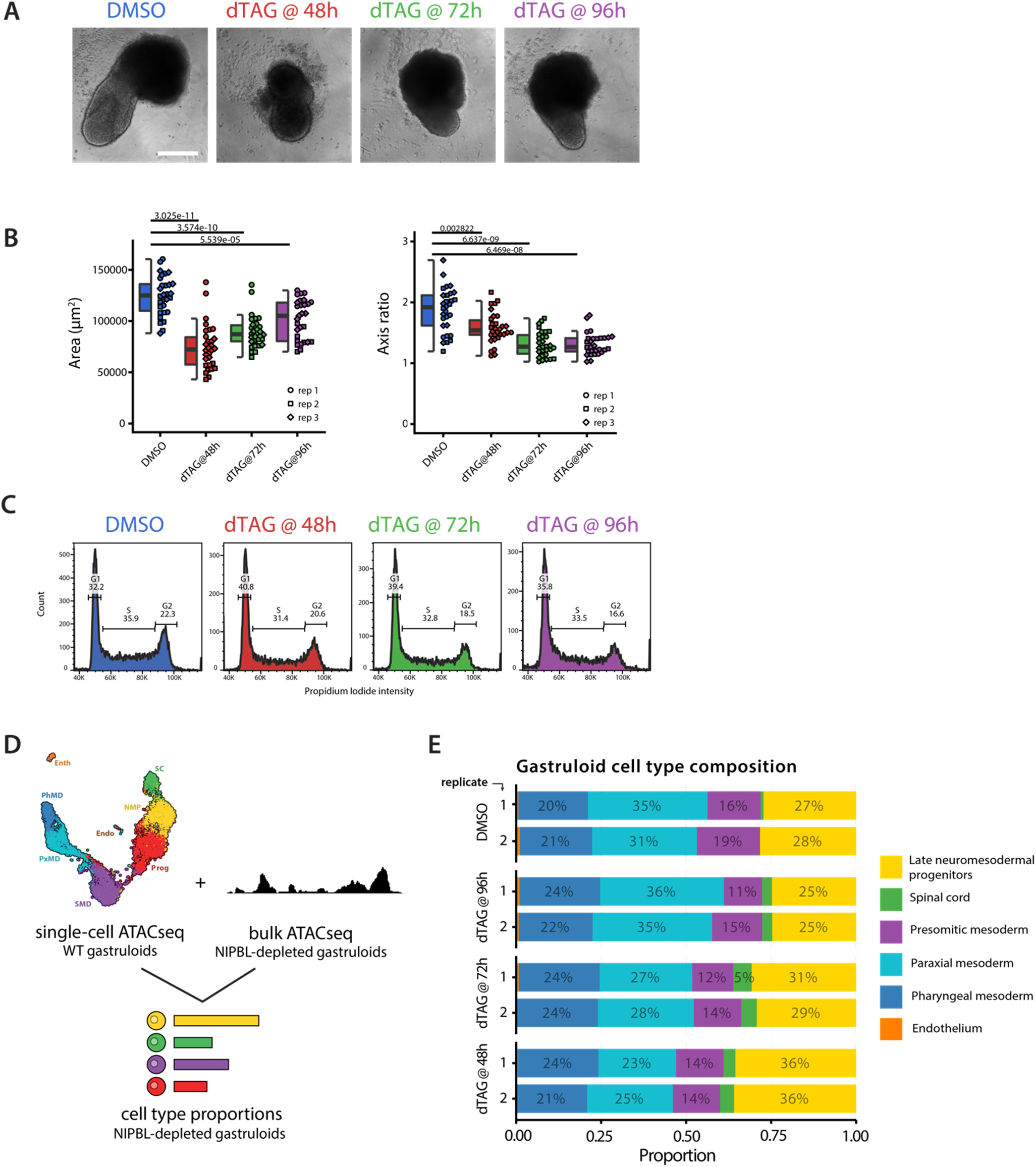
Supporting analyses for gastruloid differentiations. **(A)** Brightfield microscopy of control and NIPBL-depleted gastruloids 120h post-aggregation. dTAG was added at either 48, 72, or 96h post-aggregation as described in Fig. 2. **(B)** Quantification of gastruloid area (left) and axis ratio (right) for control and NIPBL-depleted gastruloids, highlighting disrupted elongation particularly when depletion is started earlier in differentiation (n = 3 independent experiments). **(C)** Cell cycle analyses do not show major defects in NIPBL-depleted gastruloids. **(D)** An available scATAC-seq timecourse of gastruloid development (*27*) was used to deconvolve bulk ATAC-seq (as in (*28*)) in control and NIPBL-depleted gastruloids, allowing the resolution of cell type composition. **(E)** Cell type composition of gastruloids (n = 2).

**Figure S12.**
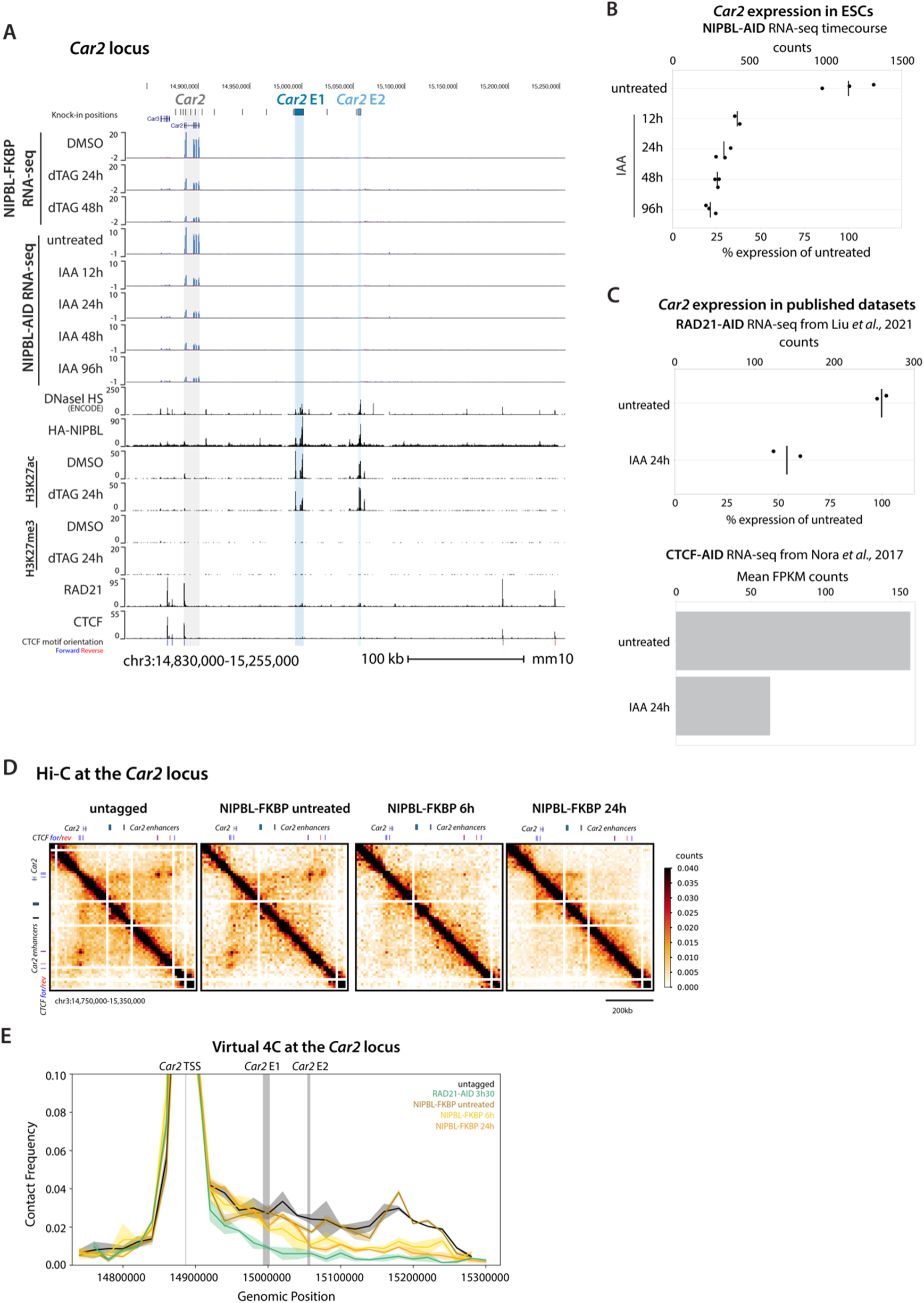
Supporting analyses of the *Car2* locus upon NIPBL depletion in ESCs. **(A)** UCSC browser view of the *Car2* locus displaying related transcriptomic data (NIPBL-FKBP RNA-seq and NIPBL-AID RNA-seq), DNaseI hypersensitivity (ENCODE), CUT&Tag for H3K27ac and H3K27me3 (NIPBL-FKBP control and dTAG 24h depletion), and ChIP-seq for NIPBL (HA tag), cohesin (RAD21), CTCF with motif orientations, and enhancer knock-in positions. Shading highlights the *Car2* gene in gray, *Car2* enhancer 1 (CE1) in dark blue, and *Car2* enhancer 2 (CE2) in light blue. **(B)** *Car2* mRNA expression decreases in NIPBL-AID ESCs throughout a 96h depletion timecourse, in agreement with NIPBL-FKBP depletion RNA-seq. Counts are size factor normalized. Each point is an individual replicate, and notches indicate averages (n = 3). **(C)** *Car2* mRNA expression decreases in RAD21-AID ESCs after 24h depletion (*8*), in agreement with NIPBL RNA-seq. Counts are size factor normalized. **(D)** Hi-C contact maps of the *Car2* locus upon NIPBL depletion, bin size = 20 kb. **(E)** Virtual 4C across the *Car2* locus demonstrating diminished long-range interactions and reduced contact frequency between *Car2* and its enhancers (CE1 and CE2) when loop extrusion is disrupted. Solid line indicates the mean of two replicates, bin size = 20 kb.

**Figure S13.**
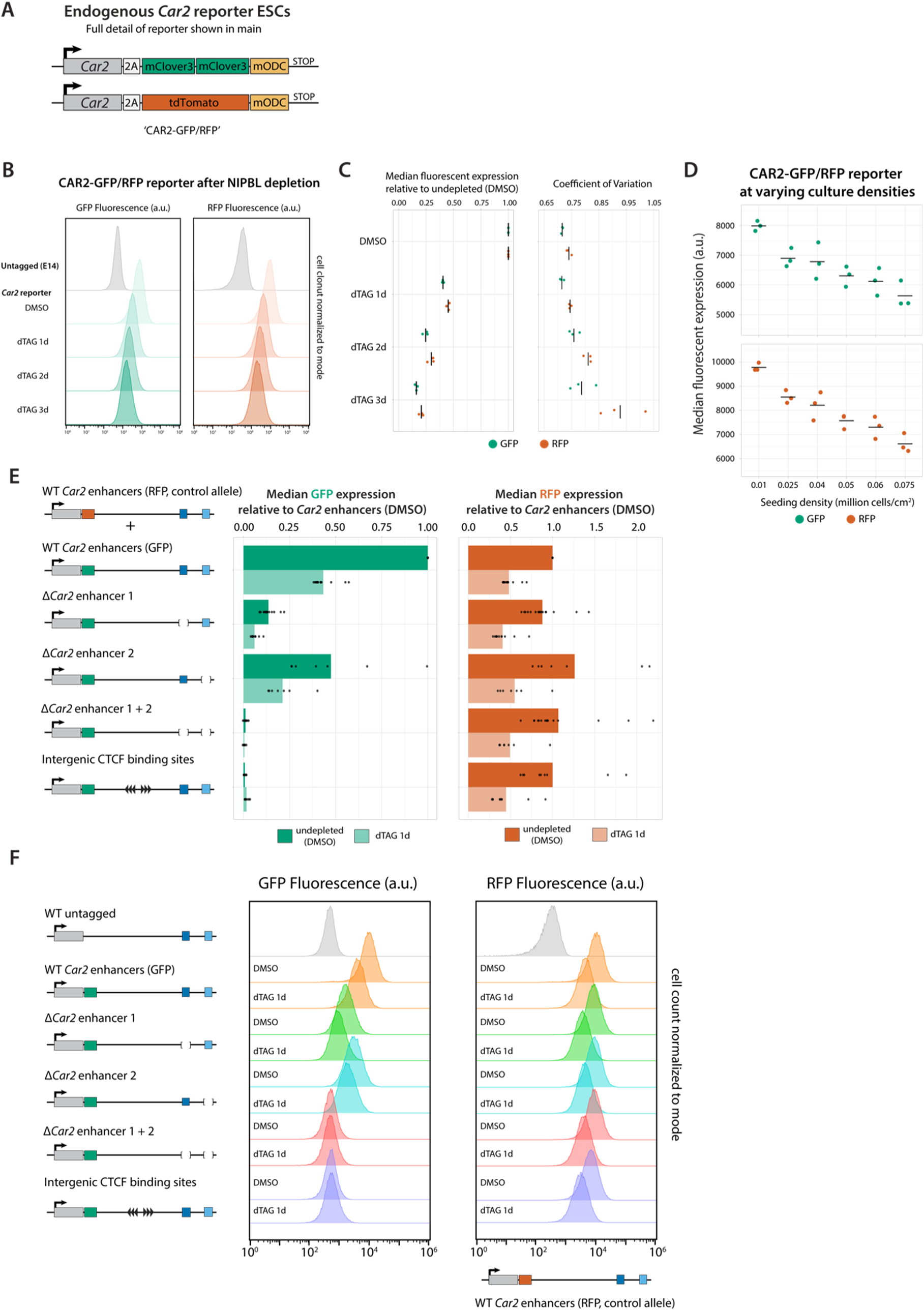
Complementary characterization Car2-reporter ESCs. **(A)** Details of the *Car2* fluorescent reporter ESCs from Fig. 3C. The endogenous *Car2* gene was edited to express a 2A-2XmClover3 (GFP) cassette from one allele and a 2A-tdTomato (RFP) cassette from the other allele on the C-terminus of CAR2. Both fluorescent reporters include the mammalian ornithine decarboxylase (mODC) destabilization domain to reduce their half-life. **(B)** Representative flow cytometry histograms of GFP (left) and RFP (right) fluorescent expression from the *Car2* reporter during a 3-day NIPBL depletion time course. Note that cells were cultured precisely at the same confluency. **(C)** Flow cytometry of *Car2* reporter ESCs over a 3-day time course of NIPBL depletion recapitulates RNA-seq results without major changes in cell-to-cell variation. Median fluorescent expression is shown relative to undepleted control (left) and the Coefficient of Variation between single cells (CV) for each allele (right) (n = 3 flows). **(D)** *Car2* expression is sensitive to cell culture density. Median fluorescent expression is shown for both GFP and RFP alleles. Each point is an individual replicate, and notches indicate averages (n = 3). These observations stress the importance of using an inter-allelic reporter strategy to avoid confounding factors that act on both alleles in *trans* (n = 3 flows). **(E)** CE1 and CE2 deletions and separate knock-in of 6X CTCF binding site cassette (3X reverse, 3X forward) between *Car2* and its enhancers. Median fluorescent expression is shown relative to reporter with WT enhancers separated by edited GFP (left) and WT RFP (right) allele. Each point is an individual flow replicate, and bars indicate averages (n ≥ 7 flows from independent cultures). **(F)** Representative flow cytometry histograms of GFP (left) and RFP (right) of enhancer deletion and CTCF binding site knock-in cell lines from E, highlighting the unimodal downregulation upon NIPBL depletion, enhancer knockouts and CTCF knock-in.

**Figure S14.**
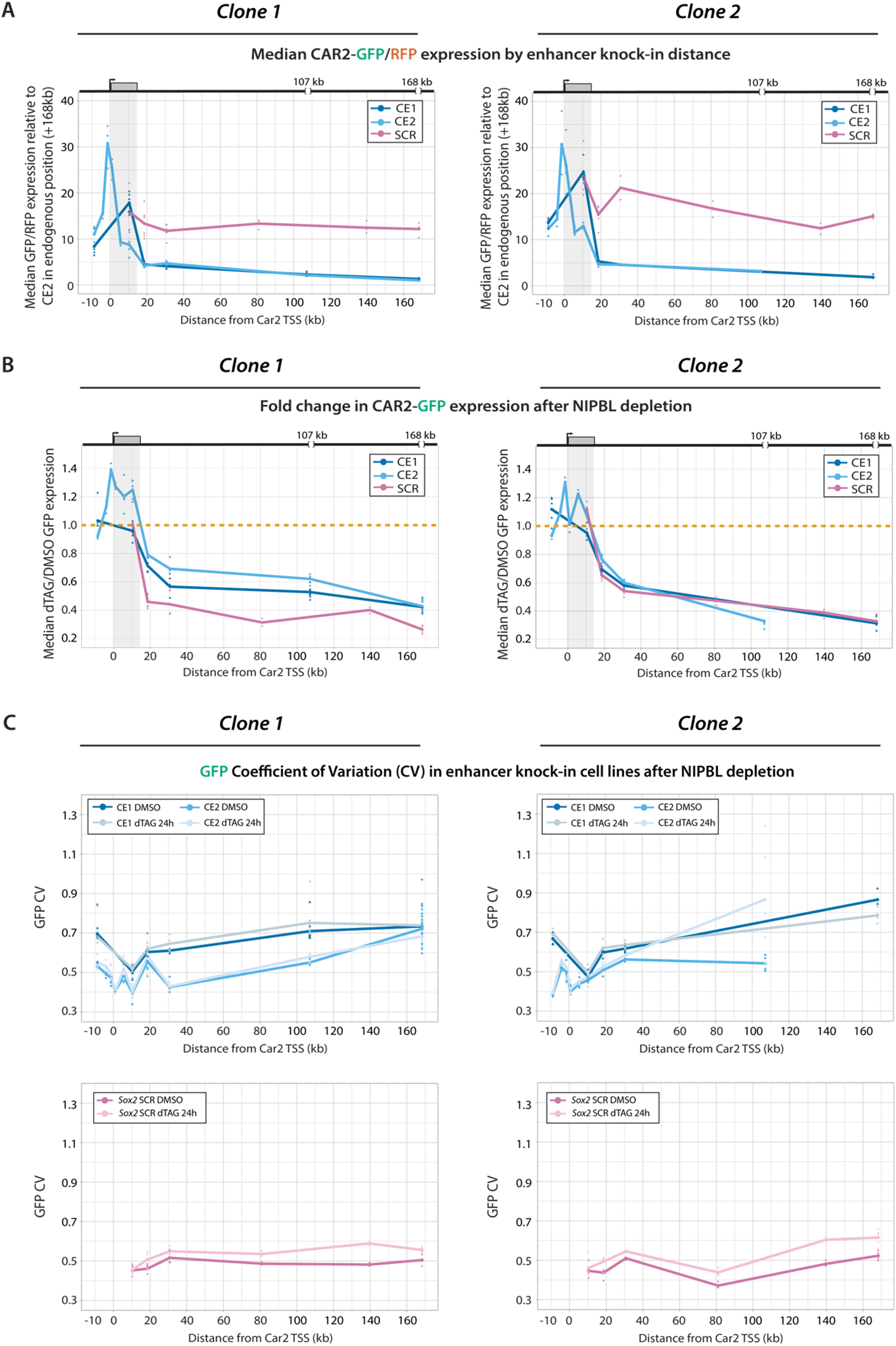
Biological replicates (second targeted clone) for each enhancer knock-in at *Car2*. **(A)** Basal *Car2* expression levels are lower for large genomic distance to the enhancer, for both CE1 and CE2. This is not the case for the SCR, which contains a strong CTCF site in the reverse orientation. Median allelic ratios for enhancer knock-ins across the *Car2* locus are shown relative to a cell line with CE2 in its endogenous location (+168 kb). Shading highlights the *Car2* gene body. Gaps indicate the positions of endogenous CE1 (+107 kb) and CE2 (+168 kb) in WT. Data is shown for two clonal cell line for each enhancer reinsertion. Note the high concordance between clone 1 (left and Figure 3G) and clone 2 (right). Each point is an individual flow replicate, and line connects averages (n ≥ 3 flows from independent cultures). **(B)** Loop extrusion is no longer required for expression of *Car2* when an enhancer is within 11 kb of the TSS. Fold change in median CAR2-GFP expression after NIPBL depletion (dTAG 24h/DMSO). Orange line indicates no change between conditions (FC = 1.0). Note the high concordance of the two clonal lines analyzed, shown for clone 1 (left and Figure 3H) and clone 2 (right). (n ≥ 3 flows). **(C)** Cell-to-cell variation fluctuates depending on *Car2* enhancer position, increasing some with distance, but is stable regardless of position for the SCR. Variation does not change after NIPBL depletion (24h). In general, cell lines with higher expression display lower coefficient of variation between single cells (n ≥ 3 flows).

**Figure S15.**
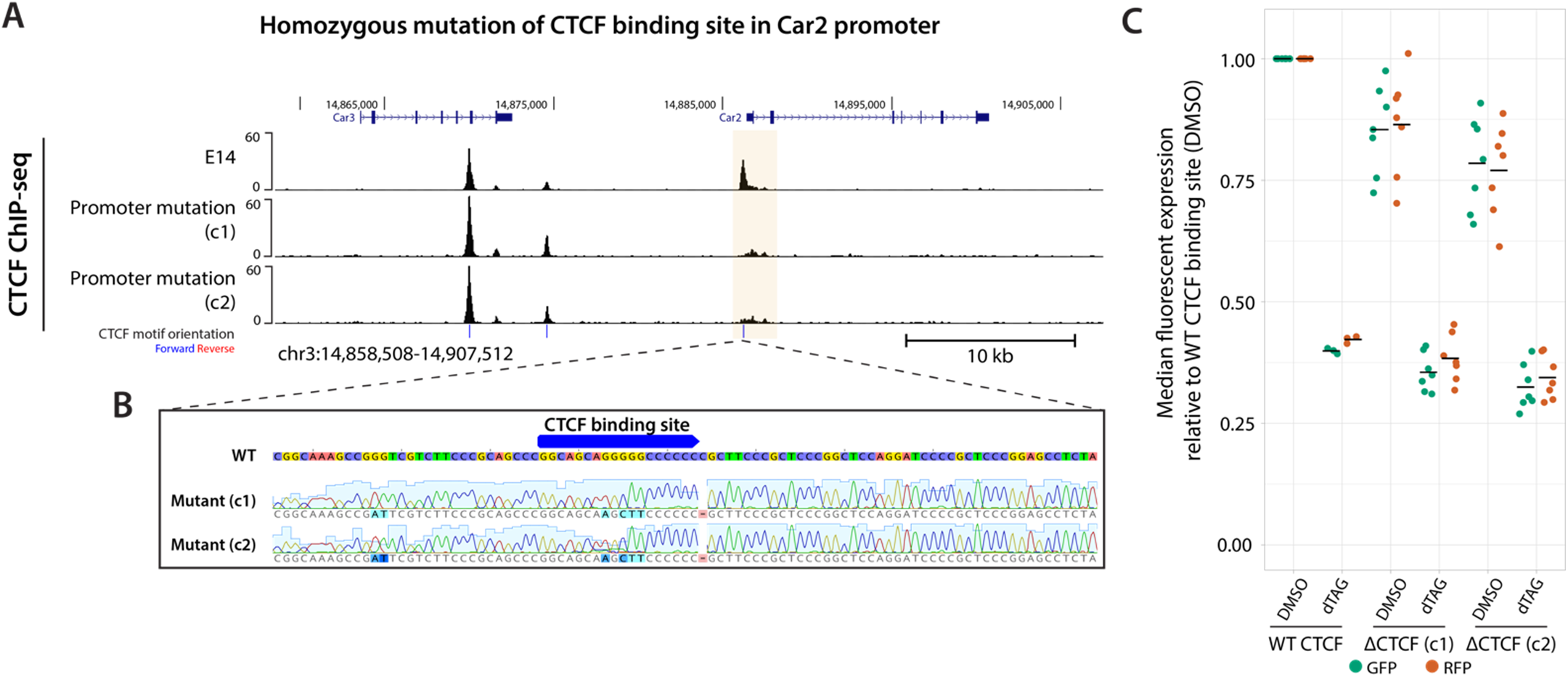
Abrogating CTCF binding at the *Car2* TSS does not alter the sensitivity of the *Car2* locus on NIPBL. **(A)** CTCF ChIP-seq highlighting binding at the Car2 TSS. **(B)** Two clonal cell lines with targeted removal of the CTCF binding site (CBS) using a knock-in vector were generated (in NIPBL-FKBP, CAR2-GFP/RFP ESCs). Sanger sequencing of CBS mutation is shown. **(C)** ΔCTCF clones exhibit a small reduction in basal Car2 expression but remain equally sensitive to the depletion of NIPBL Median fluorescent expression relative to WT DMSO in CBS mutation lines.

**Figure S16.**
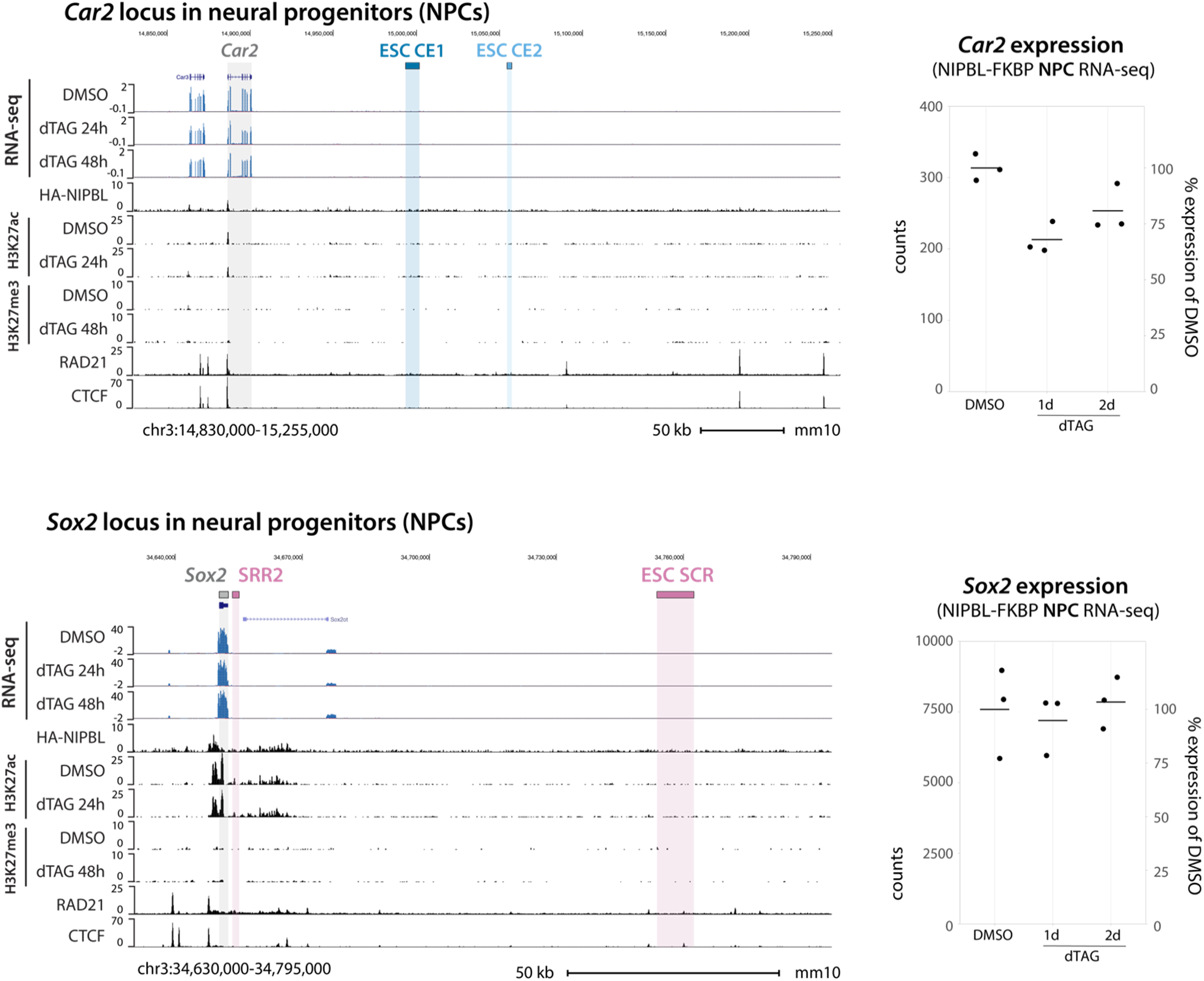
Epigenomic landscapes of *Car2* and *Sox2* in NPCs. Browser snapshot of the *Car2* locus in NPCs (top), highlighting the decommissioning of CE1 and CE2. *Car2* expression is ∼10 times lower than in ESCs (Fig. 3) and no longer significantly dysregulated. Similarly, at the *Sox2* locus in NPCs (bottom), the SCR is decommissioned and promoter-proximal H3K27ac clusters arise. *Sox2* expression is only ∼1.5 times lower than in ESCs (Fig. 4). The mild reliance of *Sox2* on NIPBL observed in ESCs is absent in NPCs, in line with the appearance of promoter-proximal H3K27ac clusters and decommissioning of the distal SCR. RNA-seq counts are size factor normalized, n = 3 replicates.

**Figure S17.**
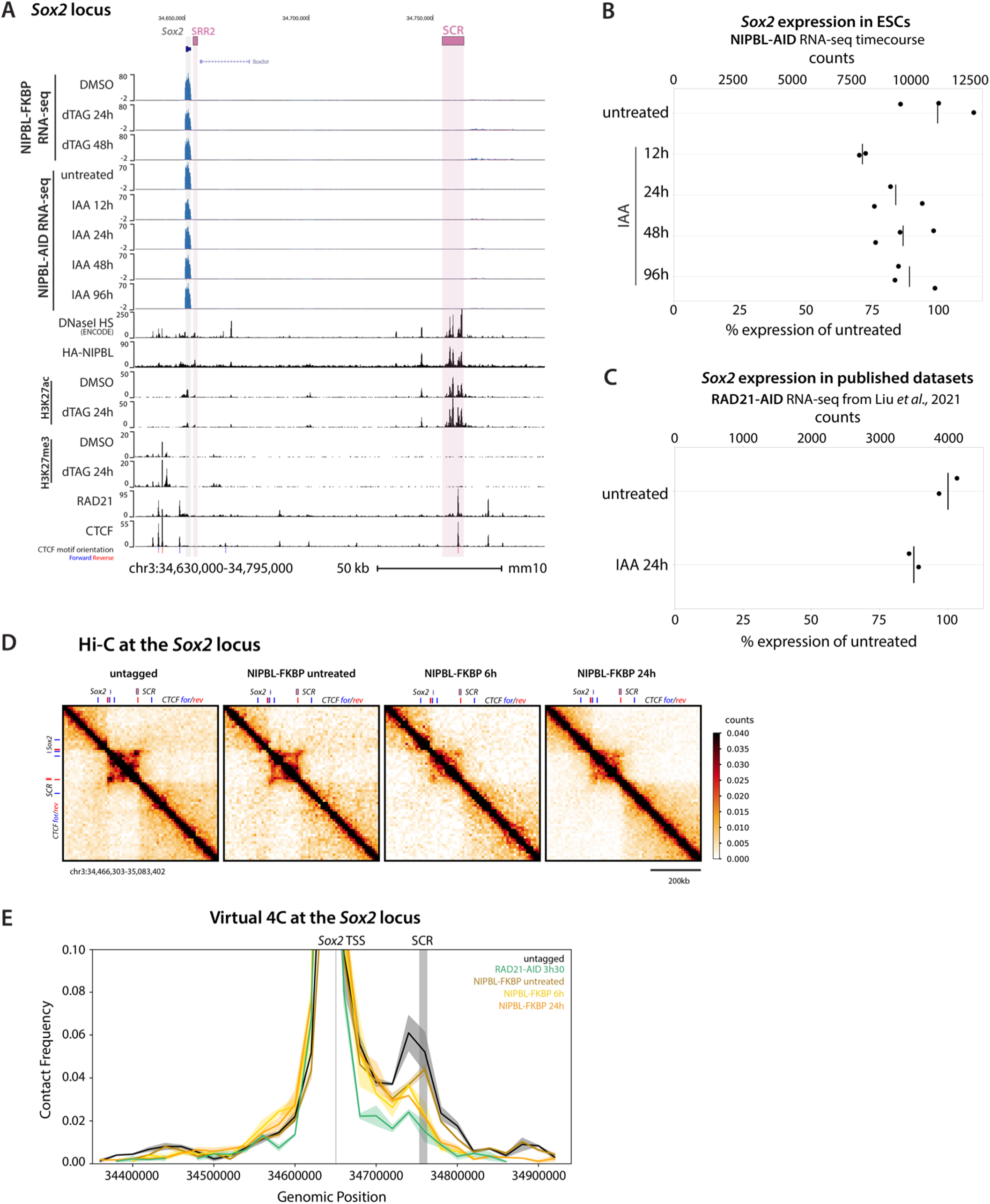
Supporting analyses of the *Sox2* locus upon NIPBL depletion in ESCs. **(A)** UCSC browser view of the *Sox2* locus displaying related transcriptomic data (NIPBL-FKBP RNA-seq and NIPBL-AID RNA-seq), DNaseI hypersensitivity (ENCODE), CUT&Tag for H3K27ac and H3K27me3 (NIPBL-FKBP control and dTAG 24h depletion), and ChIP-seq for NIPBL (HA tag), cohesin (RAD21), and CTCF with motif orientations. Gray shading highlight *Sox2* and pink shading highlights regulatory elements SRR2 and SCR. **(B)** *Sox2* mRNA expression in NIPBL-AID ESCs over a 96h depletion timecourse. Expression decreases mildly (∼25%) upon initial depletion and remains stable throughout the timecourse, in agreement with NIPBL-FKBP depletion RNA-seq. Counts are size factor normalized (n = 3). **(C)** *Sox2* mRNA expression in RAD21-AID ESCs after 24h depletion (*8*). In agreement with NIPBL depletion RNA-seq, *Sox2* expression only decreases minimally when cohesin is directly targeted. Counts are size factor normalized. **(D)** Hi-C contact maps of the *Sox2* locus upon NIPBL depletion, bin size = 20 kb. **(E)** Virtual 4C across the *Sox2* locus demonstrating diminished long-range interactions and reduced contact frequency between *Sox2* and the SCR when loop extrusion is disrupted. Solid line indicates the mean of two replicates, bin size = 20 kb.

**Figure S18.**
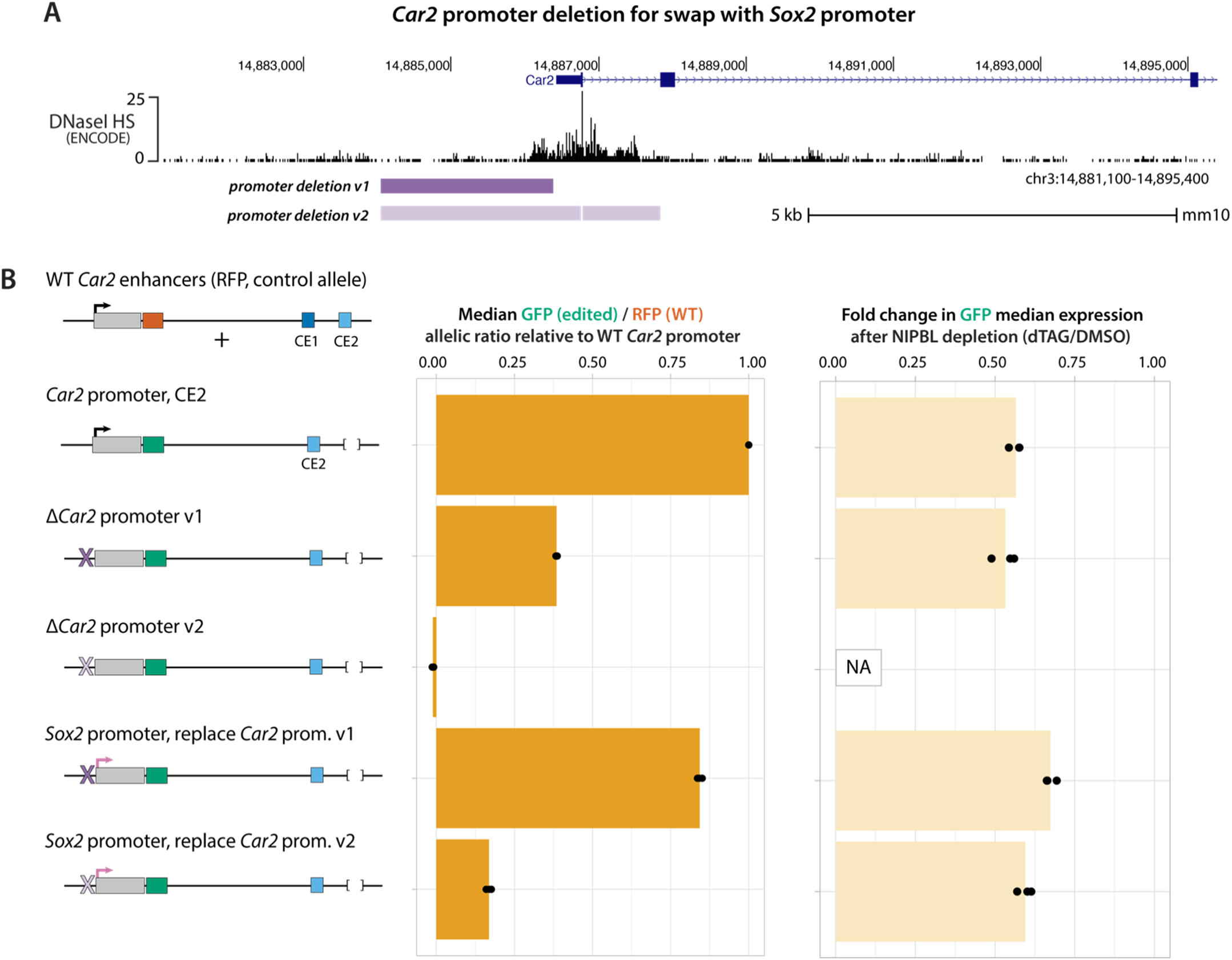
Swapping the *Car2* promoter for the *Sox2* promoter does not alter the sensitivity of the *Car2* locus on NIPBL. **(A)** DNaseI hypersensitvity (ENCODE) around the *Car2* TSS, showcasing dispersed transcription factor binding spreading into the first intron. Location of the two deletions attempted to remove the *Car2* promoter [promoter deletion version (v) 1 and v2]. **(B)** Promoter deletion v1 did not fully abrogate *Car2* expression but promoter deletion v2 did (middle), indicating that regions within the first intron have promoter activity. Knocking in the *Sox2* promoter to rescue *Car2* expression after promoter deletion v1 and v2 restored some expression. However, none of these modifications altered the sensitivity of Car2 to NIPBL depletion (right), indicating that the *Sox2* promoter relies on cohesin loop extrusion in this context (n = 3 flows).

**Figure S19.**
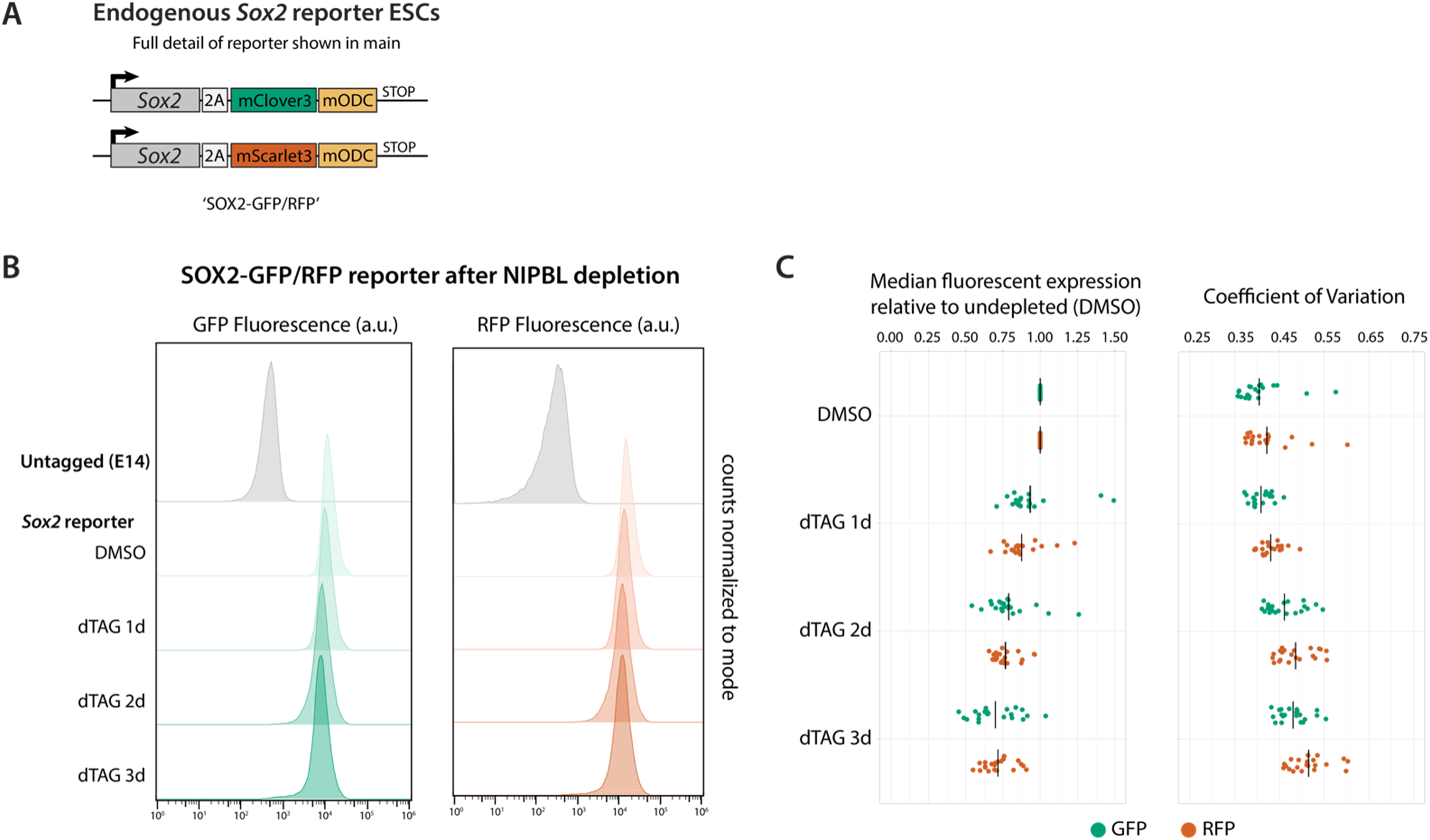
Complementary characterization of *Sox2*-reporter ESCs. **(A)** Details of the *Sox2* fluorescent reporter ESCs from Figure 4E. The endogenous *Sox2* gene was edited to express a 2A-mClover3 (GFP) cassette from one allele and a 2A-mScarlet3 (RFP) cassette from the other allele on the C-terminus of SOX2. Both fluorescent reporters include the mODC destabilization domain to reduce their half-life. **(B)** Representative flow cytometry histograms of GFP (left) and RFP (right) fluorescent expression from the *Sox2* reporter during a 3-day NIPBL depletion time course. **(C)** Flow cytometry of SOX2-GFP/RFP ESCs over a 3-day time course of NIPBL depletion recapitulates the ∼25% decreased observed by RNA-seq (left), and reveals a small increase in cell-to-cell variation (right) (n ≥ 3 flows).

**Figure S20.**
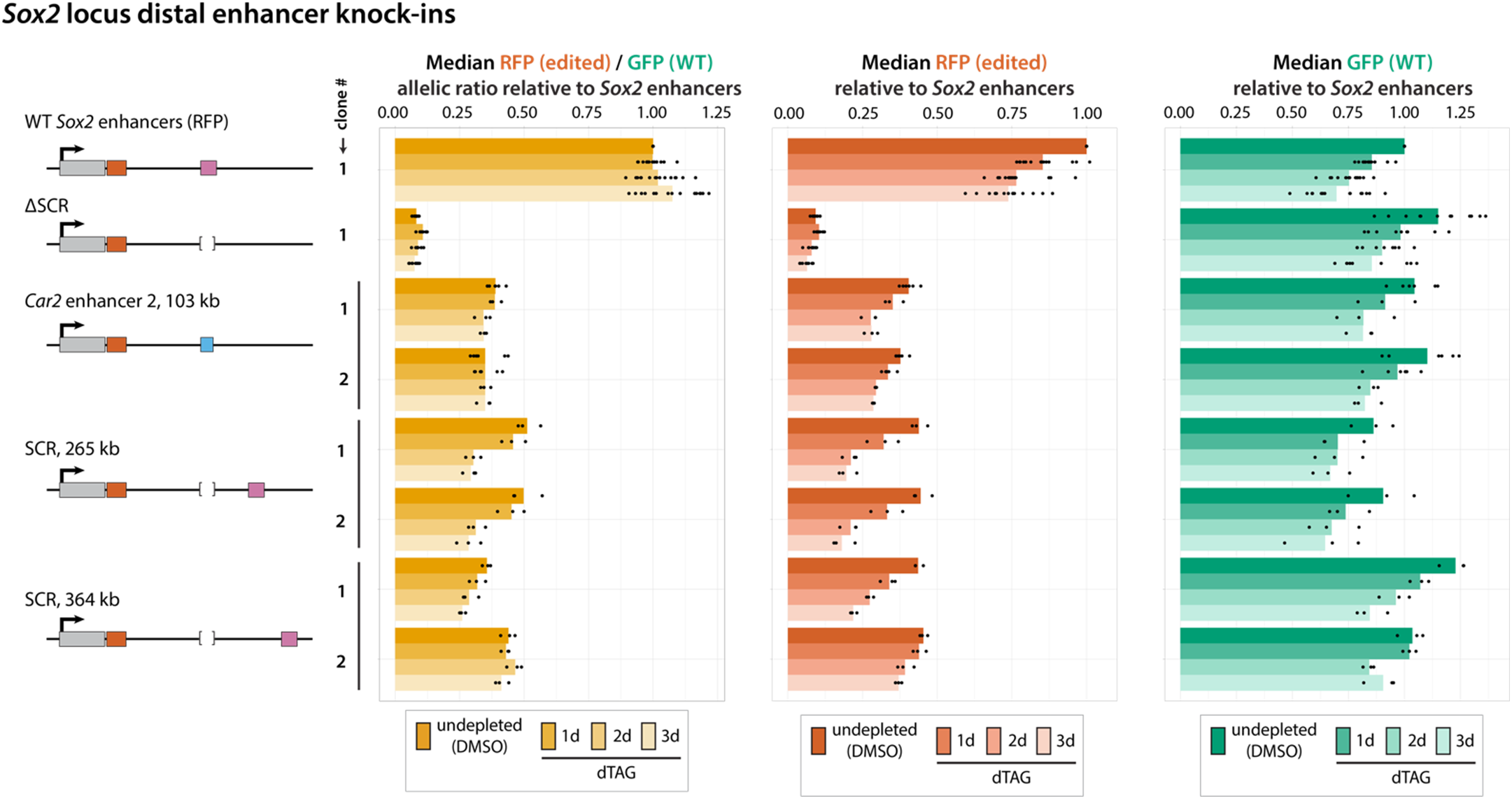
*Sox2* expression remains largely independent of NIPBL when driven by the *Car2* enhancer or remote SCR. Replacing the SCR with the *Car2* enhancer 2 reduces baseline expression but does not alter NIPBL-dependence. Moving the SCR further from the *Sox2* TSS (265 or 364 kb) reduces basal expression and can slightly sensitize *Sox2* to disrupted loop extrusion, especially after several days (n ≥ 3 flows and 2 clones).

**Figure S21.**
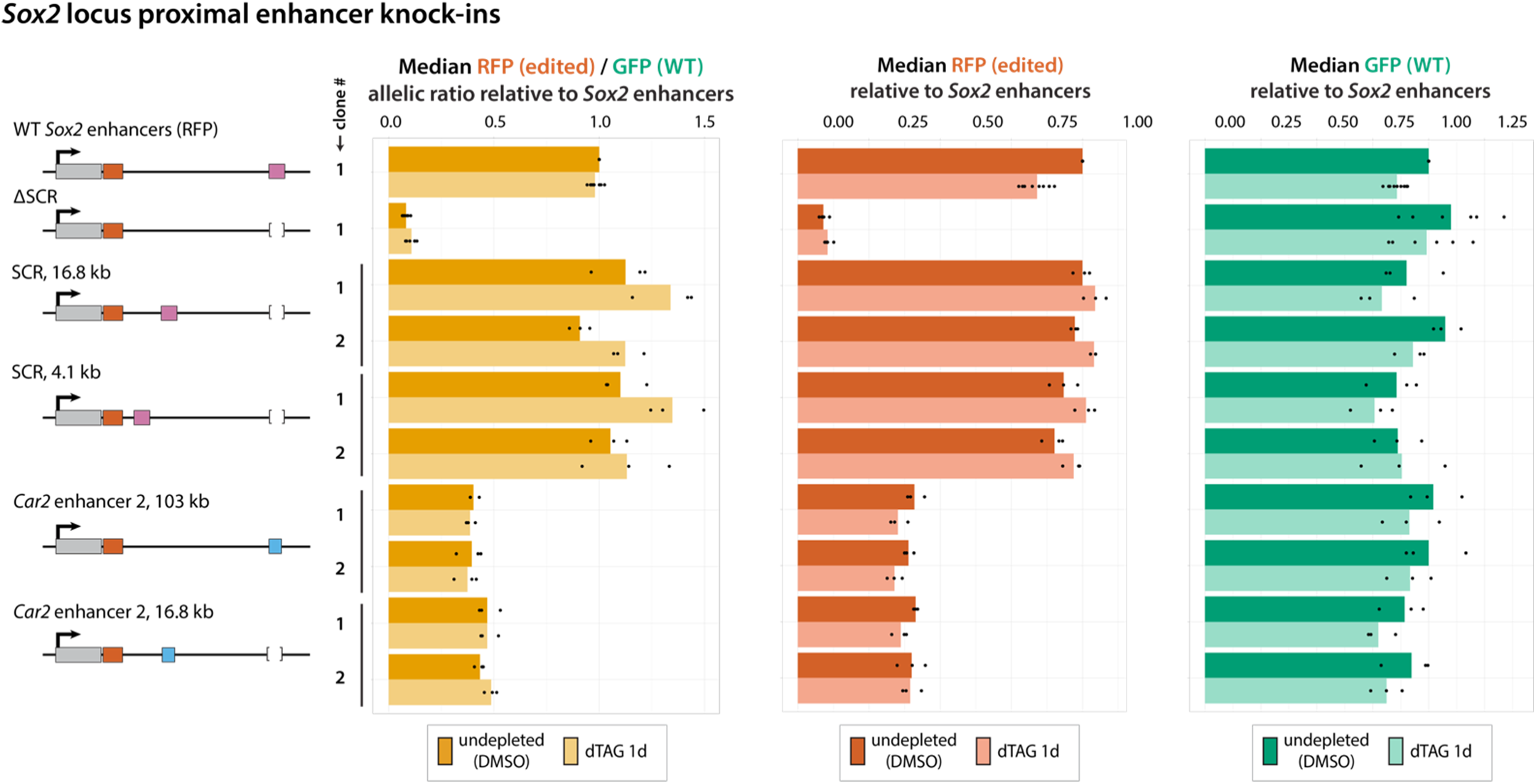
Relocating enhancers proximal to *Sox2* does not increase basal expression but renders *Sox2* entirely independent of NIPBL. Moving the SCR closer to the *Sox2* TSS does not increase basal expression. Yet, the mild downregulation observed when NIPBL is depleted is not observed with these proximal SCR reinsertions, demonstrating that the locus is usually slightly dependent on loop extrusion due to the large distance separating the SCR and *Sox2* promoter. Furthermore, proximal insertion of the *Car2* enhancer does not increase basal expression either. This suggests that the consistent basal expression observed with both enhancers, regardless of their position, reflects an inherent property of the locus rather than being limited by maximum transcriptional output (n ≥ 3 flow and 2 clones). This indicates that buffering of basal expression, in spite of bringing the enhancer closer than 100kb, is an inherent attribute of the *Sox2* locus and not due to saturating the output of the promoter (n ≥ 3 flow and 2 clones).

**Figure S22.**
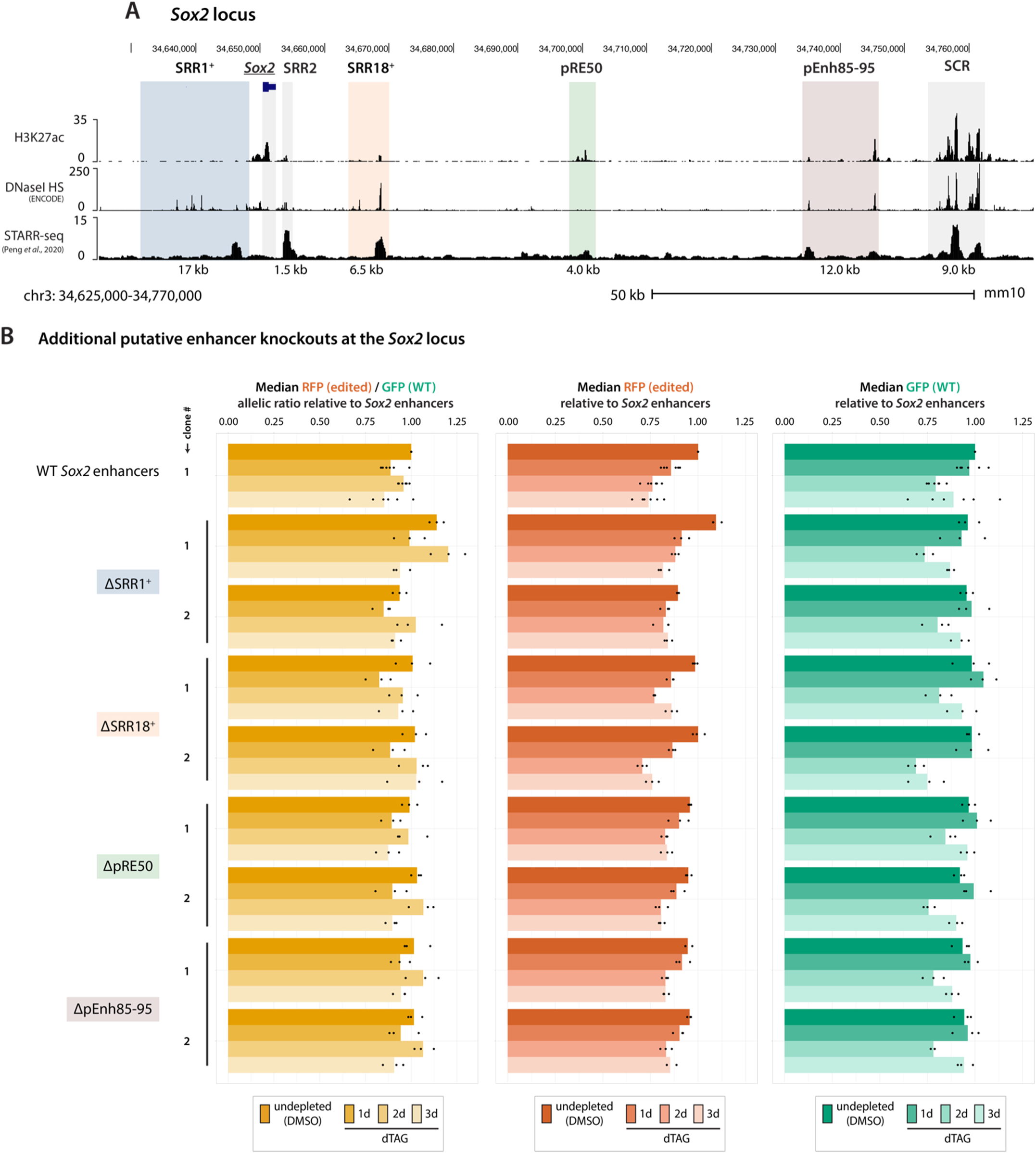
Deletions of additional candidate regulatory elements at the *Sox2* locus have no effect on expression. **(A)** H3K27ac CUT&Tag, DNaseI hypersensitivity (ENCODE), and STARR-seq (*42*) at the *Sox2* locus, highlighting candidate regulatory elements between *Sox2* and the SCR. Gray shading marks the *Sox2* gene, SCR, and SRR2. Colored shading indicates the previously studied SRR1 (dark blue), SRR18 (orange), and pEnh85-pEnh95 (mauve) elements (*32*, *33*). We identified one additional H3K27ac cluster, pRE50 (green). A “+” sign indicates that the deletions are slightly larger than the previously annotated elements (*32*). **(B)** Deletion of the remaining candidate regulatory elements had no effect on either baseline expression or NIPBL-dependence (n ≥ 3 flows and 2 clones).

**Figure S23.**
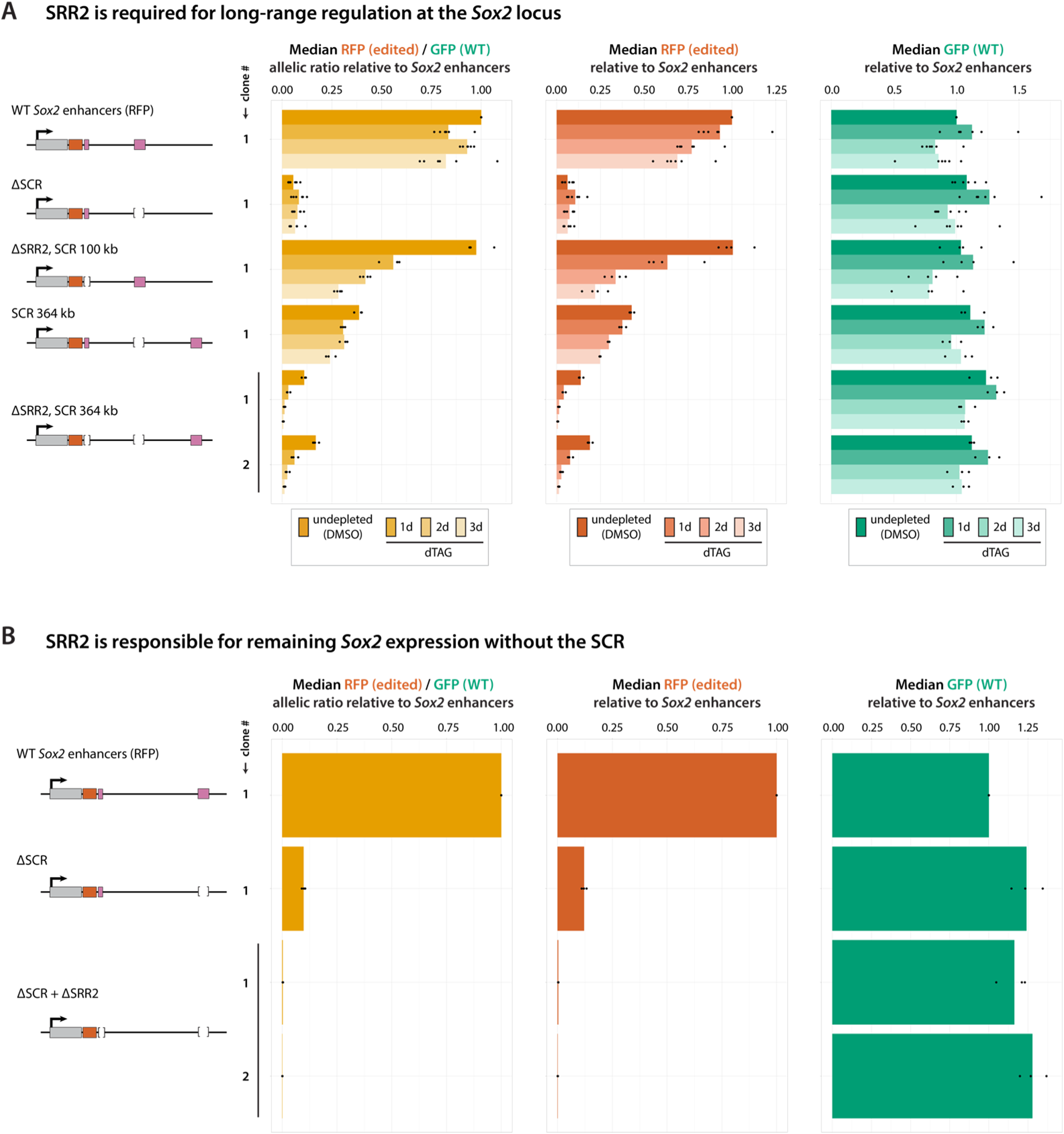
SRR2 is required for long-range regulation at the *Sox2* locus and accounts for residual expression after SCR deletion. **(A)** Deleting SRR2 when the SCR is in its endogenous position (∼100 kb) does not affect basal expression but renders *Sox2* sensitive to NIPBL depletion. Deleting SRR2 when the SCR has been moved much further (364 kb) decreases both basal expression and increases cohesin dependence, with expression becoming nearly undetectable after 3 days of NIPBL depletion (n ≥ 3 flows and 2 clones). **(B)** *Sox2* expression decreases to ∼10% after deletion of the SCR. Additional deletion of SRR2 on the same allele reduces expression to nearly undetectable levels, indicating that SRR2 can act as a weak proximal enhancer of *Sox2* (n ≥ 3 flows and 2 clones).

**Figure S24.**
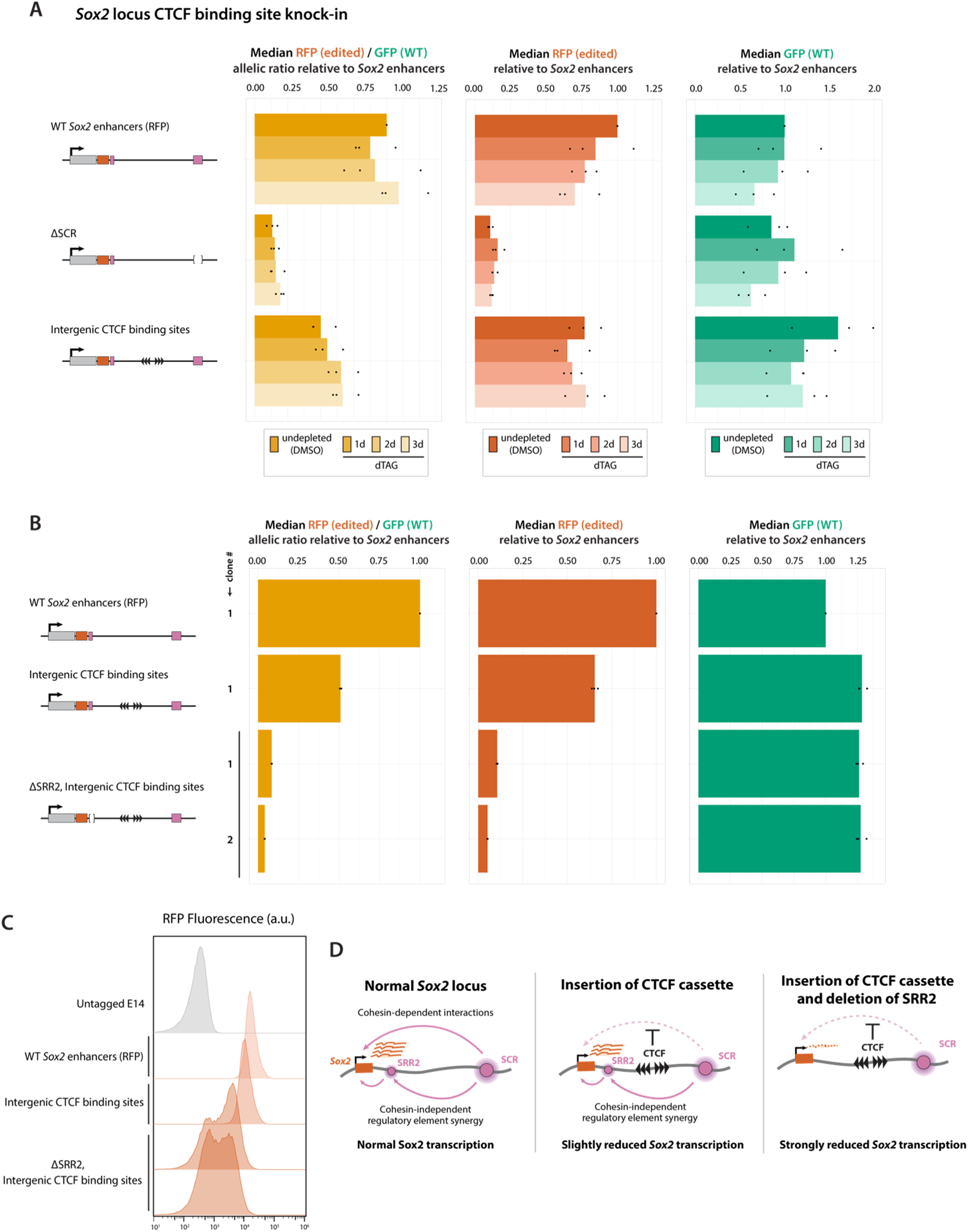
SRR2 can enable the bypass of a strong CTCF insulator at *Sox2*. **(A)** Inserting strong CTCF binding sites (3X reverse, 3X forward) between *Sox2* and the SCR only decreases expression ∼25% (RFP allele) and does not affect the sensitivity to NIPBL depletion (n ≥ 3 flows). **(B)** Deleting SRR2 on the same allele leads to a sharp reduction in *Sox2* expression, suggesting that SRR2 normally facilitates the bypass of the strong boundary and maintains communication between *Sox2* and the SCR (n ≥ 3 flows and 2 clones). **(C)** Representative flow histograms from B. **(D)** Cartoon explaining how CTCF insulators can be partially bypassed at *Sox2*. Intervening CTCF sites block cohesin from extruding across the locus but do not impede the cohesin-independent synergy between the SCR and SRR2. In the absence of SRR2, cohesin is the sole long-range regulatory mechanism; therefore, intervening CTCF sites insulate more effectively, similar to the situation at *Car2* where no SRR2-like element naturally exists.

**Figure S25.**
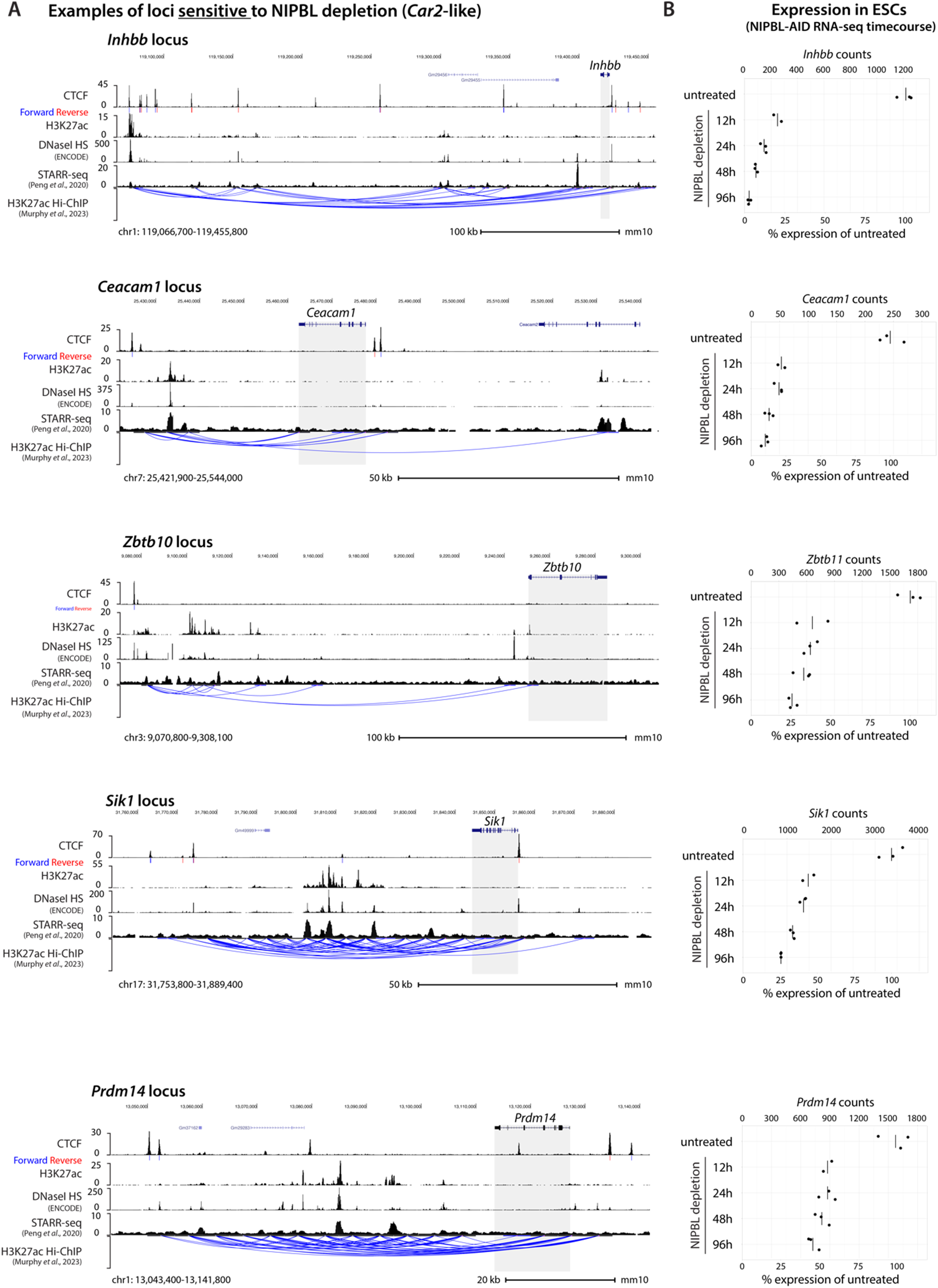
Examples of loci sensitive to NIPBL depletion (*Car2*-like). **(A)** CTCF ChIP-seq with motif orientations, H3K27ac CUT&Tag, DNaseI hypersensitivity (ENCODE), STARR-seq (*42*), and H3K27ac Hi-ChIP (*64*) at various loci that are sensitive to NIPBL depletion and may behave like *Car2*. Gray shading highlights the candidate gene. Loci were chosen based on the presence of at least one putative enhancer ≥ 20 kb from the TSS without candidate proximal regulatory elements, progressive downregulation upon depletion in the NIPBL-AID timecourse, and (not shown) downregulation upon RAD21 depletion (*8*). **(B)** Corresponding mRNA expression at additional loci sensitive to NIPBL depletion in NIPBL-AID ESCs throughout a 96h depletion timecourse. Counts are size factor normalized. Each point is an individual replicate and notches indicate averages (n = 3).

**Figure S26.**
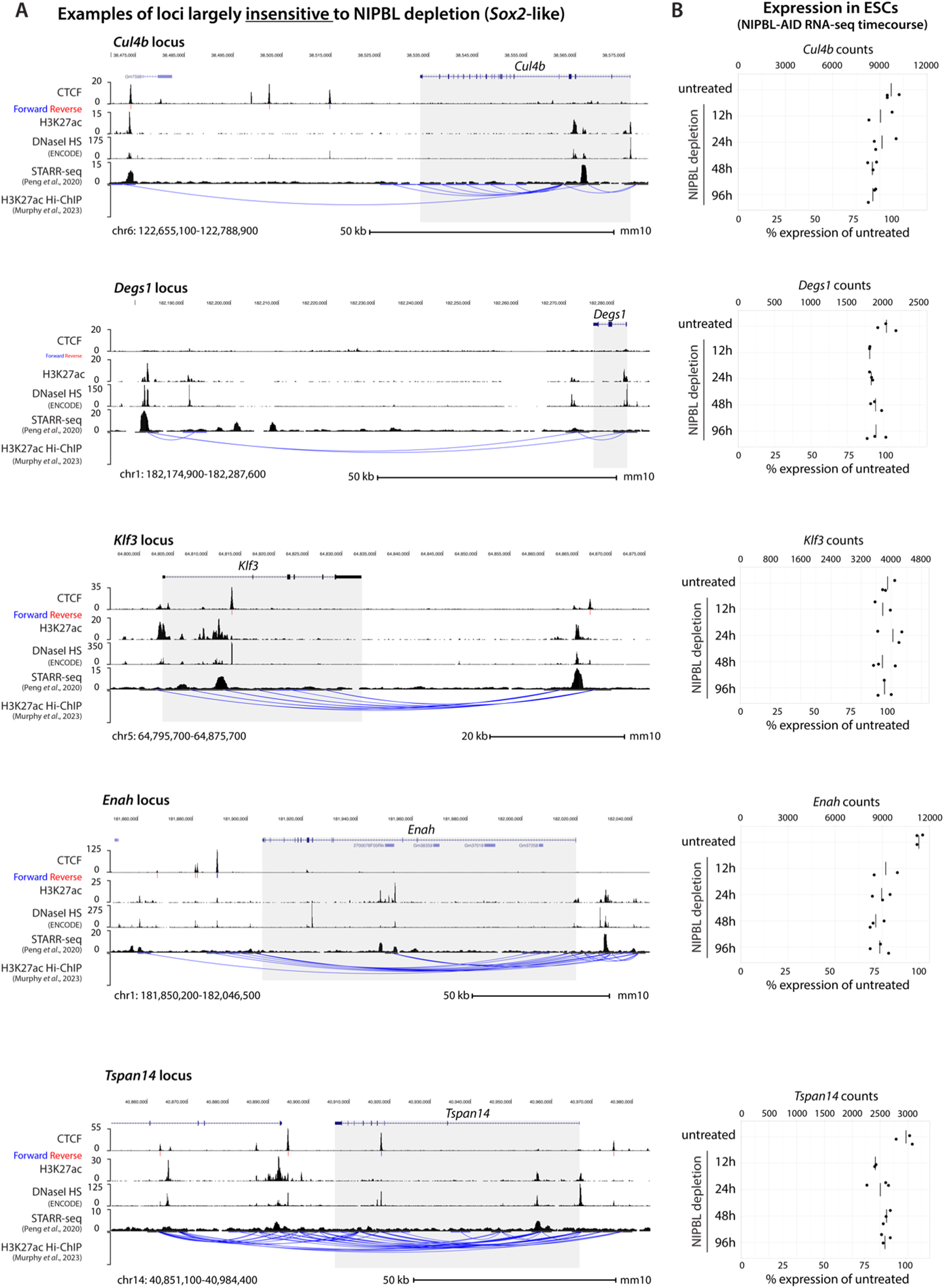
Examples of loci insensitive to NIPBL depletion (*Sox2*-like). **(A)** CTCF ChIP-seq with motif orientations, H3K27ac CUT&Tag, DNaseI hypersensitivity (ENCODE), STARR-seq (*42*), and H3K27ac Hi-ChIP (*64*) at various loci that are largely insensitive to NIPBL depletion and may behave like *Sox2*. Gray shading highlights the candidate gene. Loci were chosen based on the presence of both putative enhancers ≥ 20 kb and additional candidate regulatory elements ≤ 10 kb from the TSS, minimal downregulation throughout depletion in the NIPBL-AID timecourse, and (not shown) no/minimal downregulation upon RAD21 depletion (*8*). **(B)** Corresponding mRNA expression in NIPBL-AID ESCs throughout a 96h depletion timecourse. Counts are size factor normalized. Each point is an individual replicate and notches indicate averages (n = 3).

**Figure S27.**
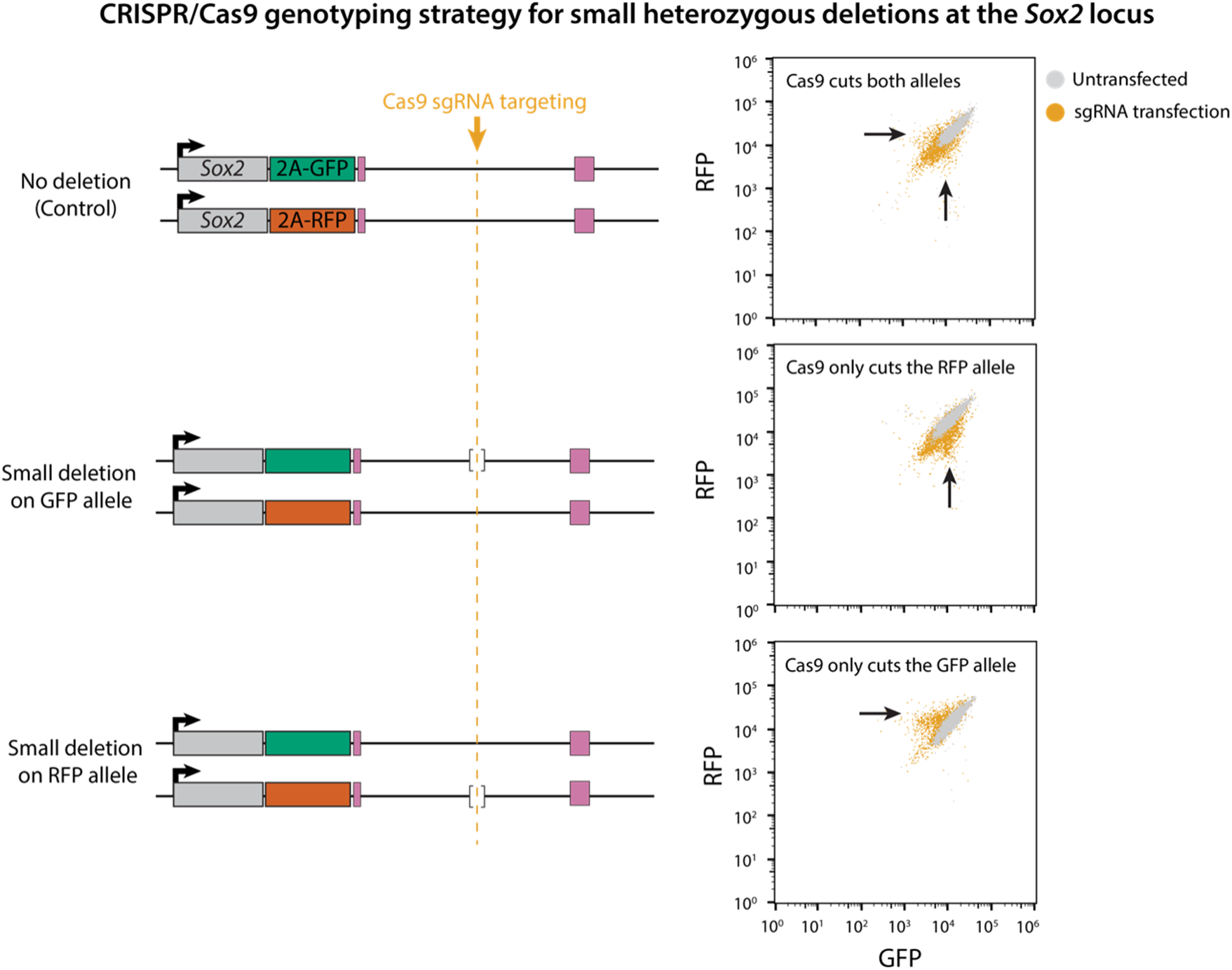
Allelic phasing in *Sox2*-reporter ESCs. Strategy for phasing genomic edits at *Sox2* to the GFP or RFP allele. We noticed temporary downregulation of Sox2 when Cas9 is directed to cut between Sox2 and the SCR, which lasts for 1 to 2 days. Directing Cas9 to cut at the site of the edit, e.g. within a heterozygous deletion or knock-in, therefore enables identifying which allele bears the targeted edit by flow cytometry. Examples of clonal lines with heterozygous deletion assigned to the green or red allele are shown.

